# Genetically informed single-cell and spatial mapping of metabolic programs in human health and disease

**DOI:** 10.64898/2026.06.25.734643

**Authors:** Heng Xu, Guangrui Huang, Linpeng Zhang, Wenbin Liu, Qixing Wu, Minshuo Chen, Dongdong Zhao, Yaqi Zhang, Anlong Xu

## Abstract

Defining cell-type-specific endogenous metabolic features, the spatial distribution of cell-level metabolic states and cellular responses to exogenous metabolites is very important for understanding disease mechanisms. However, existing transcriptome-based metabolic models primarily infer intracellular reaction or pathway-level activities, and therefore cannot directly assess associations between individual metabolite levels and cellular states, particularly for metabolites that act extracellularly as signalling molecules rather than entering cells as metabolic substrates. To overcome this problem, we introduce the gmMAP (Genetically informed metabolite trait mapping across single-cell and spatial tissues), a framework that integrates metabolite GWAS summary statistics with single-cell and spatial transcriptomes to map metabolic programmes at cellular and spatial resolution. Notably, the gmMAP enables the prediction of endogenous metabolic process activation while also revealing intrinsic associations between exogenous metabolites and diverse cellular functional states. To further capture the connectivity of cellular metabolic networks, we incorporated a constraint-based metabolic flux model to evaluate global metabolic activity. To evaluate the accuracy and generalizability of gmMAP, we applied the framework across representative biological contexts spanning human development, physiological homeostasis, inflammation and cancer. In human kidney development, the gmMAP captured dynamic metabolic programmes, which was validated using paired transcriptomic and metabolomic reference datasets, supporting its reliability in metabolite identification and metabolic-flow inference. At the organ level, the gmMAP reconstructed spatial metabolite distribution patterns across 17 mouse organs under homeostatic and autoimmune inflammatory conditions, and further extension of gmMAP to 24 normal human tissues generated a multi-scale metabolic atlas at both organ and cellular resolutions. In disease settings, gmMAP revealed metabolic reprogramming across 29 pan-cancer cell populations, and identified potential links between exogenous metabolites and inflammation-associated stromal metabolic remodelling in ulcerative colitis. Together, gmMAP can consistently connect genetically informed metabolite traits with cell states, spatial tissue organization and disease pathology.

## Introduction

The rapid advancement of single-cell and spatial transcriptomics has revolutionized the dissection of cellular heterogeneity and spatially resolved gene expression landscapes across diverse tissues, enabling unprecedented insights into embryonic development, physiological homeostasis and disease pathogenesis [1–4]. In parallel, large-scale human metabolomic studies have advanced our understanding of metabolic dysregulation in disease contexts. For instance, recent population-scale plasma metabolome studies have linked hundreds to thousands of circulating metabolic traits with human diseases, clinical phenotypes and genetic architecture, establishing metabolomics as a powerful layer for understanding disease biology and therapeutic opportunities [5,6]. Nevertheless, such population-level investigations remain largely confined to bulk circulating metabolite profiling and fail to resolve tissue-specific and celltype-specific metabolic alterations underlying disease pathogenesis. Despite technological breakthroughs, direct and high-resolution characterization of single-cell metabolic patterns and metabolic flux remains a major technical bottleneck. Emerging single-cell and spatial metabolomics technologies, including imaging mass spectrometry- and mass-spectrometry-based single-cell approaches, have begun to resolve metabolite heterogeneity in tissue contexts [7–9]. However, these approaches remain limited by cost, throughput, metabolite coverage, sensitivity, molecular annotation and the difficulty of directly tracing pathway-level metabolic flux at large scale [7–10]. Consequently, robust and cost-effective strategies for systematic single-cell metabolic annotation across tissue spatial contexts are still lacking.

Recent years have witnessed the rapid development of single-cell metabolic modelling tools, represented by Compass and scFEA, which enable the inference of cell-specific metabolic states and flux-like reaction activities from single-cell transcriptomic profiles [11,12]. These approaches have provided important frameworks for reconstructing intracellular metabolic reactions and dissecting cellular metabolic heterogeneity. However, their inference remains subject to several inherent limitations. First, these methods largely depend on curated metabolic networks and enzyme–reaction annotations, thereby restricting their analytical scope to well-characterized endogenous metabolic pathways [11,12]. Second, they rely on the assumption that the transcriptional abundance of metabolic enzymes can approximate reaction activity, an assumption that is inevitably affected by the sparsity, technical noise and limited gene detection of single-cell RNA-seq data [13,14]. More broadly, mRNA abundance does not fully capture protein abundance, enzyme activity, substrate availability, allosteric regulation, post-translational modification or subcellular compartmentalization, all of which shape true metabolic flux [15,16]. As a result, transcriptome-derived flux estimates should be interpreted as inferred metabolic activity or flux-like potential rather than directly measured biochemical flux. In addition, existing frameworks have limited capacity to resolve spatially organized metabolic abnormalities at tissue and cellular resolution and are not well suited to modelling cellular responses to exogenous or exposure-related metabolites, including hormones, gut microbiota-derived metabolites, xenobiotics, dietary metabolites and drug-associated metabolic features [17–19].

Against this backdrop, gmMAP (Genetically informed metabolite trait mapping across single-cell and spatial tissues) provides a complementary analytical strategy for single-cell and spatial metabolic inference. Rather than relying solely on transcriptome-derived enzyme activity, gmMAP integrates metabolite GWAS-derived genetic signatures with single-cell and spatial transcriptomic atlases to infer cell-level metabolite-associated programmes across a broad spectrum of metabolic traits. In this study, metabolite GWAS summary statistics were mapped to gene-level association scores using MAGMA, a generalized gene- and gene-set analysis framework for GWAS data [20]. By leveraging genetically anchored metabolite gene programmes instead of depending exclusively on the expression of individual metabolic enzymes, gmMAP reduces the impact of single-cell transcriptomic sparsity and technical noise on metabolite-trait inference. This design enables the systematic inference of metabolite-associated features at cellular resolution, including endogenous metabolites, metabolite ratios, microbiome-derived metabolites, hormones and other exposure-associated metabolic traits, thereby extending single-cell metabolic analysis beyond canonical intracellular reaction networks. Furthermore, gmMAP incorporates multi-dimensional biological priors, including metabolic pathway topology, reaction order, enzyme annotations and subcellular compartmental constraints, to derive pathway-level metabolic-flow activities across cells, tissues and developmental trajectories.

Leveraging this framework, we systematically resolved spatiotemporal metabolic landscapes across physiological, developmental and disease contexts. gmMAP first enabled high-specificity prediction of spatially resolved metabolite distribution patterns and reconstructed whole-organ spatial metabolic features across 17 mouse organs under homeostatic and inflammatory states. In renal development, gmMAP delineated dynamic metabolic programmes governing cell differentiation, and paired transcriptomic and metabolomic reference datasets validated its accuracy in metabolite identification and relative metabolic-flow inference. We further extended gmMAP to 24 normal human tissues, establishing a genetically informed pan-tissue metabolic atlas at cellular resolution. In disease settings, gmMAP revealed extensive metabolic rewiring across 29 pan-cancer cell populations and decoded inflammation-associated stromal metabolic remodelling in ulcerative colitis, uncovering potential roles of exogenous metabolites in colonic fibroblast activation during colitis pathogenesis. Together with its capacity to predict cellular developmental perturbations and metabolite-targeted drug effects, gmMAP provides a versatile framework for connecting metabolic programmes with tissue physiology, disease pathology and therapeutic intervention.

## Result

### Overview of gmMAP

The gmMAP framework establishes a genetically informed strategy for single-cell and spatial metabolite mapping by integrating metabolite GWAS-derived gene-level association signals with cell-resolved transcriptomic profiles. This framework was designed to infer associations between metabolites and cells, tissues or spatially organized anatomical regions, thereby enabling systematic characterization of cell-type-specific and spatially resolved metabolic programs. The input data of gmMAP include metabolite GWAS summary statistics, or the metabolic trait–gene Z-score matrix generated by GWAS/MAGMA analysis in this study, single-cell transcriptomic data, spatial transcriptomic data and cell-type annotations (Fig. 1a). gmMAP uses the MAGMA framework [20] to convert GWAS summary statistics from 1,091 metabolite levels and 309 metabolite ratios (Supplementary Fig S1 a-d), including endogenous metabolites, metabolite ratios and exogenous or gut microbiota-derived metabolites, into a gene-level metabolite association matrix [6]. Benchmarking analyses indicated that a 1,000-gene signature provided a robust representation of each metabolite trait and improved the prediction of metabolite-associated cellular dynamics (Fig 1b, Supplementary Fig S2).

**Fig 1.**
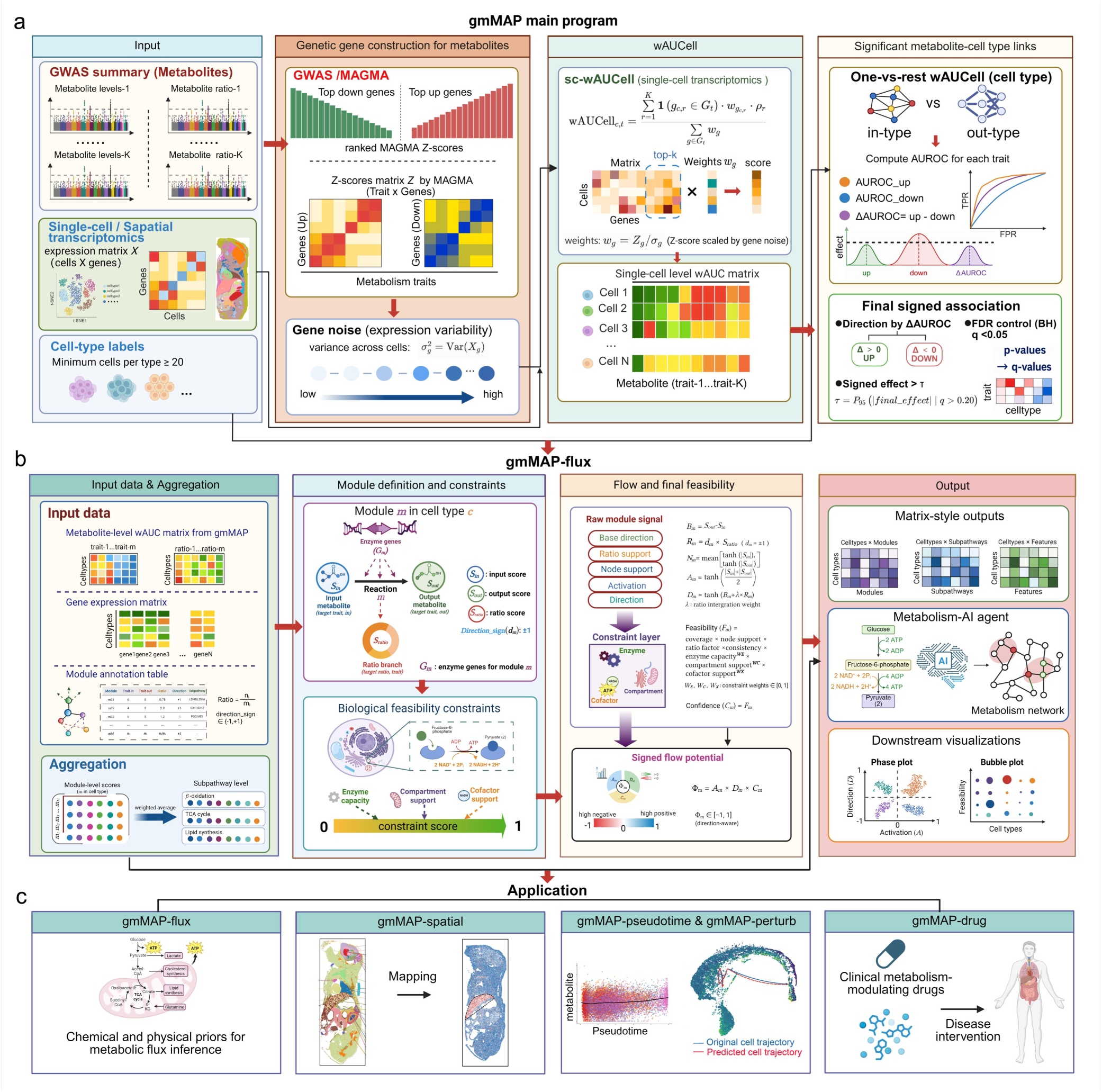
The gmMAP framework for genetically informed single-cell and spatial metabolite mapping. **a.** Schematic overview of the gmMAP workflow. gmMAP integrates metabolite GWAS summary statistics, single-cell or spatial transcriptomic expression matrices, and cell-type annotations to infer metabolite–cell-type associations at cellular or spatial resolution. Metabolite GWAS signals, including metabolite levels and metabolite ratios, are mapped to gene-level association scores using GWAS/MAGMA analysis, generating a trait-by-gene Z-score matrix. For each metabolite trait, genes are ranked according to their genetic association Z-scores to define positively and negatively associated gene programs. Gene-expression variability across cells or spatial observations is further estimated as gene noise and incorporated into gene weighting. gmMAP then applies a weighted rank-based enrichment framework, termed sc-wAUCell, to calculate metabolite-linked enrichment scores for individual cells or spatial spots. Cell-type specificity is evaluated using a one-versus-rest AUROC framework, comparing metabolite scores in a given cell type with all remaining cells. Directional AUROC effects from the up- and down-associated gene programs are integrated to define the final signed metabolite–cell-type association, with the direction determined by ΔAUROC. Statistical significance is controlled using the Benjamini–Hochberg procedure, and robust associations are further filtered using an empirical-null effect-size threshold derived from non-significant background associations. **b.** Enzyme-constrained, ratio-integrated metabolic flow potential model. To estimate the activity and feasibility of cellular metabolic flow, gmMAP-flux integrates metabolite-linked wAUCell scores, gene-expression profiles and curated metabolic module annotations. Input data include metabolite-linked cell-by-trait score matrices, gene-expression matrices and module annotation tables defining metabolic branches, subpathways and reaction directions. Each metabolic module is represented by input metabolites, output metabolites, ratio-linked traits, enzyme genes and directionality information. Module-level signals are aggregated to quantify activation, base direction, ratio-supported directionality and node support. Biological feasibility is further constrained by enzyme capacity, subcellular compartment compatibility and cofactor availability, generating a constraint score that reflects the biochemical plausibility of each module. These components are combined to calculate a signed flow potential score, which captures both the predicted activation and direction of metabolic flow. The resulting outputs include cell type-by-module, cell type-by-subpathway and cell type-by-feature matrices, which can be visualized using heatmaps, constraint landscapes, phase plots and bubble plots to characterize cell-type-specific metabolic network states. Metabolism-AI is a literature-trained, knowledge-graph-guided metabolic reasoning agent built on a tumor-metabolism corpus and dual-model framework, designed to interpret metabolite-associated cellular states, prioritize pathway mechanisms and candidate genes, and support biological validation of tumor metabolic networks. **c.** Downstream applications of gmMAP. gmMAP supports multiple analytical modules for interpreting metabolite-associated cellular programs across biological contexts. gmMAP-flux incorporates chemical, physical and biological prior knowledge to infer pathway-level metabolic flow potential. gmMAP-spatial maps genetically informed metabolite programs onto spatial transcriptomic tissues to resolve spatially organized metabolic landscapes. gmMAP-pseudotime links metabolite-associated scores with developmental or disease-related cellular trajectories, enabling the identification of metabolites associated with cell-state transitions. gmMAP-perturb predicts the impact of metabolite inhibition on cell fate. gmMAP-drug connects disease-associated metabolic programs with clinical metabolism-modulating drug signatures to nominate candidate perturbations that may reverse pathological metabolic states. This figure was created using BioRender.

For each metabolite trait, genes were ranked according to their gene level genetic association Z-scores. The top 1,000 positively associated genes were defined as the up-associated metabolite gene program, whereas the top 1,000 negatively associated genes were defined as the down-associated metabolite gene program. This directional design was used to reduce false positive associations that may arise when broad metabolite-linked genetic signatures are mapped onto sparse single cell transcriptomic profiles. To further account for gene level transcriptional variability, gmMAP estimates expression noise across cells or spatial observations and incorporates this information into gene weighting. Genes with stronger genetic association and lower expression noise are therefore assigned higher weights during metabolite program scoring (Fig. 1a).

For single-cell transcriptomic data and spatial transcriptomic data with single cell or near cellular resolution, gmMAP quantifies metabolite-associated activation and response intensity using a weighted AUCell-like enrichment strategy, referred to as sc-wAUCell. This rank-based scoring approach evaluates genetically informed metabolite gene programs in individual cells or spatial spots while reducing the influence of single-cell transcriptional sparsity and technical noise. For bulk RNA-seq or other dense, non-sparse expression matrices, gmMAP can alternatively use sMRS, a weighted linear aggregation strategy, to estimate sample-level metabolite relevance scores. For spatial transcriptomic datasets composed of mixed spots, gmMAP can be combined with existing deconvolution or spot-embedding approaches to improve the mapping of metabolite-associated programs to spatially localized tissues and cell states (Fig. 1a) [21].

Cell-type-level metabolite associations are quantified using a one-versus-rest AUROC framework. For each cell type and metabolite trait, gmMAP compares metabolite scores in the target cell type against all remaining cells and calculates directional AUROC effects for the up and down associated gene programs, denoted as AUROC-effect-up and AUROC-effect-down, respectively. Directional dominance is summarized as ΔAUROC. The sign and magnitude of ΔAUROC are used to determine the final signed metabolite–cell-type association. A positive ΔAUROC indicates an up-program-dominant association, whereas a negative ΔAUROC indicates a down-program-dominant association. Multiple testing is controlled using the Benjamini–Hochberg procedure, and robust associations are further filtered using an empirical-null effect-size threshold estimated from non-significant background associations (Fig. 1a).

To further estimate the activity and feasibility of cellular metabolic flow, gmMAP incorporates a ratio-integrated and enzyme-constrained metabolic flow potential model (Fig. 1b). This model integrates gmMAP derived metabolite-linked wAUCell score matrices, gene expression profiles and curated metabolic module annotations. Each metabolic module is defined by input metabolites, output metabolites, ratio-linked metabolite traits, enzyme genes and reaction direction information. Module-level signals are aggregated to quantify activation, base direction, ratio-supported directionality and node support. Biological feasibility is further constrained by enzyme capacity, subcellular compartment compatibility and cofactor availability. These constraints are combined to estimate module feasibility and confidence, generating a signed flow potential score that reflects both predicted activation and biological plausibility of metabolic flow. To support mechanistic interpretation of these outputs, we further developed Metabolism-AI, a literature-trained and knowledge-graph-guided metabolic reasoning agent built on a tumor-metabolism corpus and dual-model framework, which links metabolite-associated cellular states to pathway mechanisms, candidate regulatory genes and literature-supported biological processes for downstream biological interpretation and validation prioritization. The resulting outputs include cell type by module, cell type by subpathway and cell type by feature matrices, which can be visualized using heatmaps, constraint landscapes, phase plots and bubble plots to characterize cell type specific metabolic network states.

Based on these metabolite-cell and metabolite-tissue association maps, gmMAP supports five major downstream analytical modules (Fig. 1c). First, gmMAP-flux incorporates reaction order, subcellular compartmentalization, enzyme chemistry and cofactor requirements, together with biological prior knowledge, to infer pathway-level metabolic flow potential. Second, gmMAP-spatial projects genetically informed metabolite programs onto spatial transcriptomic tissues to resolve spatially organized metabolic landscapes. Third, gmMAP-pseudotime links metabolite-associated scores with developmental or disease-related trajectories, enabling the identification of metabolites associated with cell-state transitions. Fourth, gmMAP-perturb integrates metabolite-associated programs with developmental trajectories to predict how inhibition of individual metabolites may reshape cell-state transitions and alter lineage fate bias, enabling the prioritization of metabolites with potential regulatory roles in cell differentiation and disease progression. Fifth, gmMAP-drug compares disease-associated metabolite programs with metabolism-modulating drug signatures to nominate candidate perturbations that may reverse pathological metabolic states.

Compared with existing single-cell metabolism analysis methods, gmMAP provides a distinct advantage by integrating metabolite GWAS-derived genetic information with single-cell and spatial transcriptomic profiles to resolve genetically informed metabolite-associated cellular programmes (Supplementary Table S1). Compass and scFEA are primarily designed to infer reaction- or module-level metabolic activity based on enzyme expression, metabolic reaction networks or predefined metabolic graphs, whereas scMetabolism mainly estimates pathway-level activity. These approaches are therefore less suited to directly capture associations between individual metabolite traits, metabolite ratios, microbial or xenobiotic metabolites, hormones and signalling metabolites with specific cellular states. For example, Compass primarily infers transporter- and enzyme-supported intracellular reaction activity, whereas gmMAP can capture metabolite-associated cell-state programmes driven by extracellular signalling metabolites, such as deoxycholic acid acting through the surface receptor TGR5 to remodel inflammatory fibroblast states without necessarily entering intracellular metabolic flux, this receptor-mediated mode of metabolite action is further illustrated in the ulcerative colitis analysis below. In contrast, gmMAP enables cell-state-resolved mapping of individual metabolite traits, supports spatial metabolite projection, identifies cell fate-associated metabolites, evaluates metabolite-perturbation effects on cell fate, and extends the analysis to metabolite-ratio features. Thus, gmMAP complements rather than replaces flux-oriented methods by bridging genetic metabolomics and single-cell/spatial transcriptomics, providing a metabolite-resolution framework to investigate metabolic remodeling during development, disease progression and cellular perturbation.

Together, gmMAP links metabolite genetics with single-cell and spatial transcriptomic states to infer metabolite-associated cellular programs, spatial metabolic landscapes, developmental metabolic dynamics and drug-reversible disease-associated metabolic states. By integrating genetically anchored metabolite-trait inference with reaction-flow modelling, gmMAP provides a scalable framework for decoding spatiotemporal metabolic organization in development, tissue homeostasis and disease.

### Validation of gmMAP

To benchmark the capacity of gmMAP to infer metabolite activity from single-cell transcriptomes, we assembled a kidney developmental atlas comprising single-cell RNA-seq profiles [22] and a matched spatial metabolomics dataset [23] spanning overlapping renal cell populations. The spatial metabolomics data provided cell-type-resolved measurements of metabolite abundance across developing kidney compartments, thereby offering an orthogonal reference for evaluating transcriptome-based metabolic predictions. We applied gmMAP to the kidney developmental single-cell transcriptomic dataset to infer metabolite-associated activity at single-cell resolution and aggregated these predictions across matched developmental cell types. Comparison with the spatial metabolomics measurements revealed concordant cell-type-specific metabolic patterns, indicating that gmMAP accurately captured metabolite programs associated with renal lineage specification and maturation.

To identify the optimal number of GWAS-linked genes for each metabolite trait in gmMAP, we benchmarked prediction performance using paired kidney developmental datasets. The expected discovery rate peaked when selecting the top 1000 genes and declined thereafter, implying excessive genes weaken cell-type-specific metabolic signals (Fig. S2a). Small gene sets yielded unstable background estimates, which stabilized when gene number reached 500 or above (Fig. S2b). Spearman correlation and Jaccard similarity analyses further confirmed that gene sets of 500–2000 produced consistent metabolic profiles. Combining all metrics, the top 1000 genes per metabolite trait was determined as the optimal default for subsequent gmMAP analyses (Fig. S2c, d).

We first examined the developing kidney cells profiled by spatial metabolomics to define the native metabolic architecture of nephrogenesis. Unsupervised analysis of cell-type-resolved metabolite abundances revealed that metabolite-defined modules clearly separated the major nephrogenic epithelial and stromal compartments, including podocytes, proximal tubular cells, nephron progenitor cells and early epithelial intermediates. These patterns indicate that distinct renal epithelial subpopulations acquire highly divergent metabolic states during development, with specific metabolic programs preferentially allocated to defined developmental lineages. Consistently, closely related cell states, such as PT1 and PT2 cells or RVCSBa and RVCSBb intermediates, exhibited more similar metabolite profiles, supporting a tight coupling between cellular function, developmental identity and metabolic specialization (Fig. 2a, b, c). We next applied gmMAP to single-cell transcriptomic profiles from the developing kidney to infer metabolite-associated activity at single-cell resolution. Remarkably, clustering of gmMAP-predicted metabolite programs across renal cell types reproduced the lineage relationships observed in the spatial metabolomics data (Supplementary Fig. S3a, b). Functionally related cell populations displayed convergent predicted metabolic states, as exemplified by the close clustering of ICa and ICb cells, as well as NPCa, NPCb and NPCc nephron progenitor subsets (Fig. 2d, e, f).

**Fig 2.**
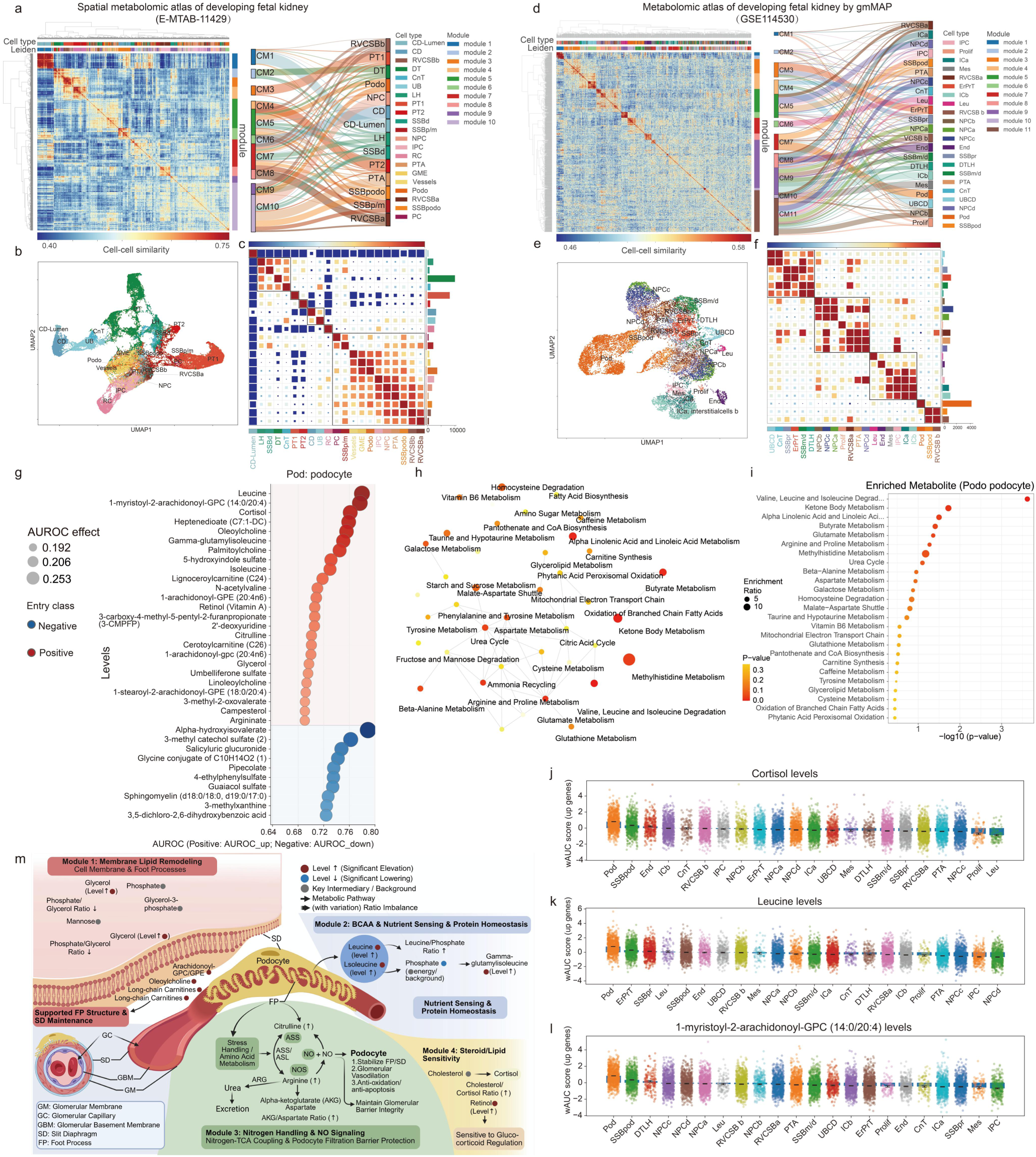
Validation of gmMAP using a developing fetal kidney spatial metabolic atlas and single-cell RNA-seq data. **a.** Heatmap of cell–cell metabolic similarity based on spatially measured metabolite abundance, with annotations for cell type, Leiden clusters and metabolite-defined modules. The accompanying alluvial plot links metabolite modules to renal developmental cell types, showing that metabolite-defined programs separate major nephrogenic and stromal compartments. **b.** UMAP representation of developing kidney cells based on spatial metabolomic profiles, showing segregation of major renal epithelial and stromal populations. **c.** Cell-type similarity matrix summarizing metabolite-based relationships across renal developmental cell states. **d.** Heatmap of cell–cell similarity inferred from gmMAP-predicted metabolite programs, with annotations for cell type, Leiden clusters and predicted metabolite modules. The alluvial plot shows the relationship between gmMAP-defined metabolic modules and renal cell identities. **e.** UMAP of cells clustered by transcriptomic data. **f.** Cell-type similarity matrix based on gmMAP-predicted metabolite programs. **g.** Ranked podocyte-associated metabolite traits for metabolite levels based on gmMAP-predicted metabolite programs. Metabolites were ordered by directional AUROC effects, with positive entries indicating enrichment of up-associated metabolite gene programs and negative entries indicating enrichment of down-associated programs. Bubble size denotes the magnitude of the absolute effect. **h.** Network representation of podocyte-enriched metabolic pathways based on gmMAP-predicted metabolite programs, highlighting coordinated modules related to amino-acid metabolism, branched-chain amino-acid metabolism, steroid-related metabolism, glycerophospholipid metabolism, sphingolipid metabolism, fatty-acid metabolism and mitochondrial fuel handling. Node size indicates enrichment ratio and node colour indicates statistical significance. **i.** Pathway enrichment analysis of podocyte-associated metabolites. **j–l.** Single-cell distribution of representative gmMAP-predicted podocyte-associated metabolites across developing kidney cell types. j, Cortisol levels, k, leucine levels and l, 1-myristoyl-2-arachidonoyl-GPC levels showed cell-type-biased activity rather than uniform pan-renal distribution, supporting the specificity of gmMAP-inferred metabolite programs. **m.** Schematic summary of the podocyte-centred metabolic program inferred by gmMAP. Four major functional axes were identified: membrane lipid remodelling and foot-process maintenance, branched-chain amino-acid nutrient sensing and protein homeostasis, nitrogen handling and nitric oxide-related signalling, and steroid/lipid sensitivity. **Abbreviations:** NPC, nephron progenitor cell; IPC, interstitial progenitor cell; UB, ureteric bud; CD, collecting duct cell; CnT, connecting tubule; DTLH, distal tubule/loop of Henle; PT, proximal tubule; Podo, podocyte; PTA, pre-tubular aggregate cell; RVCSBa, renal vesicle/comma-shaped body cell a; RVCSBb, renal vesicle/comma-shaped body cell b; SSBm/d, S-shaped body cell medial/distal; SSBpro, S-shaped body cell proximal; SSBpodo, S-shaped body cell podocytes; IC, interstitial cell; End, endothelial cell; ICa, interstitial cell a; ICb, interstitial cell b; Leu, leukocyte; Mes, mesangial cell; Prolif, proliferating cell; TCA, the tricarboxylic acid cycle; ETC, respiratory electron transport chain.

### gmMAP accurately identifies podocyte-centred metabolic specialization during kidney development

Having established that gmMAP recapitulated the global metabolic organization of developing renal cell types, we next examined whether it could nominate biologically interpretable metabolite features for individual lineages. We focused first on podocytes, a highly specialized glomerular epithelial population whose function depends on a complex actin cytoskeleton, interdigitating foot processes and a lipid-rich slit diaphragm[24,25]. gmMAP ranked podocyte-associated metabolite traits by directional AUROC effects and identified a coherent metabolic signature dominated by branched-chain amino acids, steroid-related metabolites, glycerophospholipid species, acylcarnitine-associated metabolites and sphingolipid-related pathways (Fig. 2g–i and Supplementary Fig. S3c). Notably, leucine was among the strongest podocyte-associated metabolites. This prediction is consistent with the known requirement for amino-acid transport and nutrient-sensitive protein homeostasis in podocytes, as well as recent evidence that branched-chain amino-acid catabolic defects directly remodel podocyte metabolism and promote podocyte injury in diabetic kidney disease [26–28]. Thus, gmMAP recovered an amino-acid/nutrient-sensing axis that is highly relevant to podocyte development and maintenance.

Another key characteristic of the gmMAP podocyte signature is cortisol and steroid-related metabolism. Biologically, glucocorticoids serve as systemic immunosuppressants and directly target podocytes via glucocorticoid receptors [29,30]. Evidence indicates they stabilize the podocyte actin cytoskeleton, facilitate injury repair and mitigate podocyte damage linked to proteinuria [31]. The detection of cortisol as a podocyte-associated metabolite thus validates gmMAP’s capacity to predict functionally significant steroid-responsive states.

The lipid component of the podocyte prediction was consistent with established podocyte biology. gmMAP prioritized glycerophosphocholine-related and acylcarnitine-associated metabolites, and pathway enrichment highlighted glycerophospholipid, sphingolipid, fatty-acid and mitochondrial energy-associated metabolism (Fig. 2h, i). Podocytes are particularly dependent on membrane lipid organization because slit diaphragm proteins are embedded in cholesterol- and sphingolipid-enriched membrane microdomains, and perturbation of podocyte lipid or sphingolipid metabolism disrupts foot-process architecture and filtration barrier integrity [32,33]. The enrichment of acylcarnitine and fatty-acid-related signals further suggests that gmMAP captures mitochondrial fuel-handling programs that support the energetic and structural requirements of podocyte maturation.

Cortisol, leucine and 1-myristoyl-2-arachidonoyl-GPC showed cell-type-biased activity patterns, with enriched signals in podocyte-related compartments rather than a diffuse pan-cellular distribution (Fig. 2j–l). Integrating the ranked metabolites and enriched pathways, we summarized the podocyte program into four functional axes: membrane lipid remodelling and foot-process maintenance, branched-chain amino-acid nutrient sensing and protein homeostasis, nitrogen handling and nitric oxide-related signalling, and steroid/lipid sensitivity (Fig. 2m), these axes align closely with known mechanisms of podocyte structure.

Beyond podocytes, gmMAP also resolved lineage-specific metabolic programs across other developing nephron compartments. Early proximal tubular cells were enriched for lipid, carnitine, acylcarnitine and amino-acid-associated metabolites, matching the high mitochondrial and fatty-acid oxidative capacity required for solute transport and reabsorptive function in proximal tubules (Supplementary Fig. S3d, g). [34–36]. Pretubular aggregate cells showed stronger glycine/serine, methionine/homocysteine, phosphatidylethanolamine and polyamine-related programs, consistent with biosynthetic and one-carbon metabolic demands during epithelial commitment (Supplementary Fig. S3f, h) [23]. Distal tubule/loop-of-Henle cells displayed a distinct profile involving taurine/hypotaurine, methionine, mitochondrial and lipid metabolic pathways, suggesting antioxidant and mitochondrial adaptation during tubular maturation (Supplementary Fig. S3e, i) [37,38].

### Flux-potential analysis via gmMAP-flux identifies metabolic remodeling in differentiating proximal tubules

Previous studies have demonstrated that during proximal tubular (PT) differentiation, human fetal kidney cells undergo a prominent metabolic shift from glycolysis to mitochondrial oxidative metabolism and fatty acid β-oxidation [23,39]. Spatial dynamic metabolomics combined with ¹³C-isotope tracing revealed that as cells develop from the RV/CSB and S-shaped body stages into mature proximal tubules, the expression of glycolytic genes decreases while gluconeogenic pathways are enhanced, alongside downregulation of the glycolytic rate-limiting enzyme PKM. Meanwhile, pathway analysis indicated gradual activation of the TCA/ETC and mitochondrial FAO pathways. Tracer experiments using U-¹³C₁₈-linoleate showed increased fatty acid-derived carbon contribution to citrate during proximal tubule maturation, consistent with upregulated expression of the FAO-related enzyme ACAA2 [23,39]. Together, previously reported spatial isotope-tracing metabolomics supports a metabolic switch toward enhanced FAO, acetyl-CoA input, citrate entry and TCA/ETC activity, accompanied by reduced lactate output and glutamine contribution. This schematic was used as an external reference for interpreting PT-associated metabolic remodeling (Fig. 3m).

**Fig 3.**
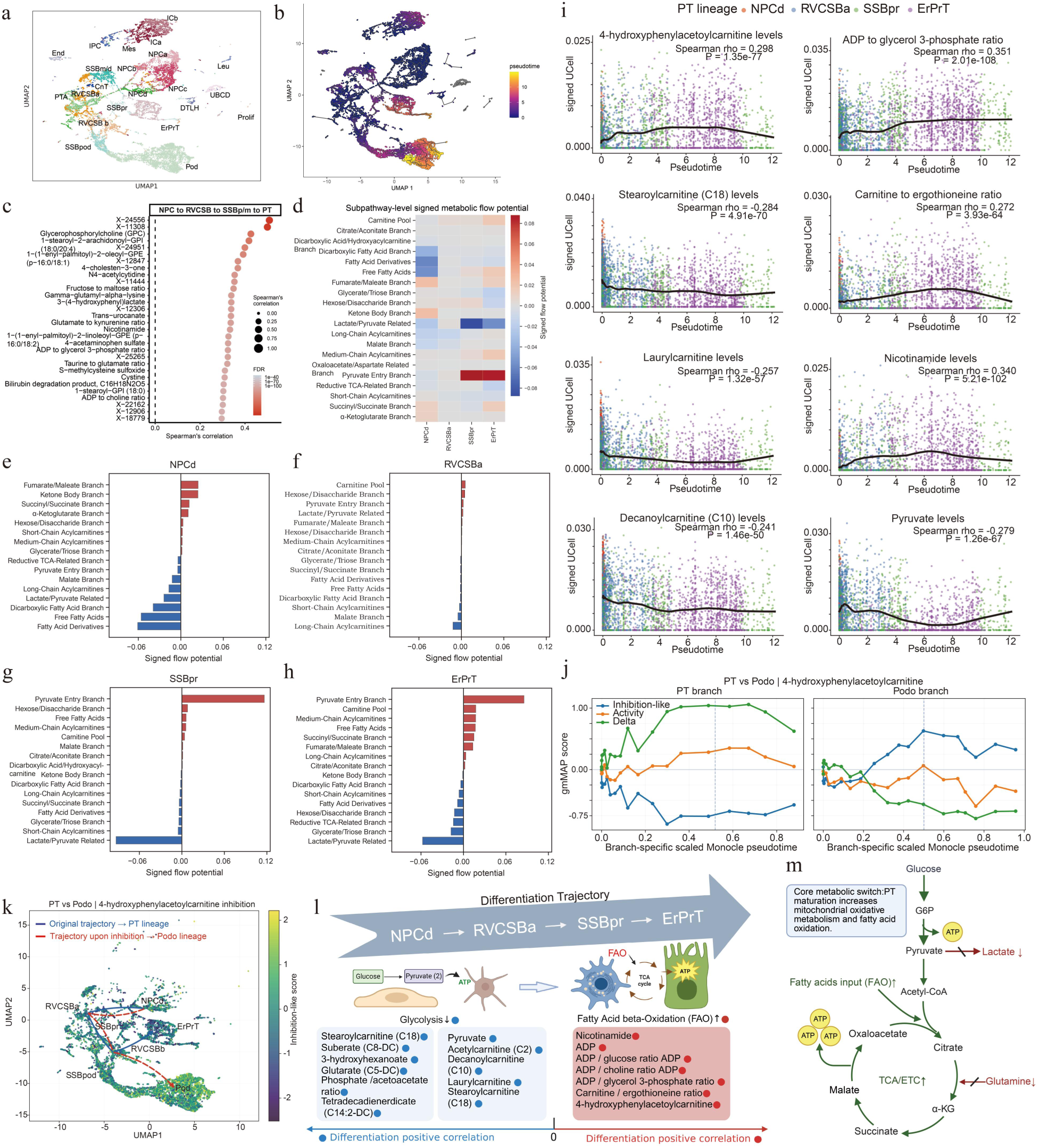
gmMAP-flux predicted metabolic dynamics and flux potential remodeling during proximal tubule differentiation. **a.** UMAP embedding of the kidney developmental dataset annotated by cell type, highlighting the proximal tubule lineage. **b.** Monocle pseudotime projected onto the UMAP, showing progressive ordering of PT-lineage cells from progenitor to differentiated states. **c.** Top positively correlated metabolites and metabolite ratios along the PT lineage pseudotime trajectory (NPCd → RVCSBa → SSBpr → PT), ranked by Spearman correlation. Dot size indicates absolute correlation strength and colour indicates significance. **d.** Heatmap of subpathway-level signed metabolic flow potential across NPCd, RVCSBa, SSBpr and ErPrT for selected glycolysis-, FAO- and TCA-related branches. Red indicates positive flux potential and blue indicates negative flux potential. **e–h.** Ranked branch-level signed metabolic flow potentials for individual PT-lineage states: e, NPCd; f, RVCSBa; g, SSBpr; h, ErPrT. Bars to the right of zero indicate positively activated branches, whereas bars to the left indicate negatively activated branches. **i.** Representative metabolite and metabolite-ratio trajectories along PT pseudotime, including 4-hydroxyphenylacetylcarnitine, ADP-to-glycerol-3-phosphate ratio, stearoylcarnitine (C18), carnitine-to-ergothioneine ratio, laurylcarnitine, nicotinamide, decanoylcarnitine (C10) and pyruvate. Points are coloured by PT-lineage cell state, and the black line denotes the smoothed trend across pseudotime. **j-k.** gmMAP-perturb predicts lineage rerouting associated with 4-hydroxyphenylacetoylcarnitine programs during nephron differentiation. (j) Branch-specific pseudotime dynamics of metabolite-associated scores along proximal tubule (PT) and podocyte (Podo) differentiation trajectories. Activity scores (orange), inhibition-like scores (blue) and integrated Δ scores (green) were calculated from genetically informed metabolite-associated gene programs inferred by gmMAP. The vertical dashed line indicates the approximate branch bifurcation point. PT-directed cells displayed increasing activity and positive Δ scores accompanied by reduced inhibition-like scores, whereas Podo-directed cells exhibited the opposite trend, resulting in progressively negative Δ scores after lineage divergence. (k) UMAP projection of inhibition-like scores for the 4-hydroxyphenylacetoylcarnitine-associated program. Each point represents a single cell colored according to inhibition-like score. Blue arrows denote the original developmental trajectory from nephron progenitors toward the PT lineage, whereas red dashed arrows indicate the direction of fate bias predicted by gmMAP-perturb under suppression of the metabolite-associated program. Cells within the podocyte branch exhibited higher inhibition-like scores than PT-lineage cells, supporting a predicted shift from PT-associated states toward podocyte-associated states upon metabolite-program inhibition. **l.** Schematic summary of metabolic remodeling during PT differentiation. Early progenitor-like states are characterized by a more glycolysis-associated program, whereas SSBpr/ErPrT cells acquire increased mitochondrial substrate entry, FAO-associated metabolism, TCA-cycle activity, oxidative phosphorylation, antioxidant buffering and membrane remodeling. Figure l was created using BioRender. **m.** Conceptual schematic illustrating the metabolic flux remodeling associated with proximal tubule maturation. PT maturation is characterized by increased mitochondrial fatty acid oxidation, enhanced acetate/fatty-acid-derived acetyl-CoA input, elevated citrate entry into the TCA cycle, and increased TCA/ETC activity, consistent with enhanced oxidative energy metabolism. By contrast, lactate production and glutamine-derived contribution to the TCA/glutamate branch are reduced. Green arrows denote increased fluxes or metabolic inputs during PT maturation, and red blocked arrows denote decreased metabolic routes. Figure m was created using BioRender. **Abbreviations:** CnT, connecting tubule; DTLH, distal tubule/loop of Henle; End, endothelial cells; ErPrT, early proximal tubule; ICb, interstitial cells b; ICa, interstitial cells a; IPC, interstitial progenitor cell; Leu, leukocyte; Mes, mesangial cells; NPCa, nephron progenitor cells a; NPCb, nephron progenitor cells b; NPCc, nephron progenitor cells c; NPCd, nephron progenitor cells d; PTA, pretubular aggregate; Pod, podocyte; Prolif, proliferating cells; RVCSB b, renal vesicle/comma-shaped body b; RVCSBa, renal vesicle/comma-shaped body a; SSBm/d, s-shaped body medial/distal; SSBpod, s-shaped body podocyte precursor cells; SSBpr, s-shaped body proximal precursor cells; UBCD, ureteric bud/collecting duct.

Herein, we applied gmMAP to accurately predict the dynamic shifts of metabolic profiles throughout proximal tubular differentiation. To define metabolic remodeling along PT differentiation, we focused on the PT lineage spanning nephron progenitor cells (NPCd), renal vesicle/comma-shaped body cells (RVCSBa), S-shaped body proximal precursor cells (SSBpr) and early proximal tubule cells (ErPrT). UMAP projection of the lineage and Monocle-based pseudotime ordering resolved a continuous developmental trajectory from progenitor to differentiated proximal tubular states, accompanied by a progressive decrease in developmental potency and emergence of the ErPrT state (Fig 3 a,b). Correlation analysis of predicted metabolite levels along pseudotime identified a set of positively PT-lineage-associated metabolites and metabolite ratios, many of which were linked to energy charge, redox state and lipid-derived intermediates (Fig 3c). Representative trajectories further showed increasing 4-hydroxyphenylacetylcarnitine, ADP-to-glycerol-3-phosphate ratio, carnitine-to-ergothioneine ratio and nicotinamide, whereas stearoylcarnitine (C18), laurylcarnitine, decanoylcarnitine (C10) and pyruvate decreased along the PT trajectory (Fig 3i). Together, these trends suggest progressive remodeling of mitochondrial cofactor demand, energy metabolism and fatty-acid-derived substrate utilization during proximal tubule maturation.

To infer metabolic directionality at the pathway scale, we next applied the flux-potential model (gmMAP-flux) to branches associated with glycolysis, fatty acid oxidation, and the TCA cycle across identical developmental states. Subpathway-level signed metabolic flow potential revealed a marked stage-specific transition (Fig 3d). Early NPCd and RVCSBa cells displayed weak or negative signals across many fatty-acid/carnitine-associated branches, whereas SSBpr and especially ErPrT showed strong positive activation of the Pyruvate Entry Branch, together with increased Carnitine Pool, Medium-Chain Acylcarnitines, Fumarate/Maleate and Succinyl/Succinate branches (Fig 3d,h). In contrast, glycolysis-associated branches, particularly Lactate/Pyruvate Related and Glycerate/Triose Branch, were reduced in ErPrT, while hexose/disaccharide metabolism remained weak or near neutral (Fig 3d,h). Cell-state-specific branch ranking further highlighted that NPCd was dominated by negative fatty-acid-related branches, RVCSBa remained metabolically modest, SSBpr represented a transitional state with induction of pyruvate entry and fatty-acid-associated branches, and ErPrT acquired the strongest oxidative metabolic profile (Fig 3e–h).

We next applied gmMAP-perturb to the PT–podocyte bifurcation to determine whether metabolite-associated transcriptional states were linked to nephron fate bias. Among the top-ranked candidates, 4-hydroxyphenylacetoylcarnitine showed a clear branch-specific pattern. Along the PT trajectory, activity scores increased and inhibition-like scores decreased, producing sustained positive Δ scores during late pseudotime. By contrast, podocyte-directed cells progressively accumulated inhibition-like scores and displayed reduced, negative Δ scores after bifurcation (Fig 3j). This reciprocal behavior indicates that the active metabolite-associated program was preferentially aligned with PT differentiation, whereas the inhibition-like program was enriched along the podocyte branch. Consistently, projection of inhibition-like scores onto the developmental manifold showed stronger signals in podocyte-lineage cells than in PT-lineage cells (Fig 3k). gmMAP-perturb therefore predicted that suppression of the 4-hydroxyphenylacetoylcarnitine-associated program would bias cells away from the PT trajectory and toward podocyte-associated states. These findings suggest that aromatic-acylcarnitine-linked transcriptional programs may mark metabolic divergence during PT–podocyte fate specification.These combined analyses support a developmental metabolic switch in the PT lineage, in which immature progenitor-like states progressively transition from a relatively glycolysis-associated program to an oxidative state characterized by enhanced mitochondrial substrate entry, TCA-cycle engagement and FAO-associated metabolic potential (Fig 3l).

### Spatial mapping of tissue-specific metabolite programs by gmMAP-spatial

To validate the capacity of gmMAP for mapping metabolite-linked signatures in structurally intricate tissues, we analyzed whole-body spatial transcriptome slices from untreated and LPS-exposed mice[40]. We performed anatomical annotation on the four slices and delineated major tissue regions, such as the cerebral cortex, caudoputamen, cerebellar cortex, meninges, nerve fiber tracts, periportal and pericentral liver zones, gastric glands, small intestine, renal cortex and medulla, lymphoid tissues, bone marrow and blood vessels (Fig 4a). Mapping metabolite-related gene signatures onto the slices revealed distinct regional distributions, demonstrating that gmMAP recovers authentic tissue metabolic features instead of random background noise (Fig 4b-n).

**Fig. 4.**
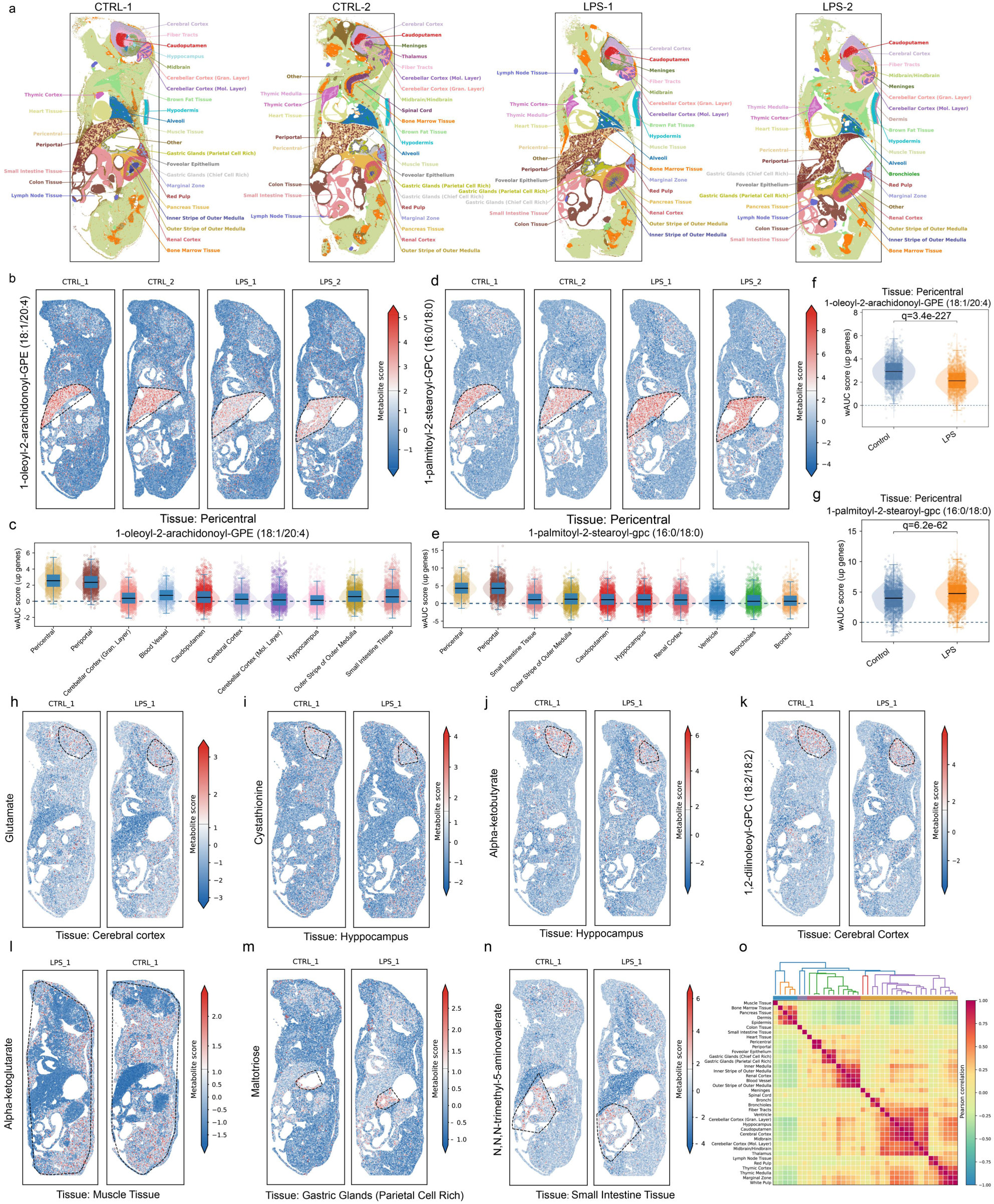
Whole-body spatial mapping of metabolite-associated programs predicted by the gmMAP algorithm in control and LPS-treated mice. **a.** Anatomical annotation of whole-body mouse spatial transcriptomic sections from two control samples and two LPS-treated samples. Major tissue compartments and subregions, including cerebral cortex, caudoputamen, meninges, hippocampus, cerebellar cortex layers, thymic tissue, lymph node tissue, bronchioles, alveoli, brown adipose tissue, liver-associated pericentral and periportal regions, gastric glands, small intestine tissue, colon tissue, pancreatic tissue, renal cortex, inner medulla and outer stripe of outer medulla, were annotated. **b.** Spatial projection of the 1-oleoyl-2-arachidonoyl-GPE (18:1/20:4)-associated gene program across control and LPS-treated sections. Colors indicate metabolite-associated wAUCell scores for the up-associated gene program. Dashed outlines mark the pericentral region. **c.** Tissue-level distribution of 1-oleoyl-2-arachidonoyl-GPE (18:1/20:4)-associated scores across top-ranked anatomical regions. Pericentral tissue showed among the highest scores, supporting spatially zonated enrichment of this arachidonoyl-containing glycerophosphoethanolamine program in the liver. **d.** Spatial projection of the 1-palmitoyl-2-stearoyl-GPC (16:0/18:0)-associated gene program across control and LPS-treated sections. Dashed outlines indicate the pericentral region. **e.** Tissue-level distribution of 1-palmitoyl-2-stearoyl-GPC (16:0/18:0)-associated scores across top-ranked anatomical regions. Pericentral and periportal liver regions showed prominent enrichment, indicating spatially organized phosphatidylcholine-related metabolism. **f.** Comparison of pericentral 1-oleoyl-2-arachidonoyl-GPE (18:1/20:4)-associated scores between control and LPS-treated samples. LPS treatment was associated with a significant reduction of this lipid-associated program. **g.** Comparison of pericentral 1-palmitoyl-2-stearoyl-GPC (16:0/18:0)-associated scores between control and LPS-treated samples. LPS treatment was associated with a significant increase of this phosphatidylcholine-associated program. **h.** Representative spatial map of the glutamate-associated program, highlighting enrichment in the cerebral cortex. **i.** Representative spatial map of the cystathionine-associated program, highlighting enrichment in the hippocampus. **j.** Representative spatial map of the alpha-ketobutyrate-associated program, highlighting enrichment in the hippocampus. **k.** Representative spatial map of the 1,2-dilinoleoyl-GPC (18:2/18:2)-associated program, highlighting enrichment in the cerebral cortex. **l.** Representative spatial map of the alpha-ketoglutarate-associated program, highlighting enrichment in muscle tissue. **m.** Representative spatial map of the maltotriose-associated program, highlighting enrichment in gastric glands enriched for parietal cells. **n.** Representative spatial map of the N,N,N-trimethyl-5-aminovalerate-associated program, highlighting enrichment in small intestine tissue. **o.** Tissue–tissue similarity matrix calculated from metabolite-associated spatial scores. Hierarchical clustering grouped anatomically related tissues and revealed coordinated metabolic similarity among neural, gastrointestinal, renal, immune and epithelial compartments.

Spatially, two distinct glycerophospholipid signatures, namely 1-oleoyl-2-arachidonoyl-GPE (18:1/20:4) and 1-palmitoyl-2-stearoyl-GPC (16:0/18:0), exhibited prominent zone-specific enrichment in the liver pericentral region (Fig 4b, c, d, e). We observed marked phospholipid remodeling in the liver pericentral zone upon LPS-triggered inflammation. Comparative analysis between control and LPS-treated samples revealed opposing expression patterns: 1-oleoyl-2-arachidonoyl-GPE was markedly downregulated, while 1-palmitoyl-2-stearoyl-GPC was significantly upregulated (Fig 4f, g). Our findings are highly consistent with previous lipidomics investigations[41–45]. Cumulative evidence has verified that LPS-induced systemic inflammation triggers pericentral zone-specific phospholipid remodeling in the liver, featuring depletion of arachidonoyl-containing glycerophosphoethanolamines and compensatory upregulation of saturated phosphatidylcholines[41,44]. This divergent metabolic regulation is driven by TLR4-mediated suppression of LPCAT3 and activation of cytosolic phospholipase A2 (cPLA2). The activated cPLA2 mobilizes arachidonic acid from membrane phospholipids and subsequently facilitates the synthesis of pro-inflammatory eicosanoids[42,45]. Mass spectrometry imaging analyses further confirm that the pericentral zone acts as the primary site of hepatic lipid remodeling, where the two lipid signatures show prominent zonal enrichment and inverse responses to LPS stimulation[41,44]. Collectively, these results demonstrate that systemic inflammation drives comprehensive remodeling of hepatic membrane phospholipid metabolism: ethanolamine phospholipids carrying arachidonoyl groups are mobilized and decreased, whereas metabolism associated with saturated phosphatidylcholine is enhanced[43].

gmMAP also identified neuron-related metabolite signatures. Glutamate-associated programs localized to the cerebral cortex, consistent with glutamate being the principal excitatory neurotransmitter in the mammalian brain and mediating most fast excitatory synaptic transmission [46–48] (Fig 4h, Supplementary Fig S4a). Cystathionine was enriched in the hippocampus, suggesting active sulfur amino acid metabolism and redox buffering, in agreement with evidence that the brain transsulfuration pathway contributes to cysteine supply and glutathione-dependent antioxidant capacity [49] (Fig 4i, Supplementary Fig S4b). Perturbations of this metabolic axis, characterized by impaired transsulfuration activity, depleted glutathione, and accumulated homocysteine, are tightly implicated in the pathogenesis of major neurodegenerative disorders including Alzheimer’s and Parkinson’s diseases[50,51]. α-Ketobutyrate, a by-product generated during cystathionine cleavage in the sulfur amino acid transsulfuration pathway, was also highly enriched in the hippocampus, while its abundance decreased markedly following LPS exposure[52,53] (Fig 4j, Supplementary Fig S4c). Consistent with previous work demonstrating that LPS-triggered systemic inflammation profoundly rewires hippocampal amino acid metabolism linked to methionine turnover, neuroinflammation and redox homeostasis, and that this includes an anti-inflammatory-like downregulation of hippocampal methylglyoxal and methionine metabolism[54], the altered level of α-ketobutyrate observed here further confirms that neural sulfur amino acid metabolism undergoes extensive reprogramming in response to inflammatory stress. The 1,2-dilinoleoyl-GPC (18:2/18:2) program was detected in the cerebral cortex and cerebellar granular layer, consistent with regional organization of glycerophosphocholine and PUFA-containing phospholipids in brain membranes [55–57]. Although choline itself exhibited a broad systemic distribution, consistent with its universal role in membrane phospholipid synthesis[58], the 1,2-dilinoleoyl-GPC (18:2/18:2) program showed a spatially restricted, brain-specific pattern, with enrichment in the cerebral cortex, hippocampus, and cerebellar granular layer (Fig 4k, Supplementary Fig S4d, h, i). As a choline-containing phosphatidylcholine, 1,2-dilinoleoyl-GPC functions as a membrane-bound choline reservoir rather than free choline, matching known requirements for neuronal phospholipid synthesis and acetylcholine production[59]. Related studies have also demonstrated cell and region selective enrichment of PUFA-containing phosphatidylcholines in the mouse brain[60]. The coexistence of widespread whole-body choline signals and cortex/hippocampus-enriched 1,2-dilinoleoyl-GPC thus highlights gmMAP’s ability to distinguish systemic choline availability from brain-specific choline-phospholipid metabolism.

Beyond liver and brain, gmMAP identified metabolite programs associated with the motor system and intestinal tissues. α-ketoglutarate showed localized signals in muscle tissue (Fig. 4l, Supplementary Fig S4e), suggesting a broader amino-acid-derived keto acid program shared between selected neural and peripheral metabolic compartments. Maltotriose was enriched in gastric glands enriched for parietal cells, supporting a spatial association with gastrointestinal carbohydrate-related metabolism (Fig. 4m, Supplementary Fig S4f). N,N,N-trimethyl-5-aminovalerate was localized to small intestine tissue, consistent with an intestinal metabolite-associated program potentially linked to diet- and microbiota-related trimethylated metabolites (Fig. 4n, Supplementary Fig S4g). Finally, a tissue–tissue similarity matrix constructed from metabolite-associated spatial scores grouped anatomically and functionally related tissues, including neural regions, gastrointestinal compartments, renal structures and immune-associated tissues. Together, these results demonstrate that gmMAP can reconstruct known spatial metabolic organization, detect inflammation-associated liver phospholipid remodeling, and nominate tissue-specific metabolite programs across neural, hepatic, renal, gastric and intestinal compartments (Fig. 4o).

### gmMAP constructs a human metabolic navigation map

Organ-resolved metabolomic studies have shown that different organs harbour distinct metabolic compositions, including divergent metabolite profiles among liver, kidney and skeletal muscle [61,62]. Emerging single-cell spatial metabolomics has begun to connect metabolites with cell-type identity within selected tissues [8]. However, owing to the high cost, limited throughput and technical complexity of single-cell-resolution spatial metabolomic profiling, its broad application to large-scale whole-body organ atlases remains challenging [63]. Consequently, a body-wide, cell-type-resolved understanding of shared and divergent metabolite-associated programmes across healthy organs remains limited. To address this gap, we compiled cross-tissue atlases of healthy cells and applied gmMAP to dissect metabolic patterns at the organ, tissue and cell-type levels [64]. We initially constructed an organ-level map of metabolite-associated programs across normal human tissues. Representative organ-enriched metabolites were selected from significant positive associations after excluding unannotated metabolites and ratio traits. Several gmMAP assignments recapitulated canonical tissue-metabolic relationships. For example, citrate was identified as a prostate-enriched metabolite-associated program, consistent with the well-known citrate-accumulating and secretory phenotype of normal prostate tissue. Creatine was mapped to skeletal muscle, matching the dominant role of the creatine– phosphocreatine system in muscle energy buffering. Glycochenodeoxycholate was enriched in the small intestine, in line with the central role of the intestine in bile-acid transport, enterohepatic recycling and bile-acid receptor signalling. The large intestine was marked by 3-indoxyl sulfate, a host – microbiota co-metabolite derived from intestinal microbial tryptophan metabolism. In addition, pyruvate was assigned to the heart, consistent with the high energetic demand and flexible substrate utilization of cardiac tissue. Eye tissue showed enrichment of a carnitine-associated metabolic program. Retinal photoreceptors have exceptionally high energy demands and are rich in mitochondria, and fatty acid oxidation is thought to contribute to meeting their metabolic requirements. Together, these concordant examples support the ability of gmMAP to recover biologically meaningful organ-specific metabolite-associated programs from transcriptomic atlases (Fig. 5a, Supplementary Figure S5a).

**Fig 5.**
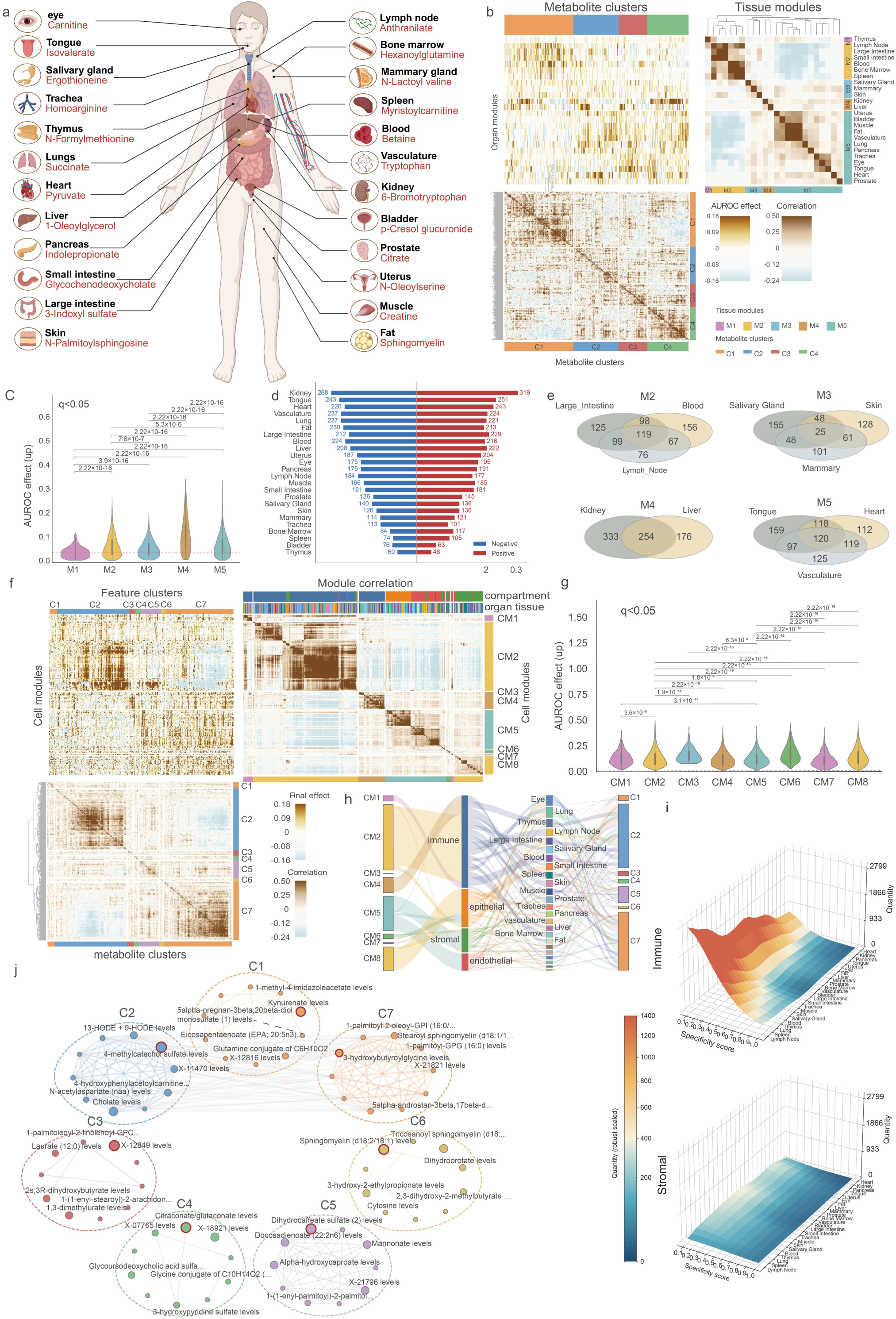
Cross-tissue and cell-type-resolved organization of metabolite-associated programmes inferred by gmMAP. **a.** Schematic illustration of organ-specific elevated metabolic features derived from gmMAP calculation. Figure a was created with BioRender. **b.** Tissue-level mapping of significant positive metabolite associations across healthy human organs. Left, clustered heatmap showing AUROC-derived metabolite association effects across organ modules and metabolite clusters. Bottom, metabolite–metabolite correlation matrix defining four major metabolite clusters. Right, tissue–tissue correlation matrix showing five tissue modules with distinct cross-organ metabolic similarity. **c.** Distribution of up-dominant AUROC effects across the five tissue modules. Statistical comparisons between modules are indicated above the violin plots. **d.** Proportion of significant metabolite features showing positive or negative directional effects across tissues. **e.** Venn diagrams showing the overlap of significant metabolites among representative tissues within tissue modules, highlighting shared module-level metabolic programmes and organ-biased metabolite associations. **f.** Cell-type-level mapping of gmMAP-derived metabolite-associated programmes. Left, clustered heatmap of final metabolite effects across cell modules and metabolite feature clusters. Bottom, metabolite feature correlation matrix identifying seven feature clusters. Right, cell module correlation matrix showing eight cell modules structured by cellular compartment and organ context. Top annotations indicate cellular compartment and tissue origin. **g.** Distribution of up-dominant AUROC effects across the eight cell modules. Statistical comparisons indicate significant differences in metabolite association strength between modules. **h.** Alluvial plot linking cell modules to major cellular compartments, organ tissues and metabolite feature clusters, revealing compartment- and tissue-dependent organization of metabolite programmes. **i.** Tissue specificity–quantity landscapes of cell-type metabolite associations differed markedly between immune and stromal compartments. Immune-associated modules exhibited broad, high-density, and highly conserved metabolite patterns across tissues, while stromal-associated modules displayed narrow, tissue-restricted specificity profiles with low cross-tissue conservation. **j.** Network representation of metabolite feature clusters. Nodes represent metabolites and edges indicate coordinated metabolite associations within the cell-type-resolved metabolic landscape. Dashed circles denote major metabolite communities, including amino-acid-, bile-acid-, sphingolipid-, glycerophospholipid-, acylcarnitine-, steroid- and xenobiotic-related features. Collectively, these analyses show that gmMAP resolves whole-body metabolic heterogeneity into coordinated tissue modules, cell-type modules and metabolite feature communities.

Systematic mapping of metabolite-associated programmes across tissues and tissue-resolved cell types revealed a modular organization of the cellular metabolic landscape (Fig. 5b). Unsupervised clustering of significant positive metabolite associations separated tissues into five tissue modules and metabolites into four major metabolite clusters, uncovering both organ-specific and cross-organ metabolic patterns (Fig. 5b). Among these modules, M2 was mainly composed of immune- and mucosal immune-enriched tissues, including blood, lymph node and large intestine, indicating a shared metabolic programme linked to immune activity and barrier-associated inflammation [65]. M3 comprised secretory and epithelial-barrier-associated organs, including salivary gland, mammary gland and skin, suggesting a convergent metabolic signature related to glandular secretion, lipid handling and epithelial renewal [66–68]. M4 contained the major metabolic organs liver and kidney, consistent with their central roles in systemic metabolite processing, detoxification, amino-acid metabolism and renal solute handling [69,70]. The gmMAP-derived metabolic effects differed significantly across tissue modules, indicating that each module captured a distinct and robust metabolite-associated programme (Fig. 5c). Directionally significant metabolite associations further revealed marked tissue bias, with the kidney displaying the largest repertoire of associated metabolites, followed by tongue, heart and vasculature (Fig. 5d, Supplementary Figure S5b). Within the same tissue module, organs shared a substantial fraction of common metabolite associations, supporting the existence of coordinated cross-organ metabolic programmes among functionally related tissues (Fig. 5e). Together, these functionally coherent modules indicate that tissue metabolic programmes are strongly shaped by organ physiology, with immune, secretory and metabolic organs displaying distinct yet partially shared metabolite-associated landscapes.

At the cell-type level, gmMAP decomposed the cross-tissue metabolic landscape into eight cell modules and seven metabolite feature clusters (Fig. 5f). These modules were structured by cellular compartment and tissue context, partitioning distinct metabolic profiles for immune, epithelial, endothelial, and stromal cells. This supports the paradigm that cell identity and local microenvironment jointly sculpt organ-scale metabolic programs [65,71]. Module correlation analysis revealed non-random, compartment-aligned metabolic patterns, with immune-enriched modules serving as the major driver of cross-tissue metabolic heterogeneity (Fig. 5f).

All cell modules exhibited predominant upregulatory metabolic effects with divergent distribution characteristics, highlighting distinct association strengths and cell-type specificities of metabolic programs across cellular populations (Fig. 5g). Alluvial analysis further integrated cell modules with cellular compartments, tissues and metabolite clusters, revealing differential contributions of immune, epithelial, endothelial and stromal compartments to global metabolic architecture (Fig. 5h). Immune-associated modules displayed broad connectivity across diverse organs and metabolite clusters, representing the core source of cross-tissue conserved metabolic variation and consistent with the central function of metabolic reprogramming in immune cell activity [65]. In contrast, epithelial, endothelial and stromal modules exhibited prominent tissue-biased metabolic features, matching their unique physiological functions, including epithelial barrier maintenance and renewal, vascular homeostasis, and stromal activation [71,72]. Specificity–quantity profiling further confirmed this compartmental divergence: immune modules possessed highly conserved, high-density metabolic signatures across tissues, while stromal modules showed restricted metabolic landscapes with poor cross-tissue conservation. This discrepancy indicates that migratory immune cells maintain stable, consistent metabolic profiles in different tissue microenvironments, whereas stromal cell metabolism is strictly tissue-dependent and context-specific (Fig. 5i). Network visualization of metabolite clusters further uncovered coordinated functional metabolite communities rather than independent metabolic changes (Fig. 5j). These communities encompass metabolites linked to amino acid metabolism, bile acid signaling, sphingolipid turnover, acylcarnitine-mediated fatty acid oxidation, steroid metabolism, glycerophospholipid remodeling and xenobiotic detoxification, all of which are critical for immune regulation, membrane dynamics, mitochondrial energy metabolism and substance clearance [73–76]. The dense intra-cluster connectivity confirms that gmMAP resolves biologically coordinated, cell-type-specific metabolic programs.

We next asked whether metabolite-associated programmes were broadly conserved across organs or instead constrained by tissue context. Analysis of metabolite associations detected in at least two positive tissues revealed a structured continuum of cross-tissue conservation and tissue restriction (Supplementary Fig 5c-h). Immune- and blood-associated cell types, including macrophages, plasma cells, monocytes, natural killer cells and erythrocytes, accounted for the largest number of positive metabolite associations, indicating that haematopoietic and immune compartments represent a major source of shared metabolic variation across organs (Supplementary Fig. 5e,f,g). In contrast, endothelial, epithelial and stromal populations showed more tissue-restricted patterns, suggesting stronger dependence on local organ physiology (Supplementary Fig. 5c,d,e). Levels–ratio paired analysis further showed that lymphoid and immune-rich tissues, including lymph node, spleen, thymus and blood, were enriched for relative metabolite imbalance, consistent with active metabolic remodeling in immune-dominant tissue environments (Supplementary Fig. 5d). By integrating tissue specificity with directional consistency, metabolite–cell-type pairs were resolved into broadly conserved, stable tissue-restricted, diffuse mixed and plastic tissue-restricted classes (Supplementary Fig. 5g,h). Thus, cross-organ metabolic heterogeneity is organized along two axes—cellular compartment and tissue context, with immune-associated programmes forming broadly shared metabolic states and structural or parenchymal compartments exhibiting more organ-specific metabolic specialization.

### Pan-cancer metabolic heterogeneity at single-cell resolution

Although we previously characterized cell-type-specific metabolic landscapes across organs and tissues under healthy conditions, disease-associated alterations in cellular metabolism across organ tissues remain unclear. Metabolic dysregulation is a defining feature of cancer, involving both tumor-intrinsic metabolic reprogramming and systemic metabolic alterations that support tumor growth, immune evasion and adaptation to the tumor microenvironment [77–79]. However, how metabolite-associated programmes are shared or diversified across cancer types, tissue origins and cellular compartments remains incompletely resolved. To fill this gap, we extended gmMAP to pan-cancer single-cell atlases to determine whether tumor tissues exhibit conserved or cancer-type-specific metabolite-associated programmes. Unsupervised embedding separated malignant and non-malignant cellular compartments across cancer types, including epithelial, stromal, endothelial, myeloid, B cell, CD4 T cell, CD8 T cell and ILC populations (Fig. 6a,b). Correlation analysis of cancer-state-level gmMAP profiles further organized tumor states into six cancer-state modules, indicating that pan-cancer metabolic heterogeneity is jointly shaped by cancer type, tissue origin and cellular compartment (Fig. 6c).

**Fig 6.**
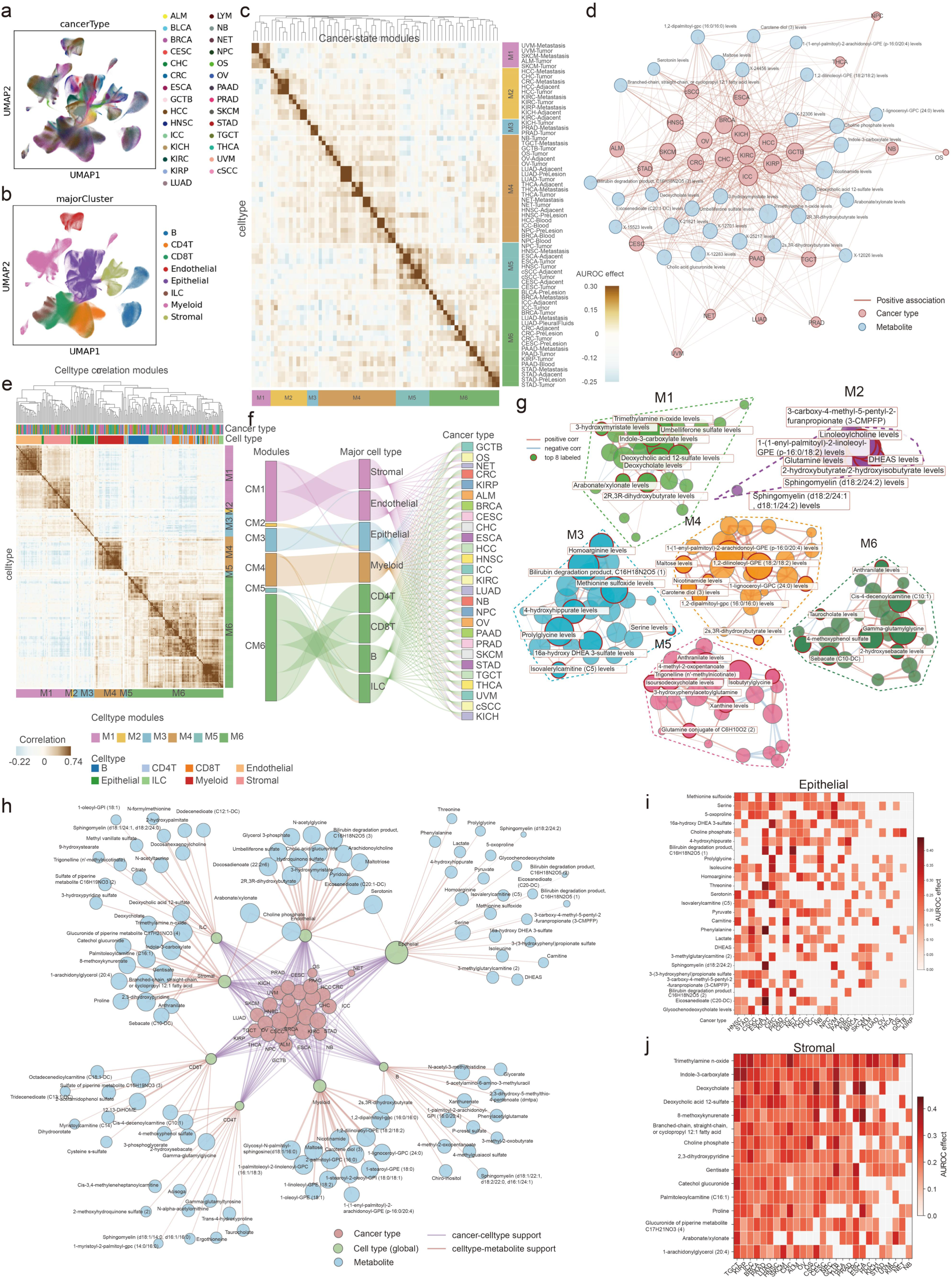
gmMAP identifies shared and cancer-type-specific metabolite programs across pan-cancer single-cell populations. **a.** UMAP visualization of single cells from multiple cancer types, coloured by cancer type. **b.** UMAP visualization of major cellular compartments, including epithelial cells, stromal cells, endothelial cells, myeloid cells, B cells, CD4 T cells, CD8 T cells and innate lymphoid cells. **c.** Correlation heatmap of cancer-state-level gmMAP profiles, revealing six cancer-state modules with distinct metabolic similarity across tumor types and disease states. Rows and columns represent cancer-state or cell-type combinations, and colours denote pairwise correlation. **d.** Cancer–metabolite network showing representative metabolite associations shared across cancer types. Cancer types and metabolites are represented as nodes, and edges indicate significant positive associations; node size reflects connectivity or association frequency. **e.** Cell-type correlation heatmap showing pan-cancer cell modules inferred from metabolite-associated profiles. Top annotations indicate cancer type and major cell compartment. **f.** Alluvial plot linking cell modules, major cell compartments and cancer types, illustrating the contribution of stromal, endothelial, epithelial, myeloid, lymphoid and ILC compartments to pan-cancer metabolic programmes. **g.** Network visualization of metabolite feature modules, identifying coordinated metabolite communities enriched for bile-acid-related, microbiota-derived, sphingolipid, amino-acid, acylcarnitine, glycerophospholipid, steroid and xenobiotic-associated features. Dashed circles denote major metabolite modules. **h.** Integrated cancer–cell-type–metabolite network showing how cancer types are connected to global cell compartments and metabolite programmes. Cancer types, cell types and metabolites are shown as distinct node classes, and edges indicate cancer–cell-type or cell-type–metabolite support. **i.** Heatmap showing representative myeloid-associated metabolite programmes across cancer types, highlighting immune-related metabolic heterogeneity. **j.** Heatmap showing representative stromal-associated metabolite programmes across cancer types. Stromal programmes were enriched for microbiota- and bile-acid-related metabolites, including trimethylamine N-oxide, indole-3-carboxylate, deoxycholate and deoxycholic acid 12-sulfate, suggesting a recurrent microbiota–bile acid–tumor microenvironment axis. Together, these analyses show that gmMAP resolves pan-cancer metabolic heterogeneity into cancer-type-specific, cell-compartment-specific and metabolite-class-specific networks.

Network analysis revealed that tumor-associated metabolites were not randomly distributed across cancer types, but formed coordinated cancer–metabolite communities (Fig. 6d). Several metabolites connected multiple cancer types, suggesting shared metabolic programmes across malignancies, whereas other metabolites showed more restricted cancer-type associations (Fig. 6d,i). These shared networks were enriched for lipid and membrane metabolites, sphingolipids, bile-acid-related features, acylcarnitines, amino-acid derivatives and xenobiotic or microbiota-associated metabolites (Fig. 6d,g,i). This organization indicates that gmMAP captures both intrinsic tumor metabolic rewiring and extrinsic metabolic inputs from the tumor microenvironment.

Cell-type-resolved correlation and network analyses further showed that stromal, myeloid, endothelial and epithelial compartments acted as major intermediates linking cancer types to metabolite programmes (Fig. 6e,f,h). In particular, stromal-associated metabolite signatures were strongly enriched for microbiota-related and bile-acid-related metabolites, including trimethylamine N-oxide, indole-3-carboxylate, deoxycholate and deoxycholic acid 12-sulfate, together with xenobiotic and lipid-related metabolites (Fig. 6h,j). This pattern suggests that the stromal tumor microenvironment may serve as an important sensor and mediator of systemic or microbiota-derived metabolic cues. Consistent with this interpretation, deoxycholic acid has been reported as a gut microbiota-derived secondary bile acid that promotes multiple cancers through inflammatory, oncogenic and immune-suppressive mechanisms [80–83]. In breast cancer, microbiota-derived deoxycholic acid accumulates in tumor tissues and activates tumor-cell-intrinsic FXR signalling, inducing NF-κB-dependent IL-6 production and promoting G-MDSC and Th17-mediated immunosuppression [80]. DCA has also been implicated in colorectal cancer by suppressing CD8 T cell effector functions and promoting tumor growth, in hepatocellular carcinoma through obesity-associated gut microbial metabolism and senescence-associated secretory programmes, and in oesophageal carcinogenesis through IL-6/STAT3 and inflammatory signalling [81–84]. Thus, the enrichment of deoxycholate-related features in the pan-cancer gmMAP network provides biologically plausible evidence for a conserved microbiota–bile acid–tumor microenvironment axis (Fig. 6h,j).

Beyond bile acids, other metabolites highlighted by the network also support known cancer-related metabolic axes. Trimethylamine N-oxide has been linked to colorectal cancer cell proliferation, angiogenesis and inflammation-associated cancer risk [85]. Indole-related microbial metabolites can regulate tumor immunity through aryl hydrocarbon receptor signalling in tumor-associated macrophages and other immune cells [86]. Acylcarnitine and carnitine-related features reflect altered fatty-acid transport and mitochondrial metabolic plasticity, which are increasingly recognized as contributors to tumor adaptation and immune–metabolic remodelling [87]. Together, these results show that gmMAP resolves pan-cancer metabolism into cancer-type-specific, cell-compartment-specific and metabolite-class-specific networks, revealing recurrent microbial, bile-acid, lipid and mitochondrial metabolic programmes across tumor tissues (Fig. 6d–j).

### gmMAP-drug prioritizes clinical drugs for reversing pan-cancer epithelial metabolic programmes

We next asked whether pan-cancer epithelial metabolic programmes could be decomposed into recurrent modules and pharmacologically queried by metabolism-modulating drugs. Consensus NMF of the epithelial metabolite final-effect matrix resolved five epithelial metabolic modules, each defined by distinct metabolite traits and supported by coherent block structures in the module correlation matrix, indicating that these modules captured reproducible epithelial metabolic states rather than random metabolite co-variation (Supplementary Figure S8a). Module activity varied across epithelial disease states, with tumor, metastatic and pleural-fluid epithelial states showing distinct activity patterns compared with adjacent or pre-lesion states, suggesting that malignant progression is accompanied by coordinated remodelling of metabolite-associated programmes (Supplementary Figure S8b). Module composition further differed across cancer types, revealing both broadly shared epithelial metabolic axes and cancer-context-restricted metabolic states (Supplementary Figure S8d).

To connect these tumor metabolic programmes with pharmacological perturbations, we developed gmMAP-drug, a drug-matching framework that compiles clinical or clinically relevant drugs capable of perturbing host metabolism and matches drug-induced transcriptional perturbation signatures to tumor-associated metabolite programmes. By identifying metabolites upregulated in cancer epithelial cells and comparing these tumor-enriched signatures with drug-associated metabolic connectivity profiles, gmMAP-drug prioritizes candidate drugs predicted to reverse tumor-associated metabolic states. This strategy is conceptually related to perturbational signature-matching approaches such as Connectivity Map and LINCS L1000, but operates at the level of metabolite-associated programmes rather than individual gene signatures [88].

gmMAP-drug revealed structured and biologically interpretable drug–metabolite connectivity patterns across the five epithelial modules (Supplementary Figure S8c). Metformin-associated perturbations were linked to nucleotide-related, redox-associated, bilirubin or bile-acid-related and lipid-associated metabolite features, consistent with the known ability of metformin to regulate AMPK–mTOR signalling, mitochondrial metabolism and cancer-cell biosynthetic vulnerability [89,90]. PPAR-targeting perturbations, including pioglitazone and lanifibranor-related signatures, were associated with lipid, acylcarnitine, sphingolipid and bile-acid-related modules, in line with the role of PPAR signalling in lipid handling, inflammation and systemic metabolic homeostasis [91]. Statin-related signatures, represented by simvastatin, were connected to lipid, sterol, acylcarnitine and sphingolipid-associated traits, consistent with inhibition of the mevalonate/cholesterol biosynthesis pathway and its known relevance to cancer growth and survival [92]. GLP-1-related perturbations, including semaglutide-associated and tirzepatide-associated signatures, were connected to energy-balance, amino-acid, lipid, bile-acid and gut-derived metabolite features; these predictions should be interpreted as metabolic-state hypotheses because current clinical evidence for GLP-1 receptor agonists in oncology remains evolving and context dependent [93].

We then applied gmMAP-drug to tumor-versus-adjacent epithelial comparisons across representative cancer contexts. Differential metabolite activity analysis identified cancer-specific sets of tumor-enriched and adjacent-enriched metabolite traits in CRC, STAD, CESC, ESCA and HCC, revealing substantial metabolic asymmetry between malignant and non-malignant epithelial states (Supplementary Figure S8i–m). Matching these tumor-associated metabolite signatures against gmMAP-drug profiles nominated distinct predicted reversing drugs across cancer types, including semaglutide HFD liver, simvastatin, BMS-21 perturbations, GLP-1 perturbations, metformin-associated signatures, pioglitazone and losartan (Supplementary Figure S8i–m). Several of these candidates are supported by prior cancer-related metabolic studies: metformin has been reported to suppress mitochondrial-dependent biosynthesis and affect AMPK–mTOR-associated cancer metabolic programmes [89,90]; statins, including simvastatin, target the mevalonate pathway, a lipid and sterol biosynthesis axis implicated in tumor growth and survival [92]; PPARγ agonists such as pioglitazone can induce metabolic and epithelial changes in cancer cells [94]; and losartan has been shown to remodel tumor stroma, improve tumor perfusion and enhance chemotherapy delivery in preclinical breast, pancreatic and ovarian cancer models [95,96]. GLP-1 receptor agonist-related predictions, including semaglutide-associated signatures, should be interpreted more cautiously, as current evidence mainly comes from cancer-risk analyses and observational or meta-analytic studies rather than direct antitumor intervention trials [93]. The ranked candidates differed among tumor types, indicating that pharmacological reversal of epithelial metabolic programmes may depend on the underlying tumor-specific metabolic state. Bilateral drug–metabolite connectivity plots further identified the metabolite features driving representative predictions: simvastatin was linked to lipid-related metabolites, acylcarnitines, sphingomyelins and bile-acid-associated traits; BMS-21 perturbations were connected to amino-acid, nucleotide, lipid and mitochondrial-associated metabolites; metformin-associated signatures showed strong connections with nucleotide-related ratios, bilirubin degradation products, redox-associated and lipid-related traits; and semaglutide-associated signatures were linked to bile-acid, lipid, acylcarnitine, amino-acid and gut-derived metabolite features (Supplementary Figure S8e–h).

Given the distinct metabolic constraints imposed by the central nervous system, including unique nutrient availability, neuronal–glial metabolic coupling and brain-specific microenvironmental regulation, glioma metabolism may differ substantially from that of extracranial solid tumors; therefore, we further performed a dedicated gmMAP analysis to dissect glioma-specific metabolic alterations and therapeutic vulnerabilities. To evaluate whether gmMAP could resolve metabolic heterogeneity in pediatric brain tumors, we applied it to a longitudinal single-cell atlas of pediatric high-grade glioma [97], which profiled malignant and non-malignant populations across molecular subtypes and disease stages. The atlas contains major tumor-associated cell compartments, including glial cells, oligodendrocytes, neurons, endothelial cells, myeloid cells, pericytes and T cells, providing a suitable framework for cell-resolved tumor–normal metabolic comparison. gmMAP identified subtype-specific tumor-associated metabolic alterations in glial cells from IDH1 R132H, NOS and H3F3A G34R/V initial tumor samples (Supplementary Fig. S6a–c,e,g). NOS tumors showed the strongest metabolic divergence, with tumor-enriched traits including N-acetylserine, glycochenodeoxycholate and phospholipid-related metabolites, whereas normal-enriched traits included octadecanedioylcarnitine and 3-ethylcatechol sulfate. H3F3A G34R/V tumors showed a distinct pattern, including enrichment of N2,N2-dimethylguanosine in tumor cells. Coupling these tumor-associated metabolite signatures with gmMAP-drug nominated subtype-dependent reversing perturbations, including GLP-1-related signatures, BMS-21, semaglutide-associated signatures, metformin-related perturbations, bezafibrate, losartan, pioglitazone and obeticholic acid-associated profiles (Supplementary Fig. S6d,f,h). These results suggest that gmMAP can capture molecular-subtype-specific metabolic abnormalities in pediatric high-grade glioma and nominate candidate metabolism-modulating perturbations that may reverse tumor-associated metabolic states.

Together, these analyses establish gmMAP-drug as a hypothesis-generating framework for linking tumor-upregulated metabolite programmes to metabolism-modulating drugs and prioritizing candidate pharmacological strategies that may reverse tumor-associated epithelial metabolic states for future experimental validation.

### gmMAP reveals gut microbial metabolite-mediated shaping of inflammatory fibroblast in ulcerative colitis

Metabolic reprogramming is a central feature of inflammatory diseases, in which local tissue metabolites and immune–stromal metabolic states shape inflammatory amplification, tissue injury and resolution, yet cell-type-specific metabolic alterations in diseased tissue microenvironments remain poorly resolved [98–101].

To determine whether gmMAP could capture disease-associated metabolic heterogeneity within stromal compartments, we applied it to fibroblast subpopulations from healthy, inflamed and non-inflamed ulcerative colitis colonic tissues. Unsupervised embedding resolved seven fibroblast states (Fig. 7a). CXCL1 expression was strongly enriched in the CXCL1⁺ fibroblast state and was most pronounced in inflamed UC tissue (Fig. 7b, c, d), whereas non-inflamed tissue retained an intermediate signal, suggesting that this population represents an inflammation-associated stromal state rather than a constitutive fibroblast subset. This is consistent with previous single-cell studies showing expansion of inflammatory fibroblasts in UC and IBD lesion [102]. Activated fibroblast programs have also been linked to neutrophil-attractant chemokine production and IL-1-dependent inflammatory tissue pathotypes, supporting the biological relevance of the CXCL1⁺ fibroblast state identified here [103]. gmMAP further revealed that inflammatory fibroblast activation was accompanied by coordinated metabolic remodeling. CXCL1⁺ fibroblasts and related activated fibroblast states were enriched for metabolite traits associated with fatty-acid oxidation and incomplete mitochondrial oxidation, including long-chain acylcarnitines, sebacate/glutarate-related metabolites and hydroxy-fatty-acid intermediates. In parallel, these cells showed higher scores for hypoxic and energetic-stress-associated traits, including lactate, hypoxanthine and adenine nucleotide ratios such as ADP-to-fructose and ADP-to-citrate (Fig. 7e-j). These findings are consistent with metabolomic studies showing broad perturbations of fatty-acid metabolism, lipid metabolism, tricarboxylic-acid-cycle intermediates, amino-acid metabolism and oxidative pathways in IBD [104,105].

**Figure 7.**
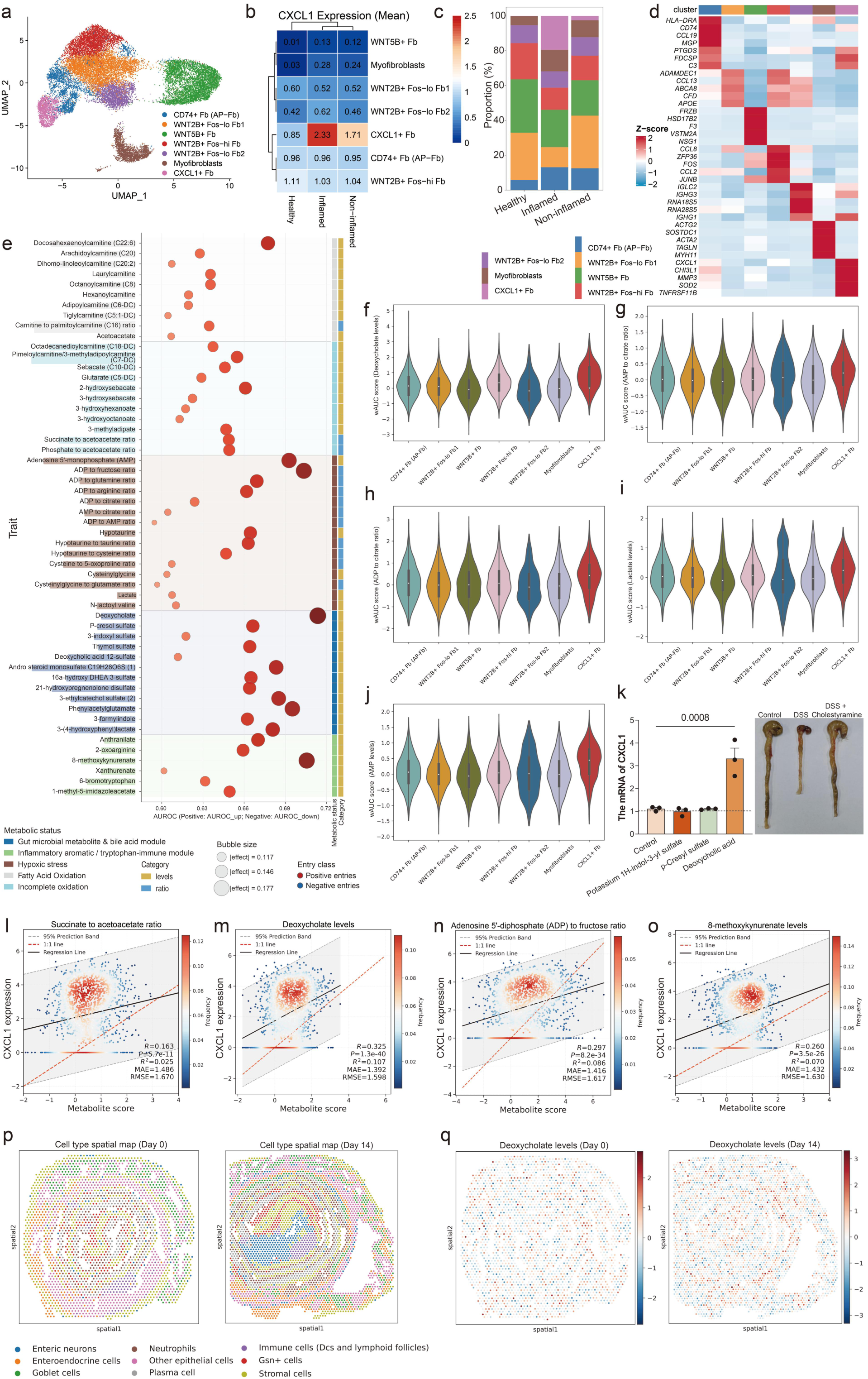
gmMAP identifies metabolically heterogeneous inflammatory fibroblast states in ulcerative colitis. **a.** UMAP embedding of colonic fibroblasts from healthy, inflamed and non-inflamed ulcerative colitis samples, colored by fibroblast subpopulation. Fibroblast states included CD74⁺ antigen-presenting fibroblasts, WNT2B⁺ FOS-low fibroblast subsets, WNT5B⁺ fibroblasts, WNT2B⁺ FOS-high fibroblasts, myofibroblasts and CXCL1⁺ fibroblasts. **b.** Heatmap showing mean CXCL1 expression across fibroblast subpopulations and disease states. CXCL1 expression was highest in the CXCL1⁺ fibroblast population, particularly in inflamed UC tissue. **c.** Stacked bar plot showing the proportional composition of fibroblast subpopulations across healthy, inflamed and non-inflamed tissues. **d.** Heatmap of scaled marker-gene expression across fibroblast clusters, showing antigen-presentation markers, matrix-remodeling genes, myofibroblast markers and inflammatory chemokine programs. **e.** Bubble plot of gmMAP-predicted metabolite–fibroblast associations. Metabolite traits are grouped into functional modules, including gut microbial and bile-acid metabolites, inflammatory aromatic/tryptophan-immune metabolites, hypoxic stress, fatty-acid oxidation and incomplete oxidation. Bubble size indicates the absolute final effect size, and color denotes the direction of the selected metabolite entry. Side annotations indicate metabolic status, metabolite entry class and enriched fibroblast cluster. **f–j.** Violin plots showing representative gmMAP wAUCell metabolite scores across fibroblast subpopulations, including docosahexaenoylcarnitine, AMP-to-fructose ratio, ADP-to-citrate ratio, lactate and AMP-related traits. CXCL1⁺ fibroblasts showed elevated scores for multiple inflammatory and energetic-stress-associated metabolite programs. **K**. Primary human colonic fibroblasts were incubated with vehicle control, 100 μM indoxyl sulfate (potassium 1H-indol-3-yl sulfate), 100 μM p-cresyl sulfate, or 100 μM deoxycholic acid for 24 hours. Total RNA was extracted and CXCL1 mRNA expression was measured by qPCR. GAPDH was used as an internal reference gene for normalization. Data are expressed as fold change relative to the vehicle control group (dashed line) and presented as mean ± SD of three independent biological replicates (each replicate is represented by a black dot). Statistical analysis was performed using one-way ANOVA with Tukey’s multiple comparisons test. Only deoxycholic acid treatment resulted in a statistically significant upregulation of CXCL1 mRNA compared to the control group (p = 0.0008). Bile acid sequestrants for the treatment of colitis model. **l–o.** Single-cell density scatter plots showing the relationship between CXCL1 expression and selected gmMAP metabolite scores, including succinate-to-acetoacetate ratio, deoxycholate levels, ADP-to-fructose ratio and 8-methoxykynurenate levels. Regression lines, 1:1 reference lines and prediction bands are shown. Positive correlations indicate that inflammatory CXCL1 expression is coupled to bile-acid, nucleotide-stress and tryptophan/aromatic metabolite programs in UC fibroblasts. **p.** Spatial transcriptomic maps of healthy control colons (Day 0) and colons collected at the recovery stage following DSS-induced colitis (Day 14). Spots were assigned to major cell populations based on transcriptomic signatures and coloured according to the dominant predicted cell type. **q.** Spatial distribution of gmMAP-derived deoxycholate-associated scores in colonic tissues from healthy control mice (Day 0) and mice at the recovery stage following DSS-induced colitis (Day 14). Each spot is coloured according to the normalized deoxycholate-associated activity score inferred by gmMAP-spatial. Red indicates higher activity and blue indicates lower activity. Colour scales were standardized across samples to enable direct comparison of spatial metabolic patterns.

In addition to host energy metabolism, gmMAP identified a prominent gut microbiota–bile acid–aromatic metabolite axis in inflammatory fibroblasts. Deoxycholate, p-cresol sulfate, 3-indoxyl sulfate, thymol sulfate and DHEA-sulfate-related traits were enriched in the inflammatory fibroblast compartment, indicating that stromal activation is coupled to microbial and bile-acid-associated metabolic programs (Fig. 7e-j). Single-cell correlation analysis further supported a direct link between inflammatory fibroblast activation and metabolite programs. CXCL1 expression was positively associated with the predicted scores of succinate-to-acetoacetate ratio, deoxycholate levels, ADP-to-fructose ratio and 8-methoxykynurenate levels (Fig. 7i-o). This is notable because altered fecal bile-acid profiles and dysregulated secondary bile acids have been repeatedly reported in UC and IBD [106,107], and deoxycholic acid has been functionally linked to gut microbial imbalance and intestinal inflammatory responses in experimental colitis models [108]. gmMAP also highlighted an inflammatory aromatic/tryptophan-immune module, including anthranilate, xanthurenate and 8-methoxykynurenate (Fig. 7e-j). This agrees with prior evidence that inflammation induces tryptophan metabolism through the kynurenine pathway and that kynurenine-pathway metabolites are associated with endoscopic inflammation and disease outcomes in UC [109,110]. To experimentally validate the gmMAP predicted effects of gut microbiota-derived metabolites on colonic fibroblasts, we treated primary human colonic fibroblasts with three representative exogenous metabolites: indoxyl sulfate (potassium 1H-indol-3-yl sulfate), p-cresyl sulfate, and deoxycholic acid. Notably, only gmMAP-predicted deoxycholic acid with the highest correlation to CXCL1⁺ fibroblasts induced a robust and statistically significant upregulation of CXCL1 mRNA expression (p = 0.0008) compared to the vehicle control (Fig. 7k). We further investigated the effects of deoxycholic acid (DCA) on colitis in vivo, continuous cholestyramine gavage efficiently sequestered intestinal bile acids and the microbial metabolite deoxycholic acid, reducing the abundance of bile acid-related pathogenic metabolites in the intestinal microenvironment. The decreased levels of bile acids and DCA effectively attenuated metabolite-mediated intestinal inflammatory damage, thereby ameliorating DSS-triggered colonic pathological atrophy (Fig. 7k). To determine whether bile-acid-associated metabolic programmes remain altered after the resolution of acute inflammation, we applied gmMAP-spatial to spatial transcriptomic sections collected from healthy control mice (Day 0) and mice at the recovery phase following DSS-induced colitis (Day 14) (Fig. 7p). Spatial mapping of genetically informed deoxycholate-associated programmes revealed widespread enrichment across the colonic tissue in both conditions (Fig. 7q). Notably, despite partial recovery of tissue architecture at Day 14, deoxycholate-associated activity remained broadly distributed and showed a greater overall spatial exposure compared with healthy controls. Regions exhibiting elevated deoxycholate-associated scores were observed throughout epithelial, stromal and immune-enriched compartments, indicating that bile-acid-associated signalling persists beyond the acute inflammatory phase (Supplementary Figure S9). These findings suggest that deoxycholate is not merely a marker of acute tissue injury but may contribute to sustained metabolic remodelling during tissue repair and recovery. The persistence of deoxycholate-associated programmes after inflammation resolution further supports a dual role for bile-acid signalling in both inflammatory propagation and post-inflammatory tissue adaptation.

To further characterize the metabolic properties of inflammatory fibroblasts, we applied the enzyme-constrained metabolic modeling framework Compass to the same dataset. Consistent with an activated inflammatory state, Compass identified increased activity of enzyme-supported metabolic reactions in CXCL1⁺ fibroblasts, including fatty-acid activation, lipid remodeling and eicosanoid biosynthesis (Supplementary Fig. SX). Notably, Compass and gmMAP captured complementary aspects of inflammatory fibroblast metabolism. Whereas Compass primarily inferred reaction-level metabolic activity supported by enzyme expression, gmMAP identified metabolite-associated cellular states linked to lactate, secondary bile acids, microbial metabolites and kynurenine-pathway metabolites. These findings suggest that metabolic reaction activity and metabolite-associated regulatory states represent distinct but complementary layers of cellular metabolism. Importantly, because Compass is constrained by intracellular metabolic reaction networks, it is not designed to capture the effects of exogenous metabolites, microbiota-derived metabolites or metabolite-mediated regulatory programs. In contrast, gmMAP linked inflammatory fibroblasts to lactate-, bile-acid- and microbiota-associated metabolite states, thereby enabling the identification of metabolite-driven regulatory mechanisms that extend beyond conventional reaction-centric metabolic models (Supplementary Figure S7g). Together, these results indicate that gmMAP robustly captures intracellular metabolic reprogramming underlying inflammatory fibroblast activation in UC, including mitochondrial substrate remodeling, hypoxic nucleotide stress, and tryptophan-kynurenine immune metabolism, while further enabling the prediction of cellular metabolic responses to exogenous gut microbiota-derived metabolites.

### gmMAP-pseudotime links lactate-associated metabolic remodeling to inflammatory fibroblast differentiation in ulcerative colitis

To investigate whether metabolic remodeling is dynamically coupled to inflammatory fibroblast differentiation, we extended gmMAP to a trajectory-aware framework, termed gmMAP-pseudotime. This approach integrates gmMAP-derived single-cell metabolite scores with pseudotime ordering to quantify metabolite-program dynamics along disease-associated cell-state transitions. Applied to ulcerative colitis fibroblasts, trajectory inference ordered fibroblast states along a continuous differentiation axis, with WNT2B⁺, CD74⁺ and intermediate fibroblast states progressing towards a terminal CXCL1⁺ inflammatory fibroblast state (Fig. 8a). The inferred pseudotime and trajectory streamlines indicated that CXCL1⁺ fibroblasts occupied the late stage of this inflammatory trajectory, suggesting that CXCL1⁺ fibroblasts represent a metabolically and transcriptionally activated endpoint of stromal inflammation (Fig. 8a). This interpretation is consistent with previous single-cell and spatial studies showing that UC and experimental colitis are accompanied by inflammatory fibroblast expansion, stromal remodeling and fibroblast–immune-cell interaction programs [111]. gmMAP-pseudotime identified lactate-associated metabolic programs as prominent features of this inflammatory trajectory. At the single-cell level, CXCL1 expression was positively correlated with predicted lactate levels and N-lactoyl valine levels, linking inflammatory chemokine activation to lactate-related metabolite states (Fig. 8d,e). Along pseudotime, N-lactoyl valine and AMP-associated scores increased progressively, indicating that inflammatory fibroblast differentiation is accompanied by accumulation of lactate-linked and energetic-stress-associated metabolite programs (Fig. 8b,c). These findings suggest that progression towards the CXCL1⁺ state is accompanied by coordinated metabolic remodeling involving glycolysis, lactate metabolism and cellular energetic stress. This is consistent with evidence that lactate can function as a signaling metabolite and promote HIF1A-associated glycolytic remodeling in fibroblasts [112,113]. Consistent with gmMAP predictions, CXCL1⁺ fibroblasts exhibited the strongest enrichment of lactate-associated metabolite programs among all fibroblast populations (Supplementary Fig. S7a), further supporting a close association between lactate metabolism and inflammatory fibroblast activation. Consistent with the gmMAP-pseudotime prediction, genes involved in lactate transport, glycolysis and hypoxia signaling showed marked remodeling across fibroblast states. HIF1A, LDHA and lactate transporter-related genes, including SLC16 family members, were enriched in activated fibroblast states, with HIF1A and LDHA showing prominent upregulation in the CXCL1⁺ fibroblast compartment (Fig. 8f–i,k). Differential expression analysis further confirmed that CXCL1⁺ fibroblasts upregulated inflammatory mediators and metabolic regulators, including CXCL1, CXCL3, CXCL6, HIF1A, SLC16A3 and LDHA, compared with other fibroblast states (Fig. 8g). Immunofluorescence staining further supported this inflammatory hypoxia program in vivo, showing increased HIF1A signal in PDPN⁺ stromal fibroblast regions in DSS-induced colitis compared with control tissue (Fig. 8j). These observations are consistent with previous studies showing that TNF can drive glycolytic reprogramming in fibroblast-like stromal cells through a GLUT1–HIF1A-dependent program and that inhibition of glycolysis can attenuate inflammatory fibroblast phenotypes [114]. They also agree with the broader role of hypoxia/HIF signaling in intestinal inflammation and IBD pathobiology [115].

**Figure 8.**
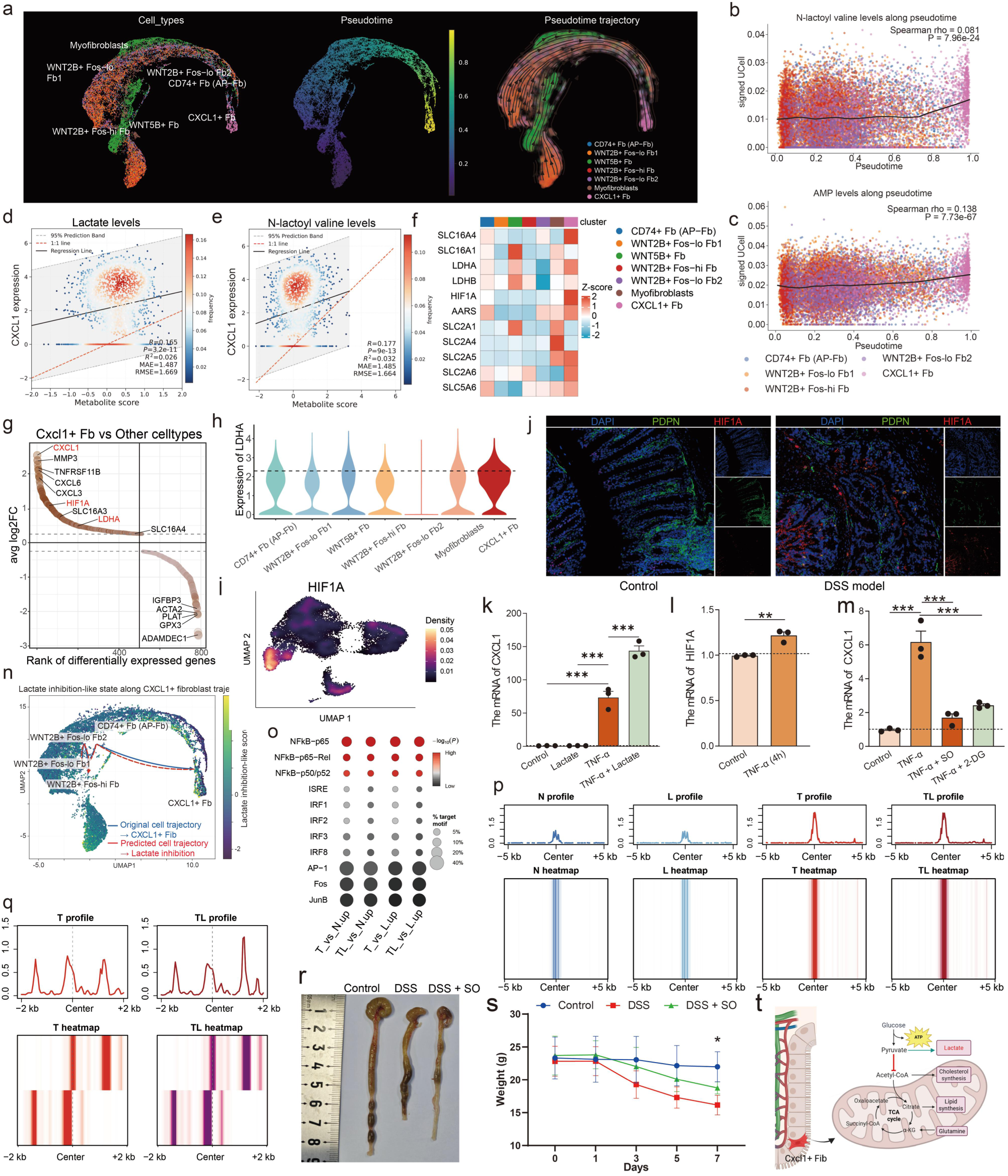
gmMAP-pseudotime links lactate-associated metabolic remodeling to CXCL1⁺ inflammatory fibroblast differentiation in ulcerative colitis. **a.** UMAP visualization of ulcerative colitis fibroblasts colored by fibroblast subtype, inferred pseudotime and pseudotime trajectory. The trajectory orders fibroblast states along a continuous inflammatory differentiation axis terminating in CXCL1⁺ fibroblasts. **b, c.** gmMAP-pseudotime analysis of N-lactoyl valine levels (b) and AMP levels (c) along the CXCL1-associated fibroblast trajectory. Each point represents a single cell colored by fibroblast subtype. Black lines indicate smoothed trends. Spearman correlation coefficients and P values are shown. **d, e.** Single-cell density scatter plots showing the relationship between CXCL1 expression and gmMAP-predicted lactate levels (d) or N-lactoyl valine levels (e). Regression lines, 1:1 reference lines and prediction bands are shown. **f.** Heatmap showing scaled expression of lactate transporters, glycolytic enzymes, hypoxia-associated genes and nutrient-transporter genes across fibroblast clusters. **g.** Ranked differential-expression plot comparing CXCL1 ⁺ fibroblasts with other fibroblast states. Inflammatory chemokines and metabolic regulators, including CXCL1, HIF1A, SLC16A3 and LDHA, are highlighted. **h.** Violin plot showing LDHA expression across fibroblast subpopulations, with elevated LDHA expression in CXCL1 ⁺ fibroblasts. **i.** UMAP density visualization of HIF1A expression, showing enrichment in inflammatory fibroblast regions. **j.** Representative immunofluorescence images of control and DSS-induced colitis tissue stained for DAPI, PDPN and HIF1A. DSS-treated tissue shows increased HIF1A signal in PDPN * stromal regions. **k.** qPCR analysis of CXCL1 mRNA expression after control, lactate, TNF-α or combined TNF- α plus lactate treatment. Lactate alone has limited effects on CXCL1 expression but markedly potentiates TNF- α-induced CXCL1 activation. **l.** qPCR analysis showing induction of HIF1A mRNA after TNF-α stimulation. **m.** qPCR analysis showing that inhibition of lactate metabolism with sodium oxamate or inhibition of glycolysis with 2-deoxyglucose suppresses TNF- α-induced CXCL1 expression. **n.** gmMAP-perturb reveals that inhibition of lactate metabolism blocks the differentiation trajectory of CXCL1⁺ fibroblasts based In vitro transcriptomic profiling of lactate-treated colonic fibroblasts. **o.** Bubble plot showing transcription factor motif enrichment in differentially accessible chromatin regions under four pairwise treatment comparisons (T vs N.up, TL vs N.up, T vs L.up, TL vs L.up). Color gradient represents −log₁₀(P) for enrichment significance (red = high), and dot size reflects the proportion of differential peaks containing each motif. NFκB family motifs display dominant enrichment compared with IRF and AP-1 family motifs across all groups. **p.** ATAC-seq aggregate plots and heatmaps illustrating chromatin accessibility at the CXCL1 promoter, with genomic regions ranging from −5 kb to +5 kb flanking the promoter center. Groups: N (control), L (lactate), T (TNF-α), TL (TNF-α + lactate). Lactate alone fails to alter basal CXCL1 promoter accessibility, and lactate co-treatment does not potentiate TNF-α-triggered opening of the CXCL1 promoter. **q.** ATAC-seq aggregate plots and heatmaps showing chromatin accessibility at a CXCL1 distal enhancer region (−2 kb to +2 kb flanking the enhancer center) in TNF-α-treated (T) and TNF-α plus lactate co-treated (TL) fibroblasts. Lactate co-treatment substantially increases chromatin openness at this CXCL1 enhancer cluster relative to TNF-α stimulation alone. **r, s.** Representative colon morphology (r) and body-weight trajectories (s) from DSS-induced colitis mice treated with sodium oxamate. Lactate-metabolism inhibition partially alleviates disease severity. **t.** Working model. Inflammatory cytokines initiate fibroblast activation, whereas lactate-associated metabolic remodeling enhances chromatin accessibility and reinforces TNF- α-dependent CXCL1 transcription, promoting differentiation towards CXCL1⁺ inflammatory fibroblasts.

To further define the inflammatory signaling programs associated with the CXCL1⁺ fibroblast state, we performed pathway activity inference using decoupleR. CXCL1⁺ fibroblasts exhibited the strongest activation of TNFα-associated transcriptional programs among all fibroblast populations, together with enrichment of inflammatory signaling pathways including NFκB and MAPK signaling (Supplementary Fig. S7b). Consistent with this prediction, canonical TNF-responsive genes, including CXCL1, CXCL3, MMP3 and TNFRSF11B, were preferentially expressed in CXCL1⁺ fibroblasts (Supplementary Fig. S7c). Functional stimulation experiments further demonstrated that TNFα treatment robustly induced expression of CXCL3, MMP3 and TNFRSF11B in primary colonic fibroblasts (Supplementary Fig. S7d–f). These findings indicate that CXCL1⁺ fibroblasts are characterized by a TNFα-dominated inflammatory transcriptional state and suggest that lactate-associated metabolic remodeling operates within a pre-existing cytokine-driven activation program rather than serving as an independent initiating signal. Functional perturbation experiments refined the mechanistic interpretation of the lactate–HIF1A–CXCL1 axis. TNF-α stimulation induced HIF1A expression in colonic fibroblasts (Fig. 8l). In contrast to our initial hypothesis, lactate alone did not substantially induce CXCL1 expression, whereas combined TNF-α and lactate treatment produced markedly stronger CXCL1 induction than TNF-α stimulation alone (Fig. 8k,m). These findings indicate that lactate is insufficient to initiate inflammatory fibroblast activation independently but instead potentiates cytokine-driven inflammatory responses. Consistent with this interpretation, inhibition of lactate metabolism using sodium oxamate or blockade of glycolytic flux using 2-deoxyglucose significantly attenuated TNF-α-induced CXCL1 expression (Fig. 8m), supporting a requirement for glycolytic–lactate metabolism in sustaining inflammatory fibroblast activation.

To systematically dissect the regulatory role of lactate in shaping colonic fibroblast heterogeneity, we performed genome-wide transcriptome profiling on primary colonic fibroblasts subjected to exogenous lactate treatment in vitro. Using single-cell trajectory inference algorithms, we computationally predicted differentiation dynamics of fibroblast subpopulations under normal and lactate-metabolism-suppressed conditions. Our trajectory prediction identified a distinct CXCL1-expressing fibroblast lineage emerging upon lactate stimulation. Notably, inhibition of lactate metabolism markedly disrupted the predicted developmental route toward CXCL1⁺ fibroblasts, effectively halting their progressive differentiation along this specific cell fate trajectory (Fig. 8n). Compared with the control group, both TNF-α treatment and combined TNF-α plus lactate treatment significantly elevated chromatin binding of NFKB family transcription factors at the genomic locus harboring the CXCL1 gene (Fig. 8o). Lactate alone failed to increase chromatin accessibility at the CXCL1 promoter region; moreover, co-administration of lactate did not further augment TNF-α-induced accessibility of the CXCL1 promoter (Fig. 8p). Nevertheless, lactate substantially boosted accessibility at one intronic/enhancer element of CXCL1 (Fig. 8q). Together, these data indicate that lactate remodels the epigenetic landscape to potentiate TNF- α-triggered CXCL1 transcription. Finally, in vivo pharmacological inhibition of lactate metabolism with sodium oxamate partially alleviated DSS-induced colitis, as reflected by improved body-weight maintenance compared with DSS-treated animals (Fig. 8r,s). Together, these findings support a model in which inflammatory cues initiate fibroblast activation, whereas lactate-associated metabolic remodeling functions as an epigenetic and metabolic amplifier that reinforces HIF1A activation, chromatin accessibility and CXCL1 expression during differentiation towards CXCL1⁺ inflammatory fibroblasts (Fig. 8t).

## Discussion

In this study, we developed gmMAP, a genetically informed framework for mapping metabolite-associated programs across single-cell and spatial transcriptomic tissues. By integrating metabolite GWAS-derived gene-level association signals with cellular and spatial transcriptomic profiles, gmMAP provides a scalable strategy to infer cell-type-specific, tissue-resolved and disease-associated metabolic patterns. Unlike conventional single-cell metabolic modelling approaches that mainly rely on enzyme expression and curated intracellular reaction networks, gmMAP uses genetically anchored metabolite-trait gene programs to quantify the transcriptional imprint of diverse metabolic traits, including endogenous metabolites, metabolite ratios, microbiome-derived metabolites, hormone-related traits, dietary metabolites and exposure-associated metabolites. This design enables gmMAP to extend single-cell metabolic inference beyond canonical enzyme-reaction models and to capture a broader spectrum of metabolite-associated cellular states. The gmMap computational framework, together with its five subordinate analytical modules, has been encapsulated into a Python software package with full open-source code freely accessible at https://github.com/Xulab-collab/gmMAP. We also constructed an interactive web portal (https://immunotcm.ac.cn/gmMAP/) to support visualization and bulk download of spatial metabolite mapping data from diverse mouse tissues, alongside trait-mouse tissue association maps.

A major advantage of gmMAP is its ability to connect population-scale metabolite genetics with cellular-resolution biology. Large-scale metabolomic studies have identified extensive associations between circulating metabolites and human diseases, and metabolite GWAS studies have revealed genetic loci that shape inter-individual variation in blood metabolites [5,118,119]. However, these associations are generally measured at the bulk population level and lack tissue or cell-type specificity. gmMAP addresses this gap by projecting metabolite-associated genetic programs onto single-cell and spatial transcriptomic atlases. Through this strategy, metabolite traits can be assigned to specific cell populations, anatomical compartments and disease states, thereby transforming metabolite GWAS signals into interpretable cellular and spatial metabolic maps. This is particularly important for metabolites whose biological effects are context-dependent, such as bile acids, indole derivatives, lipid species, amino acid metabolites, microbiota-derived metabolites and hormone-related traits.

Compared with transcriptome-derived flux inference methods such as Compass, scFEA and related approaches, gmMAP provides a complementary perspective rather than a direct replacement [11, 12, 120]. Compass-like models infer flux-like reaction activities based on the expression of metabolic enzymes and curated gene–protein–reaction relationships. Although these methods are useful for reconstructing intracellular metabolic reaction potential, their accuracy is inevitably limited by the sparsity and dropout of single-cell RNA-seq data, the incomplete correspondence between mRNA abundance and enzyme activity, and the lack of direct information on substrate availability, post-translational modification, allosteric regulation and subcellular compartmentalization. In addition, such models are generally optimized for well-annotated endogenous pathways and are less suited to capture exogenous, microbiome-derived, drug-related or hormone-associated metabolic features. gmMAP reduces these limitations by using metabolite-trait gene programs derived from human genetic associations, allowing metabolic inference to be anchored to metabolite-level phenotypes rather than only to enzyme expression. Therefore, gmMAP is especially suitable for identifying metabolite-associated cellular programs, disease metabolic remodeling and spatial metabolic organization across large-scale transcriptomic datasets.

Our analyses demonstrate that gmMAP can recover biologically meaningful metabolic organization across diverse biological systems. In human multi-tissue and mouse whole-body spatial analyses, gmMAP identified metabolite programs with clear tissue and anatomical specificity, supporting its ability to reconstruct spatially organized metabolic landscapes from transcriptomic data. The rapid expansion of single-cell reference atlases, such as Tabula Sapiens, has provided a foundation for systematic cell-type-resolved molecular annotation across human tissues [64]. gmMAP builds on this resource space by adding a metabolite-trait layer to transcriptomic cell states. In brain regions, gmMAP captured neuron-related metabolic traits such as glutamate and choline-containing lipid species, consistent with known roles of neurotransmitter metabolism and membrane lipid organization in neural tissues. In kidney and liver regions, gmMAP resolved metabolites associated with amino acid, lipid and mitochondrial metabolism, reflecting the specialized metabolic functions of these organs. These results suggest that gmMAP can identify both expected tissue metabolic features and previously underappreciated metabolite-tissue associations.

The developmental kidney analysis further highlights the utility of gmMAP for studying metabolic dynamics during cell differentiation. By applying gmMAP to nephron developmental trajectories, we found that distinct nephron lineages displayed lineage-specific metabolic programs. Proximal tubular development was associated with enhanced lipid, carnitine, acylcarnitine and fatty-acid oxidation-related traits, whereas early nephron progenitor and pretubular states showed stronger biosynthetic, one-carbon and amino acid-associated metabolic programs. These findings are consistent with a metabolic transition from proliferative and biosynthetic precursor states toward mitochondrial and fatty-acid oxidative programs during tubular maturation. By combining metabolite-associated scoring with pseudotime and pseudo-flux analyses, gmMAP provides a framework for linking metabolic remodeling to developmental progression.

In the pan-cancer epithelial analysis, gmMAP revealed substantial metabolic heterogeneity across tumor types and epithelial states. Tumor and adjacent epithelial cells showed distinct metabolite-associated programs, indicating that malignant transformation is accompanied by cancer-type-specific metabolic rewiring. Importantly, gmMAP captured not only tumor-intrinsic metabolic programs but also metabolite traits related to the tumor microenvironment, including gut microbiota-associated metabolites, bile acid derivatives and indole-related metabolites. These findings support the concept that cancer metabolism is shaped by both intrinsic epithelial programs and extrinsic metabolic inputs from host–microbiome and tissue microenvironmental interactions. Furthermore, by matching tumor-associated metabolite signatures with drug perturbation profiles, gmMAP-drug nominated candidate metabolism-modulating drugs capable of reversing cancer-associated epithelial metabolic programs. This analysis provides a translational extension of gmMAP by connecting metabolite-associated cellular states with potential therapeutic interventions.

The ulcerative colitis analysis demonstrates the ability of gmMAP to resolve disease-associated metabolic remodeling in inflammatory stromal states. Single-cell studies of ulcerative colitis have shown extensive epithelial, stromal and immune remodeling in inflamed human colon, including disease-associated fibroblast states and inflammatory cellular circuits [102]. Spatial colitis studies have further demonstrated that intestinal inflammation is accompanied by coordinated cellular biogeography and fibroblast trajectory remodeling [111].

Consistent with these observations, gmMAP identified distinct metabolite-associated programs linked to CXCL1-positive inflammatory fibroblast activation. Pseudotime analyses using independent trajectory methods further suggested that inflammatory fibroblast progression is accompanied by coordinated metabolic changes, including bile acid-, steroid-, indole-, sulfur amino acid- and exposure-associated metabolite programs. These results indicate that gmMAP can capture disease-relevant metabolic remodeling in non-malignant inflammatory tissues and nominate metabolic features associated with pathogenic cell-state transitions.

Another strength of gmMAP is its flexible output structure. At the cell or spatial-spot level, gmMAP generates up, down and net metabolite-associated scores, which can be used for visualization, clustering, differential analysis and pseudotime association. At the cell-type level, gmMAP provides AUROC-based directional association statistics and final signed effects, enabling systematic prioritization of metabolite–cell-type relationships. At the pathway level, gmMAP can incorporate metabolic priors, including pathway topology, reaction order, enzyme annotation and compartment information, to estimate pseudo-metabolic-flow activities. This multi-layered design allows gmMAP to bridge metabolite-trait inference, cellular annotation, spatial mapping, developmental dynamics and drug prioritization within one analytical framework.

Nevertheless, several limitations should be considered. First, gmMAP relies on the quality and coverage of available metabolite GWAS datasets. Metabolites with weak genetic signals, limited sample size or incomplete annotation may yield less reliable gene programs. Second, gmMAP infers metabolite-associated transcriptional programs rather than directly measuring metabolite abundance or biochemical flux. Therefore, its predictions should be interpreted as genetically informed metabolite-associated cellular states or flux-like potentials, ideally validated by spatial metabolomics, isotope tracing, targeted metabolomics or perturbation experiments. Third, genetic associations derived from circulating metabolite measurements may not always reflect local tissue metabolite concentrations, particularly for metabolites with strong tissue-specific production, rapid turnover or compartmentalized activity. Fourth, although gmMAP reduces dependence on individual enzyme expression, it still uses transcriptomic data as the cellular readout and therefore remains affected by cell-state composition, gene detection depth and batch effects. Future integration with proteomics, spatial metabolomics, single-cell metabolomics and perturbation datasets will further improve the biological resolution and causal interpretation of gmMAP predictions.

In summary, gmMAP provides a generalizable framework for linking human metabolite genetics to single-cell and spatial transcriptomic states. By overcoming several limitations of enzyme-expression-based metabolic modelling and extending analysis to endogenous, exogenous, microbiome-derived and hormone-related metabolic traits, gmMAP enables systematic discovery of cell-type-specific metabolic programs, spatial metabolic landscapes, developmental metabolic transitions, disease-associated metabolic remodeling and candidate metabolite-targeted therapies. This framework offers a scalable computational bridge between metabolomics, genetics and cellular tissue biology, and may facilitate the mechanistic understanding of metabolic regulation in development, homeostasis and disease.

## Methods

### Input datasets for the gmMAP workflow

The workflow takes as input: (i) metabolite GWAS summary statistics converted into gene-level association Z-scores; (ii) a single-cell or spatial transcriptomic expression matrix X, with observations corresponding to cells or spatial spots and columns corresponding to genes; (iii) cell-type or spatial-domain annotations for one-vs-rest association testing; (iv) metabolite trait metadata, including metabolite identifiers, reported metabolite names, and trait categories such as metabolite levels and ratios; and (v) curated prior knowledge of metabolic reactions used for pseudo-metabolic flux inference. This prior knowledge included pathway and module assignments of metabolites, reaction ordering within metabolic processes, enzyme–reaction annotations, subcellular compartment constraints, and cofactor dependencies, which together were used to construct a reaction-informed metabolic flow model.

In the implementation, only annotated cell populations with at least 20 observations were retained for downstream enrichment and association analyses. Expression matrices were library-size normalized to 10⁴ counts per observation and log-transformed when raw counts were provided, whereas matrices that already appeared log-transformed or scaled were not re-transformed. Metabolite traits were further stratified according to their metadata-defined categories, allowing metabolite abundance traits and ratio traits to be analysed and interpreted separately.

### Metabolite GWAS data

Metabolite GWAS summary statistics were obtained from the plasma metabolome GWAS reported by Chen et al [6]. This study performed genome-wide association analyses for 1,091 circulating blood metabolite levels and 309 metabolite ratios, providing a large-scale genetic resource for dissecting the inherited regulation of the human plasma metabolome. Plasma metabolites in the source study were quantified using an untargeted ultra-high-performance liquid chromatography–tandem mass spectrometry platform, also referred to as the Metabolon HD4 platform. After quality control and batch normalization, metabolites with measurements missing in fewer than 50% of samples were retained, yielding 1,091 metabolite traits and 309 metabolite ratios for downstream GWAS. Metabolite ratios were constructed from biologically related metabolite pairs sharing enzymes or transporters according to HMDB-based annotations. The original study identified genetic associations for 690 metabolites across 248 loci and for 143 metabolite ratios across 69 loci. The European GWAS data were used in the present analysis to maintain ancestry consistency with downstream gene-level mapping.

In annotation table, the 1,091 metabolite-level traits were classified into ten super-pathway groups: Lipid (395 metabolites), Amino Acid (210), Xenobiotics (130), Nucleotide (33), Cofactors and Vitamins (31), Carbohydrate (22), Peptide (21), Energy (8), Partially Characterized Molecules (21), and Unknown (220). The Xenobiotics category was retained as the major exogenous metabolite class, including sub-pathways such as food component/plant-derived metabolites, benzoate metabolism, chemical exposures, bacterial/fungal metabolites, and drug-related compounds. Steroid hormone-related metabolites were mainly annotated within the lipid super-pathway and included androgenic steroids, pregnenolone steroids, progestin steroids, and corticosteroids. Sub-pathway annotations were used for more fine-grained metabolic interpretation, including lipid subclasses, amino acid metabolism, bile acid metabolism, nucleotide metabolism, vitamin/cofactor metabolism, xenobiotic metabolism, and partially characterized or unknown metabolite groups. Supplementary Tables 1 were provided with the source publication.

For downstream analyses, metabolite identifiers were matched to their reported trait names and pathway annotations. Metabolites and metabolite ratios were kept as separate trait classes because metabolite ratios represent relative biochemical relationships rather than absolute metabolite abundance. When gene-level association scores were required, the harmonized GWAS summary statistics were subsequently converted to gene-level Z-scores using MAGMA-based SNP-to-gene mapping.

### Human single-cell transcriptomic datasets

Single-cell RNA-sequencing datasets from multiple human tissues and disease contexts were collected for gmMAP analysis. The human single-cell transcriptomic atlas released by the Tabula Sapiens Consortium contains 483,152 cells from 254 major cell types across 24 organs [64]. We also obtained a pan-cancer single-cell RNA-sequencing dataset comprising 4,146,975 cells from 29 cancer types, covering eight major cell lineages and 120 refined cell types [121]. In addition, human kidney developmental single-cell RNA-sequencing data were collected, including 17,962 single cells from 22 refined cell types across five developmental stages [22]. We further included the human brain single-cell transcriptomic dataset reported by Siletti et al., which profiled 2,480,956 cells from 88 brain tissue regions [48]. The pediatric high-grade glioma single-cell RNA-sequencing dataset reported by Sussman et al. was also incorporated, comprising 401,253 cells from seven major cell types across four molecular subtypes of pediatric high-grade glioma [97]. Finally, single-cell transcriptomic data from colon tissues of patients with ulcerative colitis and healthy individuals were included, containing 366,650 cells [102]. We followed the quality-control and preprocessing strategies implemented in the original studies, including filtering based on mitochondrial transcript proportion and gene detection thresholds, doublet removal, normalization, log-transformation and batch correction. Publicly available processed expression matrices released by the original authors were directly used for downstream gmMAP analyses. For all datasets, expression matrices that had been normalized by size factors and log-transformed were used as input for the gmMAP framework and downstream analyses.

### Spatial transcriptomics datasets

Spatial transcriptomic data from whole-body mouse sections were obtained, including datasets generated under both steady-state conditions and after LPS treatment. These data comprised 2,336,059 cells across 17 organs and tissues [40]. Spatial transcriptomic preprocessing, including spot-level quality control, normalization, spatial deconvolution and cell-type annotation, was performed according to the procedures and parameter settings described in the original study. The processed expression matrices and spatial annotations released by the original authors were used for all downstream gmMAP analyses.

### Spatial metabolomics data

To validate the metabolite prediction capability of gmMAP, we collected publicly available spatial metabolomics datasets, including spatial metabolomic profiling data of kidney cells during human kidney development. This dataset enables metabolome measurement at cell-type resolution [23].

### Human high-grade glioma specimens

Human high-grade glioma specimens were obtained from patients undergoing surgical resection at Sanbo Brain Hospital, Capital Medical University. The study was reviewed and approved by the Ethics Committee of Sanbo Brain Hospital, Capital Medical University (approval no. SBNK-YJYS-2025-042-01). Written informed consent was obtained from patients where applicable; the use of residual clinical specimens and/or waiver of informed consent was approved by the ethics committee in accordance with the approved protocol and relevant ethical regulations. Fresh tumour tissues were rapidly embedded in optimal cutting temperature compound and stored at −80 °C until further processing. For each specimen, serial adjacent cryosections were prepared to enable paired spatial transcriptomic and spatial metabolomic profiling from matched tissue regions.

### Visium HD spatial transcriptomics

Visium HD spatial transcriptomic profiling was performed on fresh frozen high-grade glioma sections with the assistance of LC-Bio Technology Co., Ltd. Tissue quality was evaluated before library preparation. RNA integrity was assessed using the RNA integrity number, and samples with RIN ≥ 4 were used for Visium HD profiling. Tissue morphology was further examined by DAPI and haematoxylin and eosin staining. According to the 10x Genomics Visium HD Fresh Frozen Tissue Preparation Handbook, 10-μm-thick frozen sections were placed on Visium HD slides and stored at −80 °C until use. Sections were subsequently stained with haematoxylin and eosin and imaged to document tissue morphology. Probe hybridization, probe ligation, slide preparation, probe release, extension and library construction were performed following the Visium HD Spatial Gene Expression Reagent Kits User Guide. Sequencing libraries were generated according to the manufacturer’s protocol and sequenced on an Illumina NovaSeq X Plus platform using paired-end reads. Raw sequencing data were processed using the 10x Genomics Space Ranger pipeline to generate spatially barcoded gene-expression matrices. Downstream quality control, normalization, dimensionality reduction, clustering and visualization were performed using standard single-cell and spatial transcriptomic analysis workflows.

### MALDI2 spatial metabolomics

Spatial metabolomic profiling was performed on adjacent frozen sections from the same high-grade glioma specimens using matrix-assisted laser desorption/ionization mass spectrometry imaging on a Bruker timsTOF fleX MALDI2 platform. Frozen tissues were sectioned at 10 μm using a Leica CM1950 cryostat. Tissue blocks were equilibrated in the cryostat chamber before sectioning, and sections were mounted onto MALDI2-compatible conductive glass slides. After mounting, sections were dried under vacuum and stored at −80 °C until matrix application and mass spectrometry imaging. For matrix deposition, NEDC matrix solution was uniformly sprayed onto tissue sections using an HTX TM-Sprayer M3 system. The matrix solvent consisted of 70% methanol with 3.4% ammonium hydroxide. Matrix spraying was performed with a nozzle temperature of 60 °C, nozzle speed of 1,200 mm min−1, flow rate of 0.08 ml min−1, nitrogen pressure of 10 psi and track spacing of 3 mm. After matrix application, slides were loaded onto the Bruker timsTOF fleX MALDI2 platform. Imaging regions were defined using Bruker Data Imaging software based on tissue morphology. Mass spectrometry imaging was performed in negative-ion mode at 10-μm spatial resolution, with an m/z detection range of 50–1,300, laser frequency of 10,000 Hz, laser energy of 70% and 200 laser shots per pixel.

### Processing and annotation of spatial metabolomics data

Raw MALDI2 imaging data were imported into SCiLS Lab for preprocessing, including baseline subtraction, peak alignment, smoothing and normalization. Root-mean-square normalization was used to reduce pixel-level intensity variation and to support downstream spatial denoising and tissue-region segmentation. Metabolite features were extracted and annotated using Tidymass by matching observed m/z values to theoretical metabolite masses in public metabolite databases. Features were annotated within a mass error threshold of <10 ppm, and, when multiple candidate annotations were available, the candidate with the smallest ppm error was used for downstream analysis. Annotated metabolites were further summarized by chemical classification and mapped back to their spatial coordinates to visualize metabolite distributions across glioma tissue regions.

### Integration of adjacent spatial transcriptomic and metabolomic sections

For paired spatial multi-omic analysis, adjacent tissue sections profiled by Visium HD and MALDI2 spatial metabolomics were aligned using matched tissue morphology from haematoxylin and eosin images. Shared anatomical and tumour regions were identified across adjacent sections, and spatial coordinates were transformed to a common tissue reference where appropriate. Spatial transcriptomic clusters, cell-state annotations and region-level gene-expression signatures were then compared with metabolite ion-intensity maps to identify spatially coordinated transcriptional and metabolic programmes in high-grade glioma tissues.

### Construction of metabolite trait–gene association Z-score matrices

Metabolite-associated GWAS summary statistics based on the GRCh38 genome build were obtained as described above. Because the major histocompatibility complex (MHC) region on chromosome 6, defined as 28,510,120 bp-33,480,577 bp, exhibits complex linkage disequilibrium (LD) patterns and distinctive genetic architecture [122], variants within this region were excluded prior to downstream analysi, in which MHC-region genes may disproportionately influence genome-wide genetic effect estimates. In parallel, an MHC-retained gene-level association matrix was generated as an HLA-aware auxiliary analysis. This version was not used as the default genome-wide analysis but was retained for dedicated immune-related investigations, including HLA-associated traits, autoimmune diseases, infection-related traits and antigen-presentation-related metabolic programs. Comparisons between MHC-excluded and MHC-retained results were used to assess the extent to which MHC-region signals dominated metabolite-associated genetic mapping.

The MAGMA framework [20] was used to perform hierarchical parsing and mapping of genome-wide genetic association data. Based on chromosomal positional information, SNPs were mapped and annotated to genes, and variant-level GWAS association signals within each gene region were aggregated to generate gene-level association statistics. This procedure converted locus-level GWAS signals into gene-level association scores. During analysis, metabolite traits with a sample size smaller than 50 were excluded to ensure sufficient statistical robustness. All other MAGMA settings were kept at their default values. Because only European-ancestry metabolite GWAS summary statistics were included in this study, a European-ancestry LD reference panel [123] was used for MAGMA-based gene-level association analysis. Finally, gene-level genetically mapped Z-score matrices were generated for all metabolite traits.

### Trait-specific up and down gene signatures

For each metabolite trait *t*, the corresponding column of the MAGMA-derived Z-score matrix was extracted as a gene-level association vector, *z_g,t_*. Genes with *z_g,t_* > 0 were ranked in descending order to define the up-signature, whereas genes with *z_g,t_* < 0 were ranked in ascending order to define the down-signature. Only the top 1000 genes derived from each direction were retained for subsequent analysis. For the down-signature, the sign was flipped so that all retained down genes carried positive magnitudes for downstream weighting. Therefore, for each trait *t*, two directional gene sets were constructed: *G_t_*^up^ = {*g*: *z_g,t_* > 0}, *G_t_*^down^ = {*g*: *z_g,t_* < 0}, with the down-weight magnitude defined as *z_g,t_*. Traits with fewer than 200 matched genes in both directions were excluded from scoring.

### Gene variability adjustment and weight assignment

To reduce the influence of genes with unstable expression profiles, each gene was assigned an expression-variability penalty based on its across-observation variability. For gene *g*, the variability term was estimated as 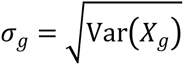, where *X_g_* denotes the normalized expression vector of gene g across all cells or spatial spots. Genes with non-finite or zero variance were excluded. For each retained metabolite-associated gene, a genetically informed and variability-adjusted weight was then defined as *w_g,t_* = 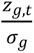, where *z_g,t_* represents the direction-corrected MAGMA-derived association score between gene g and metabolite trait *t*. This weighting scheme assigns larger weights to genes with stronger genetic associations and more stable expression profiles, while reducing the contribution of highly variable genes that may otherwise dominate rank-based enrichment scores.

### Genetically informed metabolite scoring for single-cell, spatial and bulk transcriptomic data

We implemented a weighted AUCell-like rank-based enrichment score, optimized for single-cell and spatial transcriptomic data, to quantify genetically informed metabolite programs in each cell, spatial spot or expression profile. For each metabolite trait *t*, genes were first selected from MAGMA-derived gene-level association *Z*-scores, and gene-specific weights were defined as 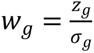, where *σ_g_* denotes gene-wise expression variability across observations. Within each observation *c*, genes were ranked by expression and the top fraction of genes was retained, with *K* = [0.05×*G*] where *G* is the total number of genes. The weighted AUCell-like score was then computed as:

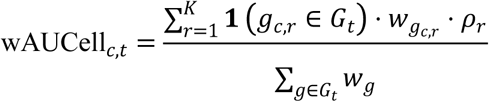

where *g_c,r_* denotes the gene ranked at position *r* in observation *c*, *G_t_* is the trait-associated gene set, and *ρ_r_* is a decreasing positional weight assigned according to within-observation rank. This formulation gives greater contribution to genes that are both genetically prioritized and highly ranked within each expression profile. In practice, it constructs observation-by-trait matrices for subsequent staged normalization and one-versus-rest association analysis. As the wAUCell-like method focuses on within-observation rank enrichment, it is highly applicable to sparse single-cell and spatial transcriptomic data.

For bulk RNA-seq analysis, we adapted sMRS as a sample-level genetically informed metabolite scoring approach. Briefly, the bulk expression matrix *X* was organized as samples by genes and was library-size normalized followed by log-transformation when raw counts were provided. For each metabolite trait *t*, gene-level GWAS/MAGMA *Z*-scores were separated into an up program and a down program by selecting the top 1,000 positively associated genes and the top 1,000 negatively associated genes, respectively, with the down program converted to positive magnitudes for weighting. Gene-specific noise was estimated across bulk samples as 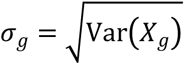, and the genetically informed weight for gene *g* in trait *t* was defined as *w_g,t_* = 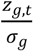. The sample-level metabolite relevance score was then calculated as a normalized weighted sum:

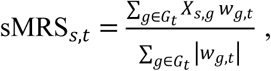

where *s* indexes bulk RNA-seq samples. Only traits with at least 200 valid matched genes were retained.In the bulk setting, this score can be interpreted as a sample-level genetically informed metabolite activity proxy derived from coordinated expression of metabolite-associated genes. Detailed derivations of formulas and symbol definitions are provided in the Supplementary information.

### Sequential normalization of metabolite score matrices

To harmonize metabolite score distributions across traits and observations, raw up and down score matrices were normalized using the S3 strategy, in which trait-wise standardization was first performed across observations, followed by observation-wise standardization across traits. For each cell or spatial spot *c* and metabolite trait *t*, the S3-normalized score was defined as

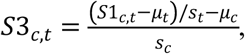

where *S1_c,t_* denotes the raw metabolite score, *μ_t_* and *s_t_* are the mean and standard deviation of trait *t* across all observations, and *μ_c_* and *s_c_* are the mean and standard deviation of the trait-standardized scores across all traits within observation *c*. Rows or columns with zero or non-finite standard deviation were set to zero after normalization to ensure numerical stability.

### One-vs-rest analysis identifies cell type associated metabolic programs

To quantify metabolite specificity for each annotated cell population, we performed one-vs-rest association testing for the metabolite scores. For a given cell type *k* and metabolite trait *t*, observations belonging to cell type *k* were treated as the positive group and all remaining observations as the reference group. Let *n_i n_* and *n_out_* denote the number of observations inside and outside the target cell type, respectively, and let *U* be the Mann–Whitney rank-sum statistic computed from the corresponding score vector. The one-vs-rest AUROC was defined as 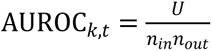, and the centered effect size was defined as effect*_k,t_* = AUROC*_k,t_* − 0.5.

To determine whether a metabolite was preferentially linked to a cell type through its up or down genetic program, we compared the two directional AUROCs. The directional contrast was defined as ΔAUROC*_k,t_* = 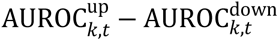. Unlike a simple variance approximation on a single *U* statistic, the significance of ΔAUROC was evaluated using DeLong’s test for two correlated AUCs by default, with a stratified bootstrap option implemented as an alternative. DeLong testing was performed in a two-sided manner, with optional stratified subsampling for very large cell populations to improve computational efficiency.

A final signed association was assigned to each metabolite–cell-type pair by integrating the directional AUROC effects (ΔAUROC) from the up and down metabolite-associated gene programs. The final effect value and its direction between celltypes and traits were determined by ΔAUROC. To control for multiple testing, *p* values were adjusted using the Benjamini–Hochberg procedure in two complementary ways: across cell types for each metabolite trait and across metabolite traits for each cell type. This generated both trait-wise and cell-type-wise false discovery rates. In downstream analyses, significant metabolite–cell-type associations were prioritized using FDR control together with the magnitude of the signed effect. Specifically, significant links were defined by q<0.05 and a sufficiently large final effect. A stringent empirical effect cutoff was estimated as the 95th percentile of final_effect among background metabolite–cell-type pairs with q>0.20, thereby enriching for robust directional associations.

### gmMAP-flux: Enzyme-constrained inference of single-cell metabolic flux potential

We developed gmMAP-flux, an enzyme-constrained metabolic-flow potential model to infer directional metabolic activity across kidney developmental cell states. gmMAP-flux integrates metabolite-linked wAUCell-S3 scores, metabolite-ratio directionality and transcriptome-derived feasibility constraints, including enzyme expression, subcellular-compartment support and cofactor-associated gene programs. The model estimates relative metabolic-flow potential and does not represent stoichiometric flux-balance analysis or absolute biochemical flux.

Each metabolic module was defined by an input metabolite trait, an output metabolite trait and, when available, a metabolite-ratio trait with a curated direction sign. For module m in cell type *c*, the input, output and ratio scores were denoted as *s-in*, *s-out* and *s-ratio*, respectively. Directional evidence was first estimated from the output-input difference:

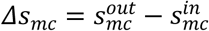

Module activation was calculated as a bounded function of the absolute input and output metabolite signals:

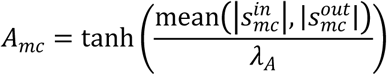

where λ*_A_* is the activation-scale parameter. When a matched ratio trait was available, the endpoint-derived direction was adjusted by ratio-derived support:

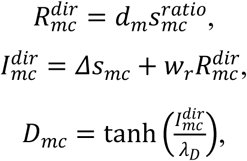

where *d_m_* is the curated ratio direction sign, *w_r_* is the ratio weight and λ*_D_* is the direction-scale parameter. For modules without an available ratio trait, direction was estimated from the input-output difference alone. Assigned but missing ratio traits were penalized, whereas modules without ratio annotation were not penalized. Directional concordance between endpoint-derived and ratio-derived signs was further encoded as a soft consistency factor.

To estimate transcriptomic feasibility, enzyme genes associated with each module were intersected with the gene-expression matrix. Enzyme-gene expression support was calculated using a bounded top-k mean expression score and combined with enzyme-gene coverage. For modules with annotated enzyme genes, enzyme capacity was defined as:

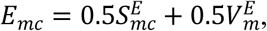

where S-enzyme denotes the bounded enzyme-expression score and V-enzyme denotes the fraction of enzyme genes detected in the expression matrix. Modules without enzyme-gene annotation were assigned a default no-enzyme factor. Compartment and cofactor support were computed analogously from curated hallmark gene panels matched to pathway class, subpathway annotation and metabolite names.

The final feasibility score, used as module confidence, combined metabolite-node support, trait coverage, ratio support, directional consistency and transcriptome-derived soft constraints:

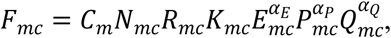

where *C* is input/output trait coverage, *N* is metabolite-node support, *R* is ratio support, *K* is directional consistency, and *E*, *P* and *Q* represent enzyme capacity, compartment support and cofactor support, respectively. The default constraint weights were:

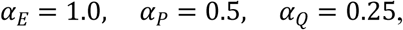

The signed metabolic-flow potential was then calculated as:

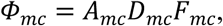

Positive values indicate output-dominant or forward-supported metabolic-flow potential, whereas negative values indicate input-dominant or reverse-supported potential. The magnitude of the signed potential reflects the relative strength of the inferred flow potential.

For subpathway-level analysis, module-level scores were aggregated using feasibility as weights. For a module-level metric *Y*, the corresponding subpathway-level score was calculated as:

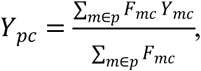

Module- and subpathway-level matrices were exported for activation, direction, confidence, signed flux, ratio support, enzyme capacity, compartment support, cofactor support, mean metabolite score and input-output score difference. Full mathematical definitions, including ratio penalties, consistency factors, node support, gene-set scoring and default missing-annotation factors, are provided in the Supplementary Methods. Detailed derivations of formulas and symbol definitions are provided in the Supplementary information.

### gmMAP-pseudotime: pseudotime-based inference of dynamic cellular developmental metabolic patterns

To characterize dynamic metabolite programs during inflammatory fibroblast differentiation, we developed gmMAP-pseudotime, a trajectory-based extension of gmMAP that integrates genetically informed metabolite scores with single-cell pseudotime. Fibroblast subpopulations were first embedded in a low-dimensional manifold and ordered along an inflammatory trajectory using single-cell trajectory inference. In this implementation, the pseudotime vector was obtained from the ptime field of a scTour-derived AnnData object, and the analysis was focused on a CXCL1-associated fibroblast lineage. scTour was selected because it jointly infers pseudotime, vector-field structure and latent cellular dynamics from single-cell transcriptomes [124].

For each metabolite trait, gmMAP-derived positive and negative metabolite-associated gene programs were scored in each cell using a rank-based UCell strategy. UCell computes robust single-cell gene-signature scores based on the Mann–Whitney U statistic and is well suited for sparse and heterogeneous single-cell data [125]. A signed metabolite score was then calculated for each cell as the difference between the positive and negative metabolite-program scores: *S_i,k_* = UCell(*G_k_*^+^) − UCell(*G_k_*^−^), where *S_i,k_* is the signed gmMAP-pseudotime score for cell *i* and metabolite trait *k*, *G_k_^+^* denotes the positive metabolite-associated gene set, and *G_k_^-^* denotes the negative metabolite-associated gene set.

Cells were retained if they had finite pseudotime values and available metabolite scores. For the CXCL1-associated trajectory, analysis was restricted to WNT2B⁺ Fos-hi fibroblasts, WNT2B⁺ Fos-low fibroblast subsets, CD74⁺ antigen-presenting fibroblasts and CXCL1⁺ fibroblasts. Metabolite identifiers were converted to reported metabolite-trait names using the trait annotation table, and requested metabolites were matched against the signed UCell score matrix. The uploaded plotting implementation reads the scTour AnnData object and signed UCell matrix, uses celltypes and ptime as cell-state and pseudotime annotations, restricts the analysis to the CXCL1 lineage, computes Spearman correlations between pseudotime and metabolite scores, and visualizes smoothed metabolite dynamics along pseudotime.

For each metabolite trait, the association between signed gmMAP score and pseudotime was quantified using Spearman correlation. Scatter plots were colored by fibroblast subtype, and smoothed trends were estimated using LOWESS smoothing when available; otherwise, rolling median smoothing was used. Nominal P values from Spearman correlation were reported in the figure. For visualization of selected metabolite programs, single-cell metabolite scores were plotted against pseudotime, and the fitted trajectory was overlaid to summarize the direction and magnitude of metabolite remodeling during inflammatory fibroblast differentiation.

### Pseudotime analysis of metabolite-associated transcriptional programs

Metabolite-associated transcriptional dynamics were quantified using a signed UCell framework. For each metabolite trait, genes were ranked according to GWAS-derived gene-level Z-scores. The top positively and negatively associated genes were used to construct directional metabolite signatures, retaining traits with at least 100 matched genes in either direction. For each cell, rank-based UCell enrichment scores were computed separately for the positive and negative signatures. A signed metabolite score was defined as the positive-signature UCell score minus the negative-signature UCell score, with negative values truncated to zero.

To identify metabolite-associated programs coupled to nephron differentiation, cells were stratified into predefined developmental lineages and ordered by precomputed Monocle pseudotime. Pseudotime was rescaled to 0–1 within each lineage. Spearman correlation was then used to associate each metabolite signed UCell score with pseudotime, followed by Benjamini–Hochberg correction across metabolites within each lineage. Positively correlated metabolites were interpreted as programs increasing during differentiation, whereas negatively correlated metabolites represented programs decreasing along pseudotime. Selected metabolites were visualized by plotting signed UCell scores against pseudotime, with cells colored by developmental cell type and LOWESS curves indicating smoothed temporal trends.

### gmMAP-perturb: a trajectory-aware framework for metabolite-associated fate rerouting analysis

To investigate whether metabolite-associated transcriptional programs were linked to cellular state transitions, we developed gmMAP-perturb, a trajectory-aware perturbation framework built upon genetically informed metabolite programs inferred by gmMAP. For each metabolite trait m, two complementary gene programs were derived from the corresponding GWAS-to-gene association profile: an activation-associated program (UP genes) and an inhibition-associated program (DOWN genes). Cell-level program activities were quantified using weighted UCell enrichment scores and subsequently standardized across cells.

For each cell c, three metabolite-associated scores were defined:

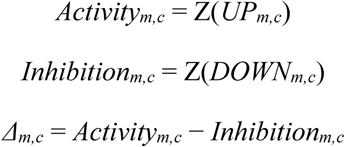

where Z(·) denotes cell-level z-score normalization. Positive Δ values indicate predominance of the activation-associated program, whereas negative values indicate predominance of the inhibition-associated program.

To evaluate the potential effect of metabolite-associated perturbation on lineage specification, cells were projected onto predefined developmental trajectories using pseudotime coordinates obtained from Monocle3. For each lineage bifurcation, a root state, intermediate states and two terminal fates were specified.

Pseudotime values were rescaled to the interval [0,1] and cells belonging to the corresponding trajectory were retained for downstream analysis.

For a given bifurcation, a lineage preference score (FateBias) was calculated as the difference between terminal-state memberships:

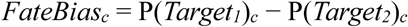

where positive values indicate preferential association with Target1 and negative values indicate preferential association with Target2.

The relationship between metabolite-associated states and lineage preference was quantified using Spearman correlation:

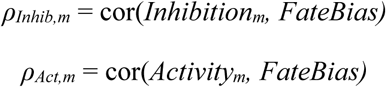

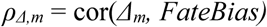

P values were calculated using asymptotic Spearman tests and adjusted across metabolites using the Benjamini-Hochberg procedure.

To identify metabolite programs potentially associated with fate rerouting, branch-specific score dynamics were examined along pseudotime. For each metabolite, terminal branch differences were estimated from the late pseudotime segments of both trajectories. Metabolites were classified according to the concordance between activation-associated, inhibition-associated and Δ-associated signals. Bidirectional-supported perturbations were defined as metabolites for which inhibition-associated programs preferentially accumulated in one terminal lineage while activation-associated or Δ-associated programs preferentially accumulated in the opposite lineage. Single-direction-supported perturbations were defined when only inhibition-associated programs showed significant lineage separation.

For visualization, metabolite-associated scores were projected onto UMAP embeddings and pseudotime trajectories. The original developmental trajectory was represented by the observed lineage path, whereas the predicted perturbation trajectory was represented by the lineage favored by the inhibition-associated program. Importantly, gmMAP-perturb does not simulate biochemical depletion of metabolites or reconstruct new developmental trajectories. Instead, it estimates how metabolite-associated activation and inhibition transcriptional states are associated with lineage bias along existing developmental manifolds.

### gmMAP-drug: drug perturbation reference construction and CMap connectivity analysis

Drug perturbation signatures were generated from publicly available or custom drug-treated transcriptomic datasets. For each drug perturbation, differential expression was calculated by comparing drug-treated samples with matched controls. The resulting signed statistic, preferentially the Wald statistic, moderated t statistic or equivalent signed z statistic, was used as the drug-induced transcriptional signature. For mouse datasets, mouse gene symbols were converted to human orthologues before downstream analysis.

For each metabolite trait, genes were ranked by signed gene-level association scores. The top positively associated genes and bottom negatively associated genes were selected as the metabolite up- and down-signature genes, respectively. CMap connectivity between a drug and a metabolite was calculated as the difference between the mean drug-induced statistic of metabolite up-signature genes and that of metabolite down-signature genes. Positive CMap scores indicated that the drug transcriptional response mimicked or reinforced the metabolite-associated state, whereas negative scores indicated that the drug response reversed or suppressed the metabolite-associated state. Drug-level scores were summarized across available perturbation signatures using the median connectivity score.

### Prediction of tumor-metabolic reversing drugs by gmMAP-drug

For each cancer or epithelial comparison, differential metabolite activity was calculated between tumor and matched adjacent epithelial states. Metabolites with positive differences were interpreted as tumor-enriched metabolic states, whereas negative differences indicated adjacent-enriched states. To prioritize candidate reversing drugs, the vector of tumor-associated metabolite effects was compared with each drug–metabolite CMap connectivity profile. Drugs predicted to negatively connect with tumor-enriched metabolites and positively connect with adjacent-enriched metabolites received higher reversal scores. Drug candidates were ranked by this reversal score, and top-ranked drugs were visualized for each epithelial cancer comparison. For visualization and filtering, module correlations, module composition, epithelial-state activity, drug–module enrichment, differential metabolite volcano plots, drug-reversal rankings and drug–metabolite connectivity bar plots were visualized using custom Python scripts. Unknown metabolites labelled with X- were excluded from primary visualizations. Level traits and ratio traits were analysed separately, because ratio traits cannot be directly interpreted as a unidirectional increase or decrease in a single metabolite. For each drug, the top negative-connected and top positive-connected metabolites were displayed as bilateral bar plots. In these plots, negative CMap connections represent predicted reversal of the metabolite-associated transcriptional state, whereas positive connections represent predicted mimicry or reinforcement.

### Pan-cancer epithelial metabolic program inference

Pan-cancer epithelial metabolic states were inferred from a drug–metabolite and tumor–metabolite connectivity framework. Briefly, epithelial cells from multiple tumor types and clinical epithelial states were analysed using a genetically informed metabolite-mapping strategy. For each metabolite trait, gene-level association scores were used to construct directional metabolite-associated gene signatures. Cell or group level metabolite activities were summarized as signed association effects, hereafter referred to as final effects. Positive final effects indicated enrichment of the metabolite-associated up-signature, whereas negative final effects indicated enrichment of the opposite transcriptional state.

To identify recurrent epithelial metabolic programs, we applied consensus non-negative matrix factorization to the pan-cancer epithelial final-effect matrix. The resulting cNMF components were interpreted as metabolite modules. Module-defining metabolites were ranked by their loading weights, and module activity was quantified across cancer types and epithelial group status, including adjacent, pre-lesion, tumor, metastasis, blood and pleural-fluid states. Pairwise correlations among module-level final-effect profiles were computed to evaluate the similarity and separation of inferred metabolic programs.

### Metabolite–cell type association matrix construction

We analyzed metabolite-associated cell type profiles using the tissue-level summary statistics generated by the upstream scMRS workflow. Unless otherwise specified, the analysis used the wAUCell-derived scores at the S3 normalization stage. For each celltype–metabolite pair, the signed Final effect was used as the primary association measure. Positive values indicated up-dominant metabolite association, whereas negative values indicated down-dominant association. When duplicated records were present for the same celltype and metabolite display name, the entry with the largest absolute final effect was retained.

### Hierarchical clustering of cell types and metabolite features

A celltype-by-metabolite matrix was constructed using signed final effect values. Missing or non-finite values were safely imputed for clustering, and features were standardized before distance calculation. Cell types and metabolite features were independently clustered using hierarchical clustering with average linkage and correlation distance, defined as 1−r, where r is the Pearson correlation coefficient. The final visualization used eight cell modules and seven metabolite feature clusters. Cluster membership was assigned by cutting the dendrogram at the specified number of clusters.

### Identification of directionally significant associations

Directionally significant celltype–metabolite associations were defined using both statistical significance and signed effect size. FDR values were prioritized from available q-value columns; when no q-value was available, Benjamini–Hochberg correction was applied to the corresponding p values within each metabolite trait. An empirical effect-size threshold was estimated from a background set of weakly significant associations, defined by q > 0.20. The threshold was calculated as the 95th percentile of the absolute final effect distribution in this background set, rounded up to the nearest 0.01 and constrained between 0.01 and 0.20. Positive associations were defined as q ≤ 0.05 and AUROC effect ≥ threshold, whereas negative associations were defined as q ≤ 0.05 and AUROC effect ≤ −threshold.

### Feature correlation network

To summarize the internal structure of metabolite feature clusters, pairwise metabolite–metabolite correlations were calculated across cell types using the signed final effect profiles. Within each feature cluster, a hub score was defined as the median absolute correlation between a metabolite and all other metabolites in the same cluster. Features with hub score ≥ 0.40 were retained as representative nodes; if no feature passed this threshold in a cluster, the highest-ranking features were retained as fallback representatives. Network edges were drawn for metabolite pairs with absolute correlation ≥ 0.30 within the same cluster or ≥ 0.55 across different clusters. Node size reflected similarity between each metabolite profile and the centroid profile of its feature cluster.

### Single-cell transcriptomic profiling of fibroblasts in ulcerative colitis

To test the application of gmMAP in inflammatory diseases, single-cell RNA-seq data from colonic tissues were processed using a standard quality-control and normalization workflow. Low-quality cells and genes detected in very few cells were removed. Gene expression matrices were library-size normalized, log-transformed and scaled before dimensionality reduction. Highly variable genes were used for principal-component analysis, followed by neighborhood graph construction, clustering and UMAP visualization. Fibroblast populations were subsetted and reclustered to resolve major fibroblast states. Fibroblast subtypes were annotated according to canonical marker expression, including CD74⁺ antigen-presenting fibroblasts, WNT2B⁺ fibroblast states, WNT5⁺ fibroblasts, myofibroblasts and CXCL1⁺ inflammatory fibroblasts. CXCL1 expression was summarized across healthy, inflamed and non-inflamed tissue groups to evaluate disease-state specificity.

### Correlation between metabolite scores and CXCL1 expression

To evaluate whether gmMAP-inferred metabolic programs were coupled to CXCL1 activation, representative metabolite scores were correlated with CXCL1 gene expression across fibroblast cells. For each selected metabolite or metabolite ratio, normalized CXCL1 expression was regressed against the corresponding gmMAP metabolite score. Pearson correlation coefficients, nominal P values, coefficient of determination, mean absolute error and root mean squared error were calculated. Density scatter plots were generated with a fitted regression line, a 95% prediction band and a 1:1 reference line.

### Pseudotime reconstruction of fibroblast differentiation

Colonic fibroblasts were subsetted from the single-cell transcriptomic dataset and reclustered to resolve major fibroblast states, including CD74⁺ antigen-presenting fibroblasts, WNT2B⁺ fibroblast subsets, WNT5⁺ fibroblasts, myofibroblasts and CXCL1⁺ inflammatory fibroblasts. To infer fibroblast state transitions, pseudotime trajectories were reconstructed using two independent approaches: Monocle3 and scTour.

For Monocle3 analysis, the fibroblast expression matrix and metadata were converted into a cell-data-set object. Dimensionality reduction, cell clustering and principal graph learning were performed on the fibroblast subset. The trajectory root was assigned to early or non-inflammatory fibroblast states, and cells were ordered along pseudotime to evaluate progression toward inflammatory fibroblast activation.

For scTour analysis, a neural-network-based latent representation was learned from the fibroblast transcriptome to infer cell-state dynamics and pseudotime ordering. The inferred scTour trajectory was projected onto the UMAP embedding to visualize directional transitions among fibroblast states. Concordance between Monocle3 and scTour was used to identify robust pseudotime-associated fibroblast remodeling patterns.

### Transcriptomic data collection for metabolism-modulating drugs

To construct a reference compendium of metabolism-modulating drug-induced transcriptional signatures, we systematically collected publicly available transcriptomic datasets from the Gene Expression Omnibus (GEO) database. We focused on pharmacological perturbations targeting major metabolic pathways, including insulin sensitization, lipid metabolism, bile-acid signalling, incretin signalling, mitochondrial and inflammatory metabolism, and cardiometabolic disease-associated pathways. The curated drug perturbations included rosiglitazone, obeticholic acid, bezafibrate, metformin, losartan, pioglitazone, simvastatin, BMS-21, GLP-1, lanifibranor, semaglutide and tirzepatide. The collected datasets covered multiple metabolically relevant tissues or cellular systems, including liver, skeletal muscle, visceral adipose tissue, peripheral nerve, peripheral atherosclerotic plaques and INS-1 pancreatic β cells.

For each dataset, processed expression matrices and sample metadata were downloaded from GEO when available. Treatment and control groups were defined according to the original study design and GEO sample annotations. For microarray datasets, normalized expression matrices provided by the original submitters or GEO series matrix files were used. For RNA-seq datasets, processed count, TPM or normalized expression matrices were preferentially used. When multiple treatment doses, time points or tissues were available, each drug–tissue–condition combination was processed independently to preserve context-specific transcriptional responses. Differential expression analysis was then performed between drug-treated and matched control samples to generate signed drug-induced transcriptional signatures. These signatures were subsequently used as the custom drug-reference matrix for gmMAP-drug analysis to evaluate whether metabolism-modulating perturbations could reverse disease- or tumor-associated metabolite programmes inferred by gmMAP.

The final reference collection included rosiglitazone-treated streptozotocin-induced diabetic peripheral nerve samples from GSE11343 [126], obeticholic acid-treated liver samples from GSE138810 [127], bezafibrate-treated streptozotocin mouse visceral adipose tissue and skeletal muscle datasets from GSE179717 and GSE179718 [128], metformin-treated metabolic tissue datasets from GSE237743/GSE237750 [129], simvastatin-, losartan- and pioglitazone-treated human peripheral atherosclerotic plaque datasets from GSE37824, GLP-1 and BMS-21 treated INS-1 cell datasets from GSE102003, semaglutide- and lanifibranor-treated liver datasets from GSE269493 [130], and semaglutide- and tirzepatide-treated high-fat-diet mouse datasets from GSE297183 [131]. Collectively, this curated transcriptomic compendium provided a context-aware reference for linking gmMAP inferred metabolic programmes to candidate metabolism-modulating drugs.

### GEO-derived metabolite-perturbation transcriptomic compendium

To construct a transcriptomic reference compendium for metabolite- and metabolism-related perturbation analysis, we curated publicly available datasets from the Gene Expression Omnibus (GEO) in which cells or metabolic tissue samples were exposed to defined metabolite treatments, metabolite-related compounds or matched vehicle/control conditions. For each GEO series, sample-level metadata and available processed expression matrices were downloaded from GEO and manually inspected to identify treatment status, control groups, experimental models and sample identifiers. A standardized sample annotation table was generated for each dataset to link GEO sample accessions with treatment and control labels.

Expression matrices were harmonized into a unified gene-by-sample format. When multiple tabular expression files were available for a GEO series, the matrix with the most complete sample coverage and genome-wide gene representation was selected as the primary expression matrix. Gene identifiers were converted to gene symbols when necessary, duplicated gene entries were collapsed, and samples were matched to the standardized sample annotation table. Datasets or files with unsupported formats, incomplete expression matrices or unmatched sample annotations were excluded from downstream matrix-based analyses. This standardization procedure generated a curated compendium comprising 44 GEO series, 734 annotated samples and 58 standardized expression matrices. For downstream drug- and metabolite-perturbation analyses, only datasets with matched treatment labels and usable primary expression matrices were retained.

The resulting compendium was used as a context-aware transcriptomic reference to evaluate whether metabolite- or drug-induced transcriptional responses were concordant with, or reversed, gmMAP-inferred metabolic programmes. Differential transcriptional signatures were computed between treated and matched control samples within each dataset, and these perturbation signatures were subsequently compared with gmMAP-derived metabolite-associated gene programmes to prioritize candidate metabolism-modulating perturbations.

### DSS-induced mouse model of ulcerative colitis

Experimental colitis was induced in mice by administration of dextran sulfate sodium (DSS). Briefly, age- and sex-matched mice were randomly assigned to control and DSS-treated groups. For the DSS group, mice were given 2.5% DSS dissolved in sterile drinking water ad libitum for 6 consecutive days. The DSS solution was freshly prepared and replaced regularly during the treatment period. Control mice received sterile drinking water without DSS under the same housing conditions. During DSS treatment, body weight, stool consistency and rectal bleeding were monitored daily to evaluate disease progression. At the experimental endpoint, mice were euthanized according to approved institutional animal care guidelines. The entire colon was excised, measured for colon length, gently flushed with cold PBS and processed for downstream analyses. Colon tissues were either fixed in 4% paraformaldehyde for histological and immunofluorescence staining or snap-frozen for RNA and protein extraction. Disease severity was assessed based on body-weight loss, clinical disease activity, colon shortening and histopathological inflammation.

### Bile acid sequestrants for the treatment of colitis model

To establish an acute colitis model, mice were administered drinking water containing 2.5% dextran sulfate sodium (DSS) continuously during the experimental period. For bile acid depletion intervention, cholestyramine (300 mg/kg) was delivered via once-daily intragastric gavage to block intestinal bile acid reabsorption. As a classic bile acid sequestrant, cholestyramine non-covalently binds intestinal endogenous bile acids and gut microbiota-derived secondary bile acid metabolites, particularly deoxycholic acid (DCA). This binding effect interrupts bile acid enterohepatic circulation and promotes the fecal excretion of bile acid metabolites, thereby reducing intestinal bile acid and DCA accumulation in vivo. Mice were randomly assigned into three groups: the blank control group (normal drinking water and daily gavage of vehicle), the DSS model group (2.5% DSS drinking water combined with daily vehicle gavage), and the DSS + cholestyramine intervention group (2.5% DSS drinking water with once-daily cholestyramine gavage). At the end of the experiment, all mice were euthanized, and colon tissues were isolated. Colon length was measured to evaluate the degree of colitis-associated pathological injury.

### Metabolite treatment of primary human colonic fibroblasts and Quantitative PCR

Primary human colonic fibroblasts were maintained in fibroblast growth medium supplemented with fetal bovine serum and antibiotics under standard culture conditions at 37°C with 5% CO₂. For metabolite stimulation experiments, cells were seeded into culture plates and allowed to adhere overnight. When cells reached approximately 70–80% confluence, the culture medium was replaced with fresh medium containing vehicle control, 100 μM indoxyl sulfate potassium salt, 100 μM p-cresyl sulfate, or 100 μM deoxycholic acid. Metabolites were prepared as concentrated stock solutions in an appropriate solvent according to their solubility, and the same final vehicle concentration was used across all treatment groups. Cells were incubated with metabolites for 24 h. After treatment, total RNA was extracted from fibroblasts using a standard RNA extraction reagent or column-based RNA isolation kit according to the manufacturer’s instructions. RNA concentration and purity were assessed, and equal amounts of RNA were reverse-transcribed into cDNA. Quantitative PCR was performed to measure CXCL1 mRNA expression, with GAPDH used as the internal reference gene. Relative gene expression was calculated using the 2^−ΔΔCt method and normalized to the vehicle control group. Three independent biological replicates were included for each condition. Data were presented as mean ± SD, and statistical significance was assessed by one-way ANOVA followed by Tukey’s multiple comparisons test.

### Immunofluorescence and in vitro perturbation assays of the TNF-α/HIF1A/lactate/CXCL1 regulatory axis in colonic fibroblasts

In in vitro experiments, primary colonic fibroblasts were exposed to lactate (30 mM), TNF-α (20ng/ml), or a combined treatment of TNF-α and lactate. The mRNA expression of CXCL1 and HIF1A was quantified via quantitative real-time PCR (qPCR), and all data were normalized against the untreated control.

To clarify whether glycolysis and lactate metabolism mediate TNF α triggered inflammatory chemokine expression, we employed two classic glycolytic inhibitors for intervention. TNFα stimulated fibroblasts were separately incubated with 10 mM 2-deoxyglucose (2-DG) or 20 mM sodium oxamate (SO) to suppress glycolytic flux and lactate generation. qPCR was then applied to measure CXCL1 transcript levels and evaluate the inhibitory efficacy of the above treatments.

In vivo experiments were performed using tissue samples from control mice and mice with DSS-induced colitis. Immunofluorescence staining for DAPI, PDPN and HIF1A was carried out on colon tissue sections to characterize stromal HIF1A activation during intestinal inflammation.

### ATAC-seq library preparation and data analysis

Primary mouse colonic fibroblasts were treated with vehicle control, TNF-α (20 ng/ml), or combined TNF-α and lactate stimulation. After treatment, cells were harvested for ATAC-seq to profile chromatin accessibility. Briefly, nuclei were isolated from each sample and subjected to Tn5 transposase-mediated fragmentation. Sequencing libraries were amplified, purified and sequenced on an Illumina platform to generate paired-end reads. Raw sequencing reads were first assessed for quality and adaptor contamination, followed by adaptor trimming and removal of low-quality reads. Clean reads were aligned to the mouse reference genome using Bowtie2. PCR duplicates, mitochondrial reads, unmapped reads and low-quality alignments were removed before downstream analysis. Accessible chromatin regions were identified by peak calling with MACS2. Genome-wide normalized ATAC-seq signal tracks were generated and visualized around the Cxcl1 promoter region. Average accessibility profiles and heatmaps were plotted within ±5 kb of the Cxcl1 promoter center to compare chromatin accessibility among control, TNF-α-treated and TNF-α plus lactate-treated fibroblasts.

### Sodium oxamate intervention in the DSS-induced colitis model

To establish an acute colitis model, mice were administered drinking water containing 2.5% dextran sulfate sodium (DSS) continuously for 6 days. For pharmacological inhibition of lactate dehydrogenase activity, sodium oxamate was used as an LDH inhibitor to suppress pyruvate-to-lactate conversion and reduce lactate-associated metabolic signalling during intestinal inflammation. Sodium oxamate was freshly prepared in sterile PBS and administered once daily by intraperitoneal injection at 300 mg/kg during the DSS modelling period.

Mice were randomly assigned into three groups: the blank control group, receiving normal drinking water and daily PBS injection; the DSS model group, receiving 2.5% DSS drinking water and daily PBS injection; and the DSS + sodium oxamate intervention group, receiving 2.5% DSS drinking water combined with daily sodium oxamate treatment. Body weight, stool consistency and rectal bleeding were monitored during the experiment to evaluate disease activity. At the end of the experiment, mice were euthanized, and colon tissues were isolated. Colon length was measured as an indicator of colitis-associated pathological injury, and colon samples were collected for subsequent histological, molecular and inflammatory analyses.

### TSA-based multiplex immunofluorescence staining of PDPN and HIF1A in DSS-treated mouse intestinal tissues

Colon tissues were collected from control and DSS-treated mice, fixed, dehydrated, embedded in paraffin, and sectioned at 4–5 μm thickness. Sections were deparaffinized, subjected to antigen retrieval, permeabilized, and blocked with 3% H₂O₂ to quench endogenous peroxidase activity. Sequential TSA staining was performed using the tyramide signal amplification system. Primary antibodies against PDPN and HIF1A were applied sequentially overnight at 4°C, followed by incubation with HRP-conjugated secondary antibodies for 1 h at room temperature. Fluorophore-conjugated tyramide substrates with distinct emission spectra were used for signal development. Antibody stripping was performed between rounds to remove primary/secondary antibody complexes, eliminating cross-reactivity even when using primary antibodies from the same host species. After completion of all staining rounds, nuclei were counterstained with DAPI, and sections were mounted with antifade medium. Fluorescence images were acquired using a confocal microscope or fluorescence slide scanner under identical exposure conditions. PDPN signal was used to identify stromal and lymphatic-associated structures, while HIF1A expression was evaluated by nuclear/perinuclear fluorescence intensity. Quantitative analysis of PDPN and HIF1A fluorescence intensity, percentage of HIF1A-positive cells, and their spatial colocalization was performed using ImageJ in comparable mucosal and submucosal regions. Negative controls without primary antibodies were included to assess nonspecific staining.

### Development of Metabolism-AI agent to resolve cellular metabolic networks

We developed Metabolism-AI, a literature-guided metabolic knowledge agent for structured interpretation of tumor metabolite-associated outputs from GCST/trait differential tables, ordinary metabolite tables, single-cell annotations, differential genes, pathway activity scores and metabolite-ranking results. To construct the knowledge base, we assembled a tumor-metabolism literature corpus from PubMed and locally curated full-text PDFs covering publications from January 2015 to May 2026. The search combined cancer-related terms (cancer, tumor/tumor, neoplasm, carcinoma, malignancy, oncology and Neoplasms MeSH) with metabolism-related terms (metabolism, metabolic, metabolite, metabolomics, metabolome, cancer/tumor metabolism, metabolic reprogramming, metabolic rewiring, metabolic plasticity, energy metabolism, bioenergetics, Metabolism MeSH and Metabolomics MeSH), retrieving approximately 140,000 records; a journal impact-factor filter (IF > 5) retained approximately 50,000 articles as the core interpretive corpus. The agent used a dual-model framework combining a general large language model and DeepSeek-based reasoning modules to perform entity recognition, synonym normalization, GCST-to-trait mapping, metabolite ratio parsing, evidence retrieval and rule-constrained mechanistic reasoning. Extracted entities, including metabolites, genes, enzymes, pathways, tumor types, cell types and metabolic phenotypes, were organized into a local knowledge graph linking metabolite features to pathway annotations, candidate regulatory genes and literature-supported biological processes. GCST-only inputs required a local annotation table to map accessions to reported traits and candidate metabolites; metabolite ratios were decomposed into numerator and denominator components and flagged when directional interpretation was ambiguous. For tumor metabolic interpretation, the agent mapped cell-type- or tumor-state-specific metabolite features to pathways and candidate genes, prioritized mechanisms such as lipid biosynthesis, fatty-acid oxidation, glycolysis, glutaminolysis, TCA-cycle activity, branched-chain amino-acid metabolism, bile-acid metabolism, sphingolipid metabolism, redox metabolism and tumor microenvironment-associated metabolic rewiring, and assigned confidence tiers based on annotation consistency, literature support, overlap with upstream differential or single-cell results, cell-type specificity and biological coherence. Outputs included normalized metabolite tables, metabolite–pathway mappings, ranked pathway summaries, candidate gene lists, pathway–gene networks, evidence tables and concise interpretation reports, with ambiguous metabolites, weak evidence, missing GCST annotations and association-not-causation boundaries explicitly flagged. The system was used only for biological interpretation and validation prioritization, not for clinical decision-making. Source code is available at https://github.com/chenms2000/meta_down_analysis and archived as release v1.0.0-paper under the MIT License, whereas annotation tables, upstream data, third-party databases and literature-derived resources are managed as separate data artifacts subject to their original licenses.

### Disclosure of artificial intelligence tools use

ChatGPT was used to assist with formatting and refinement of code used in this study, as well as linguistic polishing of manuscript text and figure legends. The authors took full responsibility for reviewing, adjusting and finalizing all outputs generated by the AI assistant to ensure scientific accuracy and logical rigor.

## Supporting information

All result

## Code availability

A complete Python package integrating the core gmMap algorithm and five complementary sub-tools has been developed, with all source code released openly at https://github.com/Xulab-collab/gmMAP. The repository contains the core gmMAP workflow for constructing genetically informed metabolite-associated gene programs, scoring single-cell and spatial transcriptomic datasets, identifying cell-type-specific metabolite associations, and running the downstream application modules described in this study, including gmMAP-spatial, gmMAP-pseudotime, gmMAP-flux, gmMAP-perturb and gmMAP-drug. Example configuration files and command-line usage are provided in the repository to facilitate reproducible analysis. The released code does not include restricted GWAS summary statistics, unpublished single-cell or spatial transcriptomic datasets, intermediate analysis results, or private annotation tables.

We have developed an interactive web tool (https://immunotcm.ac.cn/gmMAP/) for visualizing and downloading spatial metabolite mapping profiles across multiple mouse tissues and their corresponding trait-tissue association maps.

The code of Metabolism-AI Agent is available at https://github.com/chenms2000/meta_down_analysis and archived as release v1.0.0-paper. Code and documentation are released under the MIT License; real annotation tables, upstream data, third-party databases, and literature-derived resources are managed as separate data artifacts subject to their original licenses and citation requirements.

## Data availability

Published public datasets: All datasets analysed in this study are publicly available from the repositories associated with the original publications. Metabolite GWAS summary statistics were retrieved from plasma metabolome GWAS datasets deposited in the GWAS Catalog (accession IDs: GCST90200448–GCST90203010). Processed single-cell RNA-sequencing matrices and annotations were obtained for the Tabula Sapiens human atlas (GSE201333), human pan-cancer single-cell atlas (https://zenodo.org/records/15169362), spatial metabolomic atlas of human kidney development (E-MTAB-11429), single-cell RNA-sequencing atlas of human kidney development (GSE114530), spatial transcriptomic data of whole-body mouse under healthy and LPS-challenged conditions (GSE248904), single-cell transcriptomic data of human ulcerative colitis (SCP259), 10x Visium spatial transcriptomic data of DSS-induced colitis mouse model (GSE169749), human brain atlas (Human Brain Cell Atlas v1.0), Multi-subtype pediatric high-grade glioma atlas (https://cellxgene.cziscience.com/collections/9ceda3d2-a55e-4b82-a91a-0190d947459a).

Public drug-perturbation transcriptomic datasets were curated from the Gene Expression Omnibus, including GSE11343, GSE138810, GSE179717, GSE179718, GSE237743, GSE237750, GSE37824, GSE102003, GSE269493 and GSE297183. These datasets covered rosiglitazone, obeticholic acid, bezafibrate, metformin, simvastatin, losartan, pioglitazone, GLP-1, BMS-21, semaglutide, lanifibranor and tirzepatide perturbations across metabolic tissues, vascular lesions and cellular models, and were used to construct a context-aware reference for linking gmMAP-inferred metabolic programmes to candidate metabolism-modulating drugs. Public metabolite-perturbation transcriptomic datasets were curated from the Gene Expression Omnibus to construct a context-aware reference compendium for gmMAP-based perturbation analysis. The curated collection comprised 44 GEO series and 734 annotated samples, including GSE173611, GSE129756, GSE216833, GSE210692, GSE51834, GSE132410, GSE185598, GSE263024, GSE155325, GSE155326, GSE161342, GSE165737, GSE160307, GSE140269, GSE207077, GSE115354, GSE267694, GSE37852, GSE242872, GSE214587, GSE131754, GSE131901, GSE144398, GSE214153, GSE168928, GSE148143, GSE200309, GSE248578, GSE180948, GSE219145, GSE274169, GSE243083, GSE164806, GSE161730, GSE164740, GSE155715, GSE212738, GSE238241, GSE184821, GSE231988, GSE118230, GSE302368, GSE235357 and GSE4817.

Data produced in this study: The spatial metabolomic, spatial transcriptomic, and single-cell RNA-seq datasets derived from high-grade glioma specimens in this study have been deposited in the Zenodo open-access repository and can be accessed via the DOI: 10.5281/zenodo.20816032. The transcriptome data generated from lactic acid-stimulated colonic fibroblasts can be accessed on the Zenodo repository with the DOI: 10.5281/zenodo.20816386. Single-cell RNA-seq data of colon tissues from healthy control mice and DSS-induced colitis mice can be accessed on the Zenodo repository with the DOI: https://10.5281/zenodo.20849018.

## Acknowledgements

We gratefully acknowledge the investigators, consortia and data contributors who generated and made publicly available the single-cell transcriptomic, spatial transcriptomic and spatial metabolomic datasets used in this study. In particular, we thank the Tabula Sapiens Consortium for providing the human multi-tissue single-cell atlas; the contributors of the pan-cancer single-cell atlas, human kidney developmental atlas, human brain single-cell atlas, pediatric high-grade glioma atlas and ulcerative colitis colon atlas for releasing processed expression matrices and cell annotations; and the investigators who generated the whole-body mouse spatial transcriptomic datasets under steady-state and inflammatory conditions. We also acknowledge the researchers who produced the spatial metabolomics datasets of human kidney development, which enabled independent evaluation of gmMAP-derived metabolite predictions. These openly available resources were essential for developing, benchmarking and applying gmMAP across human tissues, developmental contexts, inflammatory disease and cancer.

## Author Contributions

A.X., H.X. and G.H. conceived and designed the study. H.X. developed the gmMAP framework, implemented the main computational workflow and completed most of the bioinformatics analyses. H.X., L.Z. and W.L. developed and optimized the downstream analytical modules, including metabolite-program scoring, cell-type association analysis, spatial mapping, pseudotime analysis, metabolic-flow potential inference, perturbation prediction and drug-reversal analysis. G.H., L.Z. and W.L. contributed to data collection, data preprocessing, benchmarking analyses and biological interpretation of the results. Q.W., M.C., D.Z. and Y.Z. contributed to data organization, visualization, literature curation and validation analysis. H.X. prepared the figures and wrote the initial manuscript draft. G.H., L.Z., W.L. and A.X. provided critical comments and revised the manuscript. A.X. supervised the project and provided funding support. All authors discussed the results, provided feedback and approved the final manuscript. H.X., G.H., L.Z. and W.L. contributed equally to this work.

## Ethic

All animal experiments were approved by the Institutional Animal Care and Use Committee of Beijing University of Chinese Medicine (Approval No. BUCM-2025040106-2094) and were conducted in accordance with the relevant institutional guidelines and regulations. All human glioma clinical specimens were collected in accordance with the ethical approval granted by the Institutional Review Board (IRB) of Sanbo Brain Hospital, Capital Medical University (Approval No. SBNK-Y.JYS-2025-042-01).

