## Supplementary material for "Genetically informed single-cell and spatial mapping of metabolic programs in human health and disease": All result

#### Comparison of gmMAP, Compass, scFEA and scMetabolism

**Supplementary table S1**. Comparison of gmMAP with representative single-cell metabolism analysis methods. Boxes shaded in light orange represent biological functions with superior performance predicted by gmMAP

| Feature | Compass | scFEA | scMetabolism | gmMAP |
|---|---|---|---|---|
| Prevent transcript-protein-enzyme activity mismatch | ✗ | ✗ | ✗ | ✓ |
| GWAS-informed metabolite traits | ✗ | ✗ | ✗ | ✓ |
| Individual metabolite-cell state associations | ✗ | ✗ | ✗ | ✓ |
| External/microbial metabolite–cell state associations | ✗ | ✗ | ✗ | ✓ |
| Spatial metabolite mapping | ✗ | ✗ | ✗ | ✓ |
| Cell fate-associated metabolites | Limited | Limited | Limited | ✓ |
| Metabolite-perturbation on cell fate | ✗ | ✗ | ✗ | ✓ |
| Flux inference | ✓(Recon3D-based) | ✓ | ✗ | ✓Relative flow potential |
| Algorithm runtime | Slow | Fast | Fast | Fast |
| Metabolite-ratio support | ✗ | ✗ | ✗ | ✓ |
| Reaction-level activity | ✓ | ✓ | ✗ | ✗ |
| Pathway activity | ✓ | ✓ | ✓ | ✓ |
| Hormones and signalling metabolites | Limited | ✗ | ✗ | ✓ |

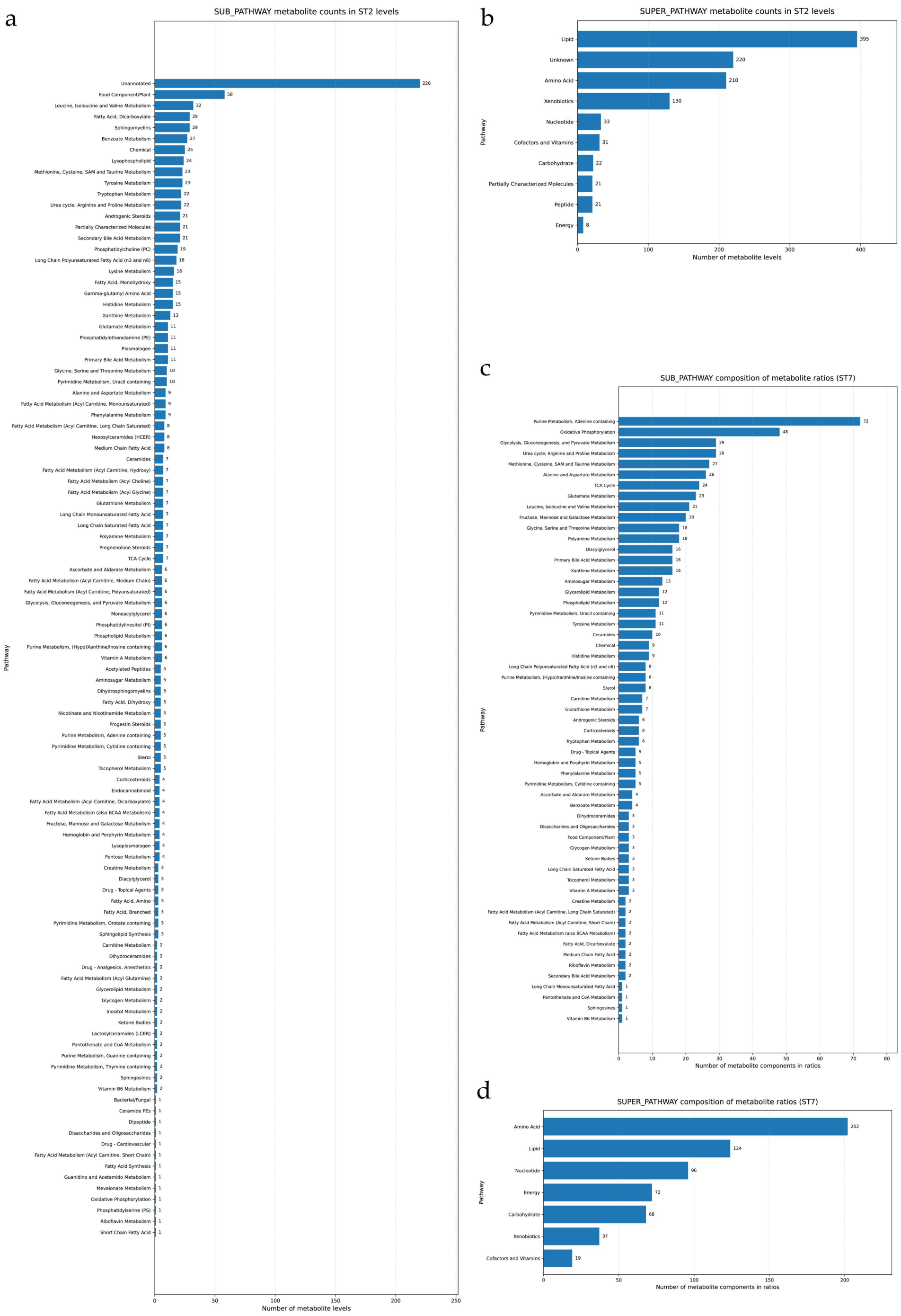


**Supplementary Figure S1. Statistical plot of metabolic pathways of metabolites and metabolic ratios.**

a. Horizontal bar chart showing the total counts of annotated individual metabolite levels grouped at the subpathway level under primary metabolic classification. Unassigned metabolites represent metabolites without definitive subpathway annotation. b. Summary quantification of metabolite-level abundances aggregated into top-tier superpathway categories. c. Subpathway-level statistical distribution of constituent metabolite numbers forming quantified metabolite ratios. Each bar corresponds to the total count of ratio-involved metabolites per individual subpathway. d. Superpathway-level aggregated statistics of metabolite components participating in all detected metabolite ratios.

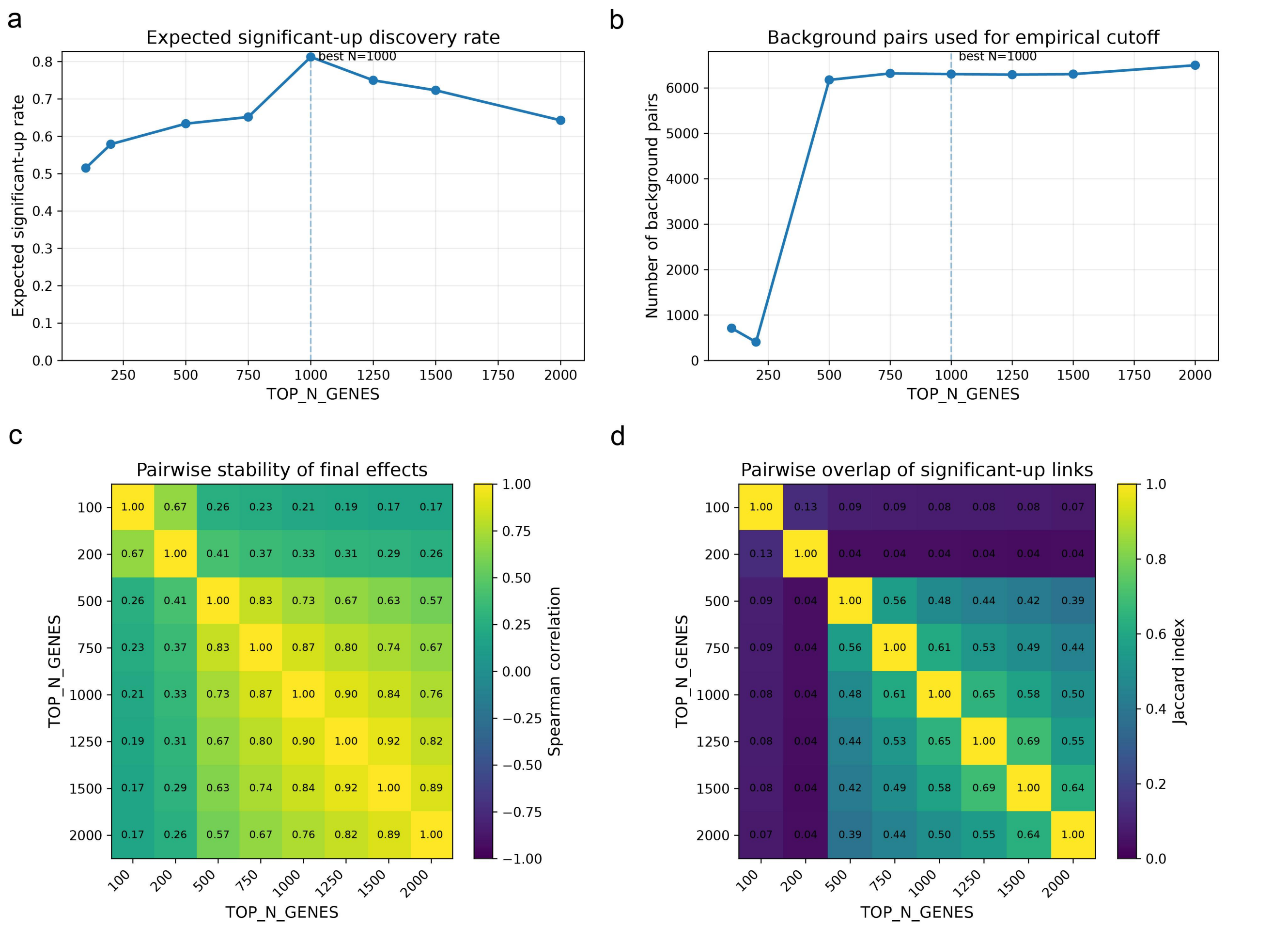


**Supplementary Figure S2. Optimization of metabolite trait gene-set size for gmMAP using paired developmental kidney metabolite and single-cell transcriptomic data.**

a, Expected significant-up discovery rate across different numbers of GWAS-derived trait-associated genes used for each metabolite trait. Significant positive metabolite–cell-type links were defined by $q < 0.05$ and a positive final effect exceeding the empirical effect cutoff estimated from background metabolite–cell-type pairs. The expected discovery rate peaked at TOP_N_GENES = 1000, indicating the strongest recovery of expected cell-type-level metabolic associations.

b, Number of background metabolite–cell-type pairs used for empirical final-effect cutoff estimation across gene-set sizes. Background pairs were defined as non-significant associations with $q > 0.20$. The background set was unstable for very small gene sets but reached a plateau after TOP_N_GENES ≥ 500, supporting more reliable empirical threshold estimation in this range.

c, Pairwise Spearman correlation heatmap of final metabolite–cell-type effects across different gene-set sizes. Larger gene sets from 500 to 2000 showed substantially higher cross-parameter stability, whereas 100 and 200 genes produced less stable effect landscapes.

d, Pairwise Jaccard overlap of significant-up metabolite–cell-type links across gene-set sizes. Positive significant links were more consistently recovered among gene sets from 500 to 2000, with TOP_N_GENES = 1000 showing strong overlap with adjacent settings. Dashed vertical lines in a and b indicate the selected default setting, TOP_N_GENES = 1000.

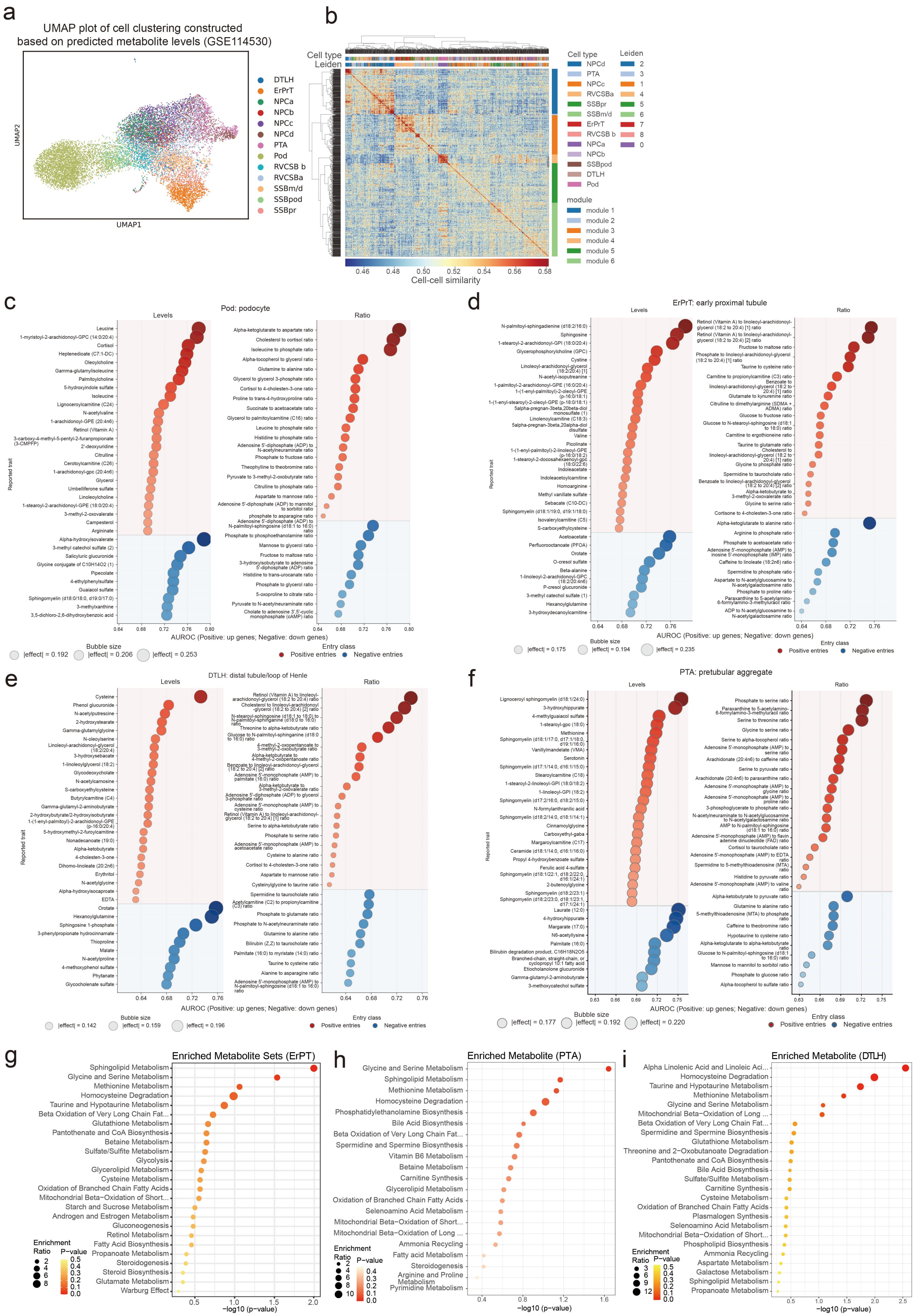
a
UMAP plot of cell clustering constructed based on predicted metabolite levels (GSE114530)
UMAP1
UMAP2
DTLH
ErPrT
NPCa
NPCb
NPCc
NPCd
PTA
Pod
RVCSB b
RVCSBa
SSBm/d
SSBpod
SSBpr
b
Cell type
Leiden
Cell-cell similarity
module
module 1
module 2
module 3
module 4
module 5
module 6
c
Pod: podocyte
Levels
Ratio
Reported trait
AUROC (Positive: up genes; Negative: down genes)
Bubble size
|effect| = 0.192
|effect| = 0.206
|effect| = 0.253
Entry class
Positive entries
Negative entries
d
ErPrT: early proximal tubule
|effect| = 0.175
|effect| = 0.194
|effect| = 0.235
e
DTLH: distal tubule/loop of Henle
|effect| = 0.142
|effect| = 0.159
|effect| = 0.196
f
PTA: pretubular aggregate
|effect| = 0.177
|effect| = 0.192
|effect| = 0.220
g
Enriched Metabolite Sets (ErPT)
Sphingolipid Metabolism
Glycine and Serine Metabolism
Methionine Metabolism
Homocysteine Degradation
Taurine and Hypotaurine Metabolism
Beta Oxidation of Very Long Chain Fat...
Glutathione Metabolism
Pantothenate and CoA Biosynthesis
Betaine Metabolism
Sulfate/Sulfite Metabolism
Glycolysis
Glycerolipid Metabolism
Cysteine Metabolism
Oxidation of Branched Chain Fatty Acids
Mitochondrial Beta-Oxidation of Short...
Starch and Sucrose Metabolism
Androgen and Estrogen Metabolism
Gluconeogenesis
Retinol Metabolism
Fatty Acid Biosynthesis
Propanoate Metabolism
Steroidogenesis
Steroid Biosynthesis
Glutamate Metabolism
Warburg Effect
Enrichment Ratio
P-value
-log10 (p-value)
h
Enriched Metabolite (PTA)
Glycine and Serine Metabolism
Sphingolipid Metabolism
Methionine Metabolism
Homocysteine Degradation
Phosphatidylethanolamine Biosynthesis
Bile Acid Biosynthesis
Beta Oxidation of Very Long Chain Fat...
Spermidine and Spermine Biosynthesis
Vitamin B6 Metabolism
Betaine Metabolism
Carnitine Synthesis
Glycerolipid Metabolism
Oxidation of Branched Chain Fatty Acids
Selenoamino Acid Metabolism
Mitochondrial Beta−Oxidation of Short...
Mitochondrial Beta−Oxidation of Long ...
Ammonia Recycling
Fatty acid Metabolism
Steroidogenesis
Arginine and Proline Metabolism
Pyrimidine Metabolism
i
Enriched Metabolite (DTLH)
Alpha Linolenic Acid and Linoleic Aci...
Homocysteine Degradation
Taurine and Hypotaurine Metabolism
Methionine Metabolism
Glycine and Serine Metabolism
Mitochondrial Beta−Oxidation of Long ...
Beta Oxidation of Very Long Chain Fat...
Spermidine and Spermine Biosynthesis
Glutathione Metabolism
Threonine and 2−Oxobutanoate Degradation
Pantothenate and CoA Biosynthesis
Bile Acid Biosynthesis
Sulfate/Sulfite Metabolism
Carnitine Synthesis
Cysteine Metabolism
Oxidation of Branched Chain Fatty Acids
Plasmalogen Synthesis
Selenoamino Acid Metabolism
Mitochondrial Beta−Oxidation of Short...
Phospholipid Biosynthesis
Ammonia Recycling
Aspartate Metabolism
Galactose Metabolism
Sphingolipid Metabolism
Propanoate Metabolism

**Supplementary Figure S3. gmMAP resolves cell-type-specific metabolite programs across developing nephron lineages.**

a. UMAP visualization of developing kidney cells clustered according to gmMAP-predicted metabolite levels from the single-cell RNA-seq dataset GSE114530. Cells are coloured by annotated renal developmental cell type, showing that metabolite-based predictions preserve major nephron lineage relationships. b. Cell–cell similarity heatmap based on gmMAP-predicted metabolite profiles. Top annotations indicate cell type, Leiden cluster and metabolite module assignment. Predicted metabolite programs separated major renal developmental populations and grouped functionally related cell states. c–f. Ranked metabolite traits predicted by gmMAP for representative renal cell types. For each cell type, metabolite levels and metabolite ratios are shown separately. Bubble position indicates AUROC value, with positive entries reflecting enrichment of up-associated metabolite gene programs and negative entries reflecting enrichment of down-associated programs; bubble size denotes the absolute effect magnitude. c. Podocytes showed enrichment of leucine, cortisol, glycerophosphocholine-related lipids, acylcarnitine-associated metabolites and amino-acid or lipid-related ratios, consistent with nutrient sensing, steroid responsiveness and membrane lipid remodelling. d. Early proximal tubular cells showed prominent lipid, sphingolipid, carnitine/acylcarnitine and amino-acid-associated metabolite features, consistent with mitochondrial fuel metabolism and reabsorptive specialization. e. Distal tubule/loop-of-Henle cells displayed distinct amino-acid, taurine/hypotaurine, lipid and mitochondrial-associated metabolite signatures, suggesting metabolic adaptation during tubular maturation. f. Pretubular aggregate cells were enriched for sphingolipid, phospholipid, glycine/serine, methionine/homocysteine and polyamine-related features, consistent with biosynthetic and membrane-remodelling activity during epithelial commitment. g. Early proximal tubular cells were enriched for sphingolipid metabolism, glycine and serine metabolism, methionine metabolism, homocysteine degradation, taurine/hypotaurine metabolism, glutathione metabolism, fatty-acid metabolism and mitochondrial β-oxidation-related pathways. h. Pretubular aggregate cells showed enrichment of glycine/serine metabolism, sphingolipid metabolism, methionine metabolism, homocysteine degradation, phosphatidylethanolamine biosynthesis, bile-acid biosynthesis and polyamine-related pathways. i. Distal tubule/loop-of-Henle cells were enriched for alpha-linolenic and linoleic acid metabolism, homocysteine degradation, taurine/hypotaurine metabolism, methionine metabolism, mitochondrial β-oxidation and amino-acid metabolic pathways. Node size indicates enrichment ratio and colour indicates enrichment significance.

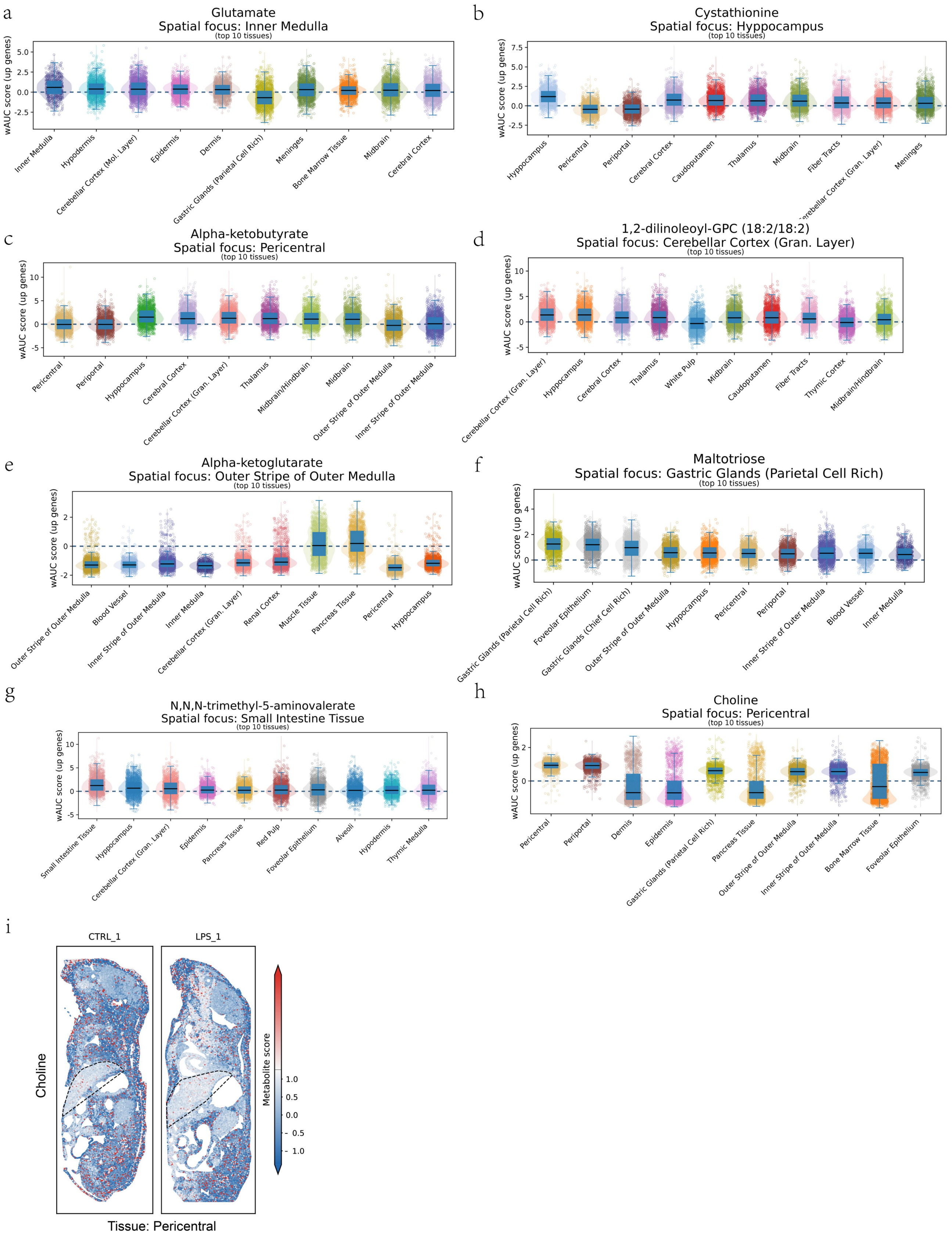


**Supplementary Figure S4. Tissue-level distributions and spatial localization of representative metabolite-associated programs.**

a, Distribution of glutamate-associated wAUCell scores across the top-ranked tissues. The inner medulla was

identified as the primary spatial focus, with additional signals in hypothalamus, cerebellar cortex molecular layer, epidermis, dermis, meninges, midbrain and cerebral cortex, indicating that the glutamate-associated program captures both renal and neural tissue-related metabolic features.
b, Distribution of cystathionine-associated scores across top-ranked tissues. The hippocampus showed the strongest tissue-level focus, with additional enrichment across cerebral cortex, caudoputamen, thalamus, midbrain, fiber tracts, cerebellar cortex granular layer and meninges, consistent with a neural sulfur amino acid and redox-associated metabolic program.
c, Distribution of alpha-ketobutyrate-associated scores across top-ranked tissues. Pericentral tissue was highlighted as the spatial focus, while substantial signals were also observed in hippocampus, cerebral cortex, cerebellar cortex granular layer, thalamus, midbrain and renal medullary regions, suggesting inflammation- and tissue-context-dependent remodeling of amino acid-related metabolism.
d, Distribution of 1,2-dilinoleoyl-GPC (18:2/18:2)-associated scores across top-ranked tissues. The cerebellar cortex granular layer showed the primary tissue focus, with additional signals in hippocampus, cerebral cortex, thalamus, white pulp, midbrain, caudoputamen, fiber tracts and thymic cortex, supporting accurate detection of brain-enriched choline-containing phospholipid-associated programs.
e, Distribution of alpha-ketoglutarate-associated scores across top-ranked tissues. The outer stripe of the outer medulla was identified as the spatial focus, with additional signals across renal medullary regions, blood vessels, cerebellar cortex granular layer, renal cortex, muscle tissue, pancreas tissue, pericentral tissue and hippocampus, consistent with broad mitochondrial and tricarboxylic acid cycle-related metabolic activity.
f, Distribution of maltotriose-associated scores across top-ranked tissues. Gastric glands enriched for parietal cells showed the strongest tissue-level focus, with additional signals in foveolar epithelium, gastric glands enriched for chief cells, renal outer medulla, hippocampus, pericentral and periportal tissues, supporting a gastrointestinal carbohydrate-associated metabolic program.
g, Distribution of N,N,N-trimethyl-5-aminovalerate-associated scores across top-ranked tissues. Small intestine tissue showed the strongest spatial focus, with additional signals in hippocampus, cerebellar cortex granular layer, epidermis, pancreas tissue, red pulp, foveolar epithelium, liver, hypothalamus and thymic medulla, indicating an intestine-associated metabolite program with broader tissue-level variation.
h, Distribution of choline-associated scores across top-ranked tissues. Pericentral tissue was identified as the primary spatial focus, with prominent signals also observed in periportal regions, gastric glands, pancreatic tissue, renal outer medulla, bone marrow tissue and foveolar epithelium, consistent with broad systemic choline metabolism and liver-associated phosphatidylcholine turnover.
i, Representative spatial maps of the choline-associated program in pericentral tissue under control and LPS-treated conditions. The spatial maps show localized choline-associated metabolic scores within the tissue section, illustrating the ability of gmMAP to resolve region-specific metabolite-associated programs and their remodeling under inflammatory perturbation.

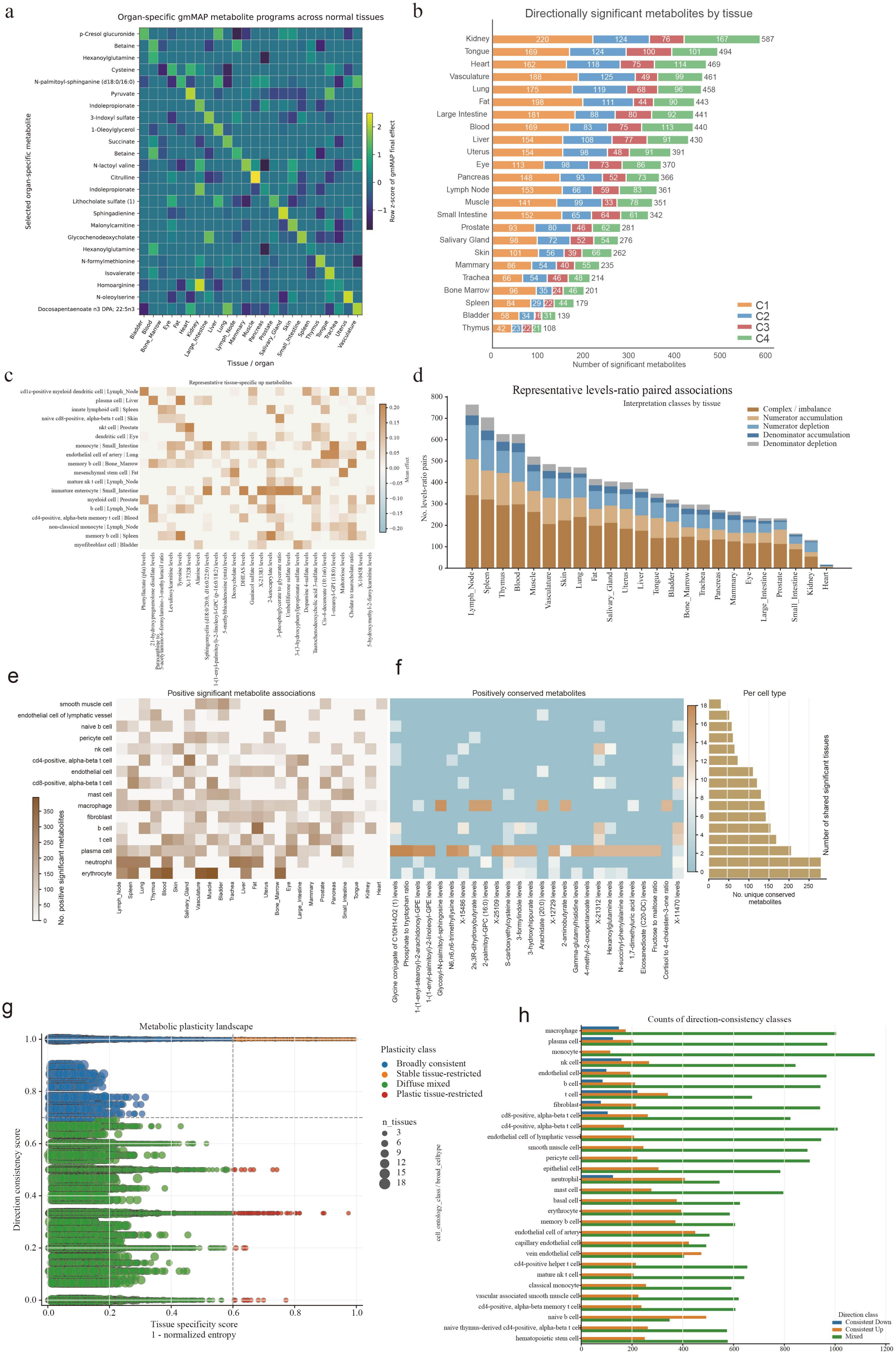
a
Organ-specific gmMAP metabolite programs across normal tissues
Selected organ-specific metabolite
p-Cresol glucuronide
Betaine
Hexanoylglutamine
Cysteine
N-palmitoyl-sphinganine (d18:0/16:0)
Pyruvate
Indolepropionate
3-Indoxyl sulfate
1-Oleoylglycerol
Succinate
Betaine
N-lactoyl valine
Citrulline
Indolepropionate
Lithocholate sulfate (1)
Sphingadienine
Malonylcarnitine
Glycochenodeoxycholate
Hexanoylglutamine
N-formylmethionine
Isovalerate
Homoarginine
N-oleoylserine
Docosapentaenoate n3 DPA; 22:5n3
Bladder
Blood
Bone_Marrow
Eye
Fat
Heart
Kidney
Large_Intestine
Liver
Lung
Lymph_Node
Mammary
Muscle
Pancreas
Prostate
Salivary_Gland
Skin
Small_Intestine
Spleen
Thymus
Tongue
Trachea
Uterus
Vasculature
Tissue / organ
Row z-score of gmMAP final effect
b
Directionally significant metabolites by tissue
Kidney 220 124 76 167 587
Tongue 169 124 100 101 494
Heart 162 118 75 114 469
Vasculature 188 125 49 99 461
Lung 175 119 68 96 458
Fat 198 111 44 90 443
Large Intestine 181 88 80 92 441
Blood 169 83 75 113 440
Liver 154 108 77 91 430
Uterus 154 98 48 91 391
Eye 113 98 73 86 370
Pancreas 148 93 52 73 366
Lymph Node 153 66 59 83 361
Muscle 141 99 33 78 351
Small Intestine 152 65 64 61 342
Prostate 93 80 46 62 281
Salivary Gland 98 72 52 54 276
Skin 101 56 39 66 262
Mammary 86 54 40 55 235
Trachea 66 54 46 48 214
Bone Marrow 96 35 24 46 201
Spleen 84 29 22 44 179
Bladder 58 34 16 31 139
Thymus 42 23 22 21 108
C1
C2
C3
C4
Number of significant metabolites
c
Representative tissue-specific up metabolites
cd1c-positive myeloid dendritic cell | Lymph_Node
plasma cell | Liver
innate lymphoid cell | Spleen
naive cd8-positive, alpha-beta t cell | Skin
nkt cell | Prostate
dendritic cell | Eye
monocyte | Small_Intestine
endothelial cell of artery | Lung
memory b cell | Bone_Marrow
mesenchymal stem cell | Fat
mature nk t cell | Lymph_Node
immature enterocyte | Small_Intestine
myeloid cell | Prostate
b cell | Lymph_Node
cd4-positive, alpha-beta memory t cell | Blood
non-classical monocyte | Lymph_Node
memory b cell | Spleen
myofibroblast cell | Bladder
Mean effect
d
Representative levels-ratio paired associations
Interpretation classes by tissue
Complex / imbalance
Numerator accumulation
Numerator depletion
Denominator accumulation
Denominator depletion
No. levels-ratio pairs
Lymph_Node
Spleen
Thymus
Blood
Muscle
Vasculature
Skin
Lung
Fat
Salivary_Gland
Uterus
Liver
Tongue
Bladder
Bone_Marrow
Trachea
Pancreas
Mammary
Eye
Large_Intestine
Prostate
Small_Intestine
Kidney
Heart
e
Positive significant metabolite associations
smooth muscle cell
endothelial cell of lymphatic vessel
naive b cell
pericyte cell
nk cell
cd4-positive, alpha-beta t cell
endothelial cell
cd8-positive, alpha-beta t cell
mast cell
macrophage
fibroblast
b cell
t cell
plasma cell
neutrophil
erythrocyte
No. positive significant metabolites
f
Positively conserved metabolites
Glycine conjugate of C10H14O2 (1) levels
Phosphate to tryptophan ratio
1-(1-enyl-stearoyl)-2-arachidonoyl-GPE levels
1-(1-enyl-palmitoyl)-2-linoleoyl-GPE levels
Glycosyl-N-palmitoyl-sphingosine levels
N6,n6,n6-trimethyllysine levels
X-15486 levels
2s,3R-dihydroxybutyrate levels
2-palmitoyl-GPC (16:0) levels
X-25109 levels
S-carboxyethylcysteine levels
3-formylindole levels
3-hydroxyhippurate levels
Arachidate (20:0) levels
X-12729 levels
2-aminobutyrate levels
Gamma-glutamylhistidine levels
4-methyl-2-oxopentanoate levels
X-21312 levels
Hexanoylglutamine levels
N-succinyl-phenylalanine levels
1,7-dimethyluric acid levels
Eicosanedioate (C20-DC) levels
Fructose to maltose ratio
Cortisol to 4-cholesten-3-one ratio
X-11470 levels
Per cell type
Number of shared significant tissues
No. unique conserved metabolites
g
Metabolic plasticity landscape
Direction consistency score
Tissue specificity score
1 - normalized entropy
Plasticity class
Broadly consistent
Stable tissue-restricted
Diffuse mixed
Plastic tissue-restricted
n_tissues
h
Counts of direction-consistency classes
cell_ontology_class / broad_celltype
macrophage
plasma cell
monocyte
nk cell
endothelial cell
b cell
t cell
fibroblast
cd8-positive, alpha-beta t cell
cd4-positive, alpha-beta t cell
endothelial cell of lymphatic vessel
smooth muscle cell
pericyte cell
epithelial cell
neutrophil
mast cell
basal cell
erythrocyte
memory b cell
endothelial cell of artery
capillary endothelial cell
vein endothelial cell
cd4-positive helper t cell
mature nk t cell
classical monocyte
vascular associated smooth muscle cell
cd4-positive, alpha-beta memory t cell
naive b cell
naive thymus-derived cd4-positive, alpha-beta t cell
hematopoietic stem cell
Direction class
Consistent Down
Consistent Up
Mixed
No. retained celltype–metabolite pairs

**Supplementary Figure S5. Conserved and tissue-restricted metabolite-associated programmes across organs and cell types.**

**a.** Heatmap of tissue-specific upregulated metabolites. **b.** Number of directionally significant metabolites in each tissue, stratified by metabolite cluster. Kidney showed the largest repertoire of significant metabolite associations, followed by tongue, heart, vasculature, lung, fat, large intestine, blood and liver. **c.** Heatmap showing representative tissue-specific top metabolites across selected tissue–cell-type combinations. Colour denotes the AUROC-derived effect size, highlighting organ-biased metabolite associations within specific cell types. **d.** Stacked bar plot showing representative levels–ratio paired associations across tissues. Colours indicate interpretation classes, including numerator accumulation, numerator depletion, denominator accumulation, denominator depletion and complex or imbalanced ratio patterns. **e.** Heatmap showing the number of positive significant metabolite associations across major cell types and tissues, revealing broad enrichment of metabolite-associated programmes in immune and blood-related populations as well as tissue-biased signals in endothelial, epithelial and stromal cells. **f.** Heatmap of positively conserved metabolites across cell types, with the right bar plot showing the number of unique conserved metabolites per cell type. **g.** Metabolic plasticity landscape of retained metabolite–cell-type pairs. The x axis represents tissue specificity, defined as one minus normalized entropy, and the y axis represents direction consistency across tissues. Points are coloured by plasticity class and sized by the number of tissues in which the pair was detected. Dashed lines indicate the thresholds used to classify broadly consistent, stable tissue-restricted, diffuse mixed and plastic tissue-restricted programmes. **h.** Bar plot showing the number of retained metabolite–cell-type pairs assigned to each direction-consistency class for each cell type. Together, these analyses distinguish conserved cross-tissue metabolite programmes from tissue-restricted and directionally plastic metabolic states.

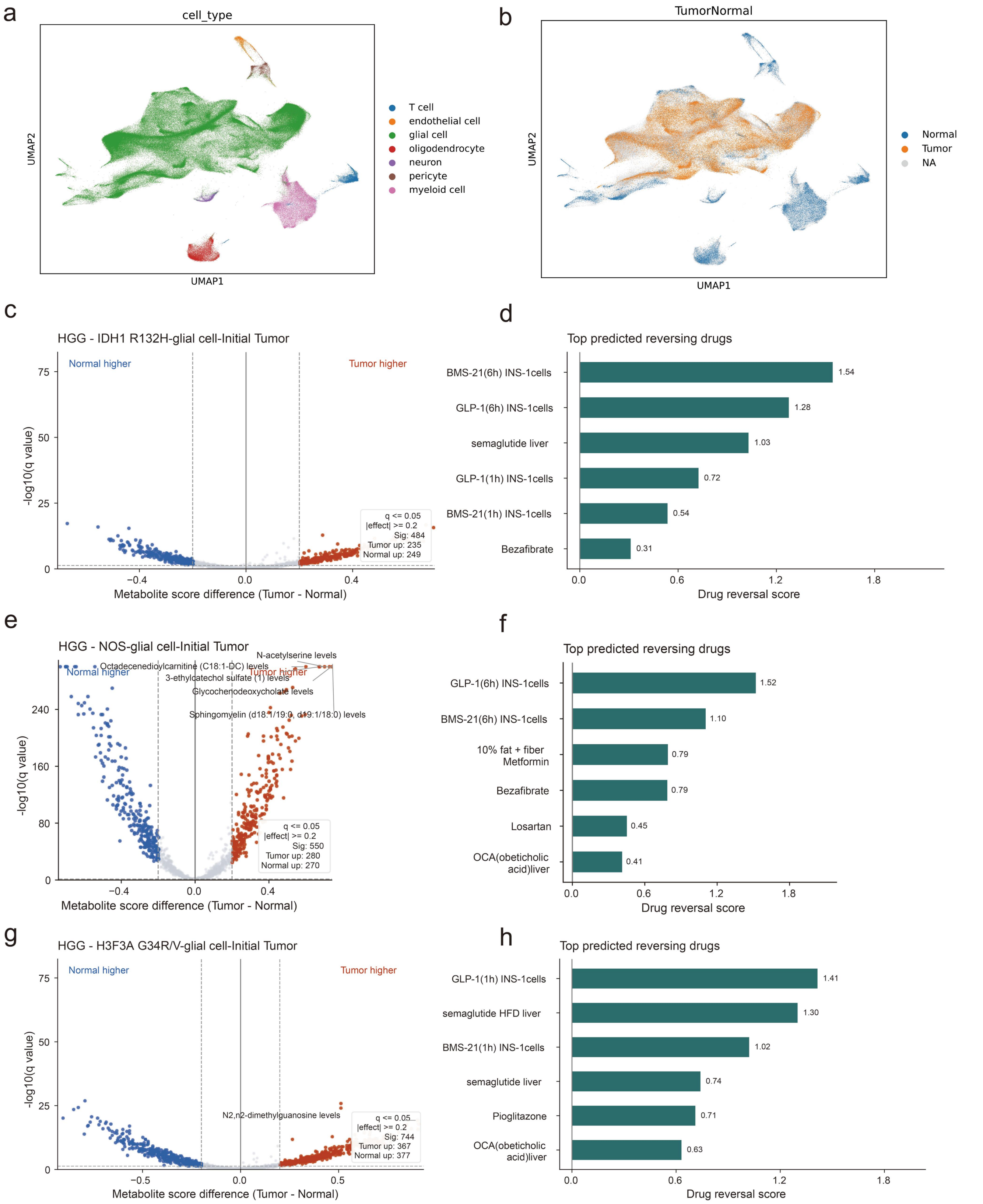


**Supplementary Figure S6. gmMAP identifies tumor-associated metabolic alterations and candidate metabolism-modulating drugs in pediatric high-grade glioma.**

**a,** UMAP visualization of the pediatric high-grade glioma single-cell atlas annotated by major cell types, including T cells, endothelial cells, glial cells, oligodendrocytes, neurons, pericytes and myeloid cells.

**b,** UMAP visualization of tumor and normal cell annotations across the same atlas. Tumor cells and normal cells occupy partially overlapping but distinct regions, supporting downstream comparison of tumor-associated metabolic programs.

**c,** Volcano plot showing differential gmMAP metabolite scores between tumor and normal glial cells in the IDH1 R132H pediatric high-grade glioma initial tumor sample. The x axis represents the metabolite score difference between tumor and normal cells, and the y axis represents −log10-transformed q values. Red points indicate tumor-enriched metabolite traits, blue points indicate normal-enriched metabolite traits and grey points indicate non-significant traits.

**d,** Top predicted reversing drugs for the IDH1 R132H tumor-associated metabolic signature identified by gmMAP-drug. Bars indicate drug reversal scores, with higher values representing stronger predicted reversal of tumor-enriched metabolic programs.

**e,** Volcano plot showing differential gmMAP metabolite scores between tumor and normal glial cells in NOS glial cell initial tumor samples. Representative tumor-enriched metabolites include N-acetylserine, glycochenodeoxycholate and phospholipid-related traits, whereas octadecanedioylcarnitine and 3-ethylcatechol sulfate are enriched in normal cells.

**f,** Top predicted reversing drugs for the NOS glial cell tumor-associated metabolic signature. GLP-1 perturbation, BMS-21, metformin-associated signatures, bezafibrate, losartan and obeticholic acid-associated profiles were among the top-ranked candidates.

**g,** Volcano plot showing differential gmMAP metabolite scores between tumor and normal glial cells in H3F3A G34R/V pediatric high-grade glioma initial tumor samples. N2,N2-dimethylguanosine was among the tumor-enriched metabolite traits.

**h,** Top predicted reversing drugs for the H3F3A G34R/V tumor-associated metabolic signature, including GLP-1 perturbation, semaglutide-associated liver signatures, BMS-21, pioglitazone and obeticholic acid-associated profiles

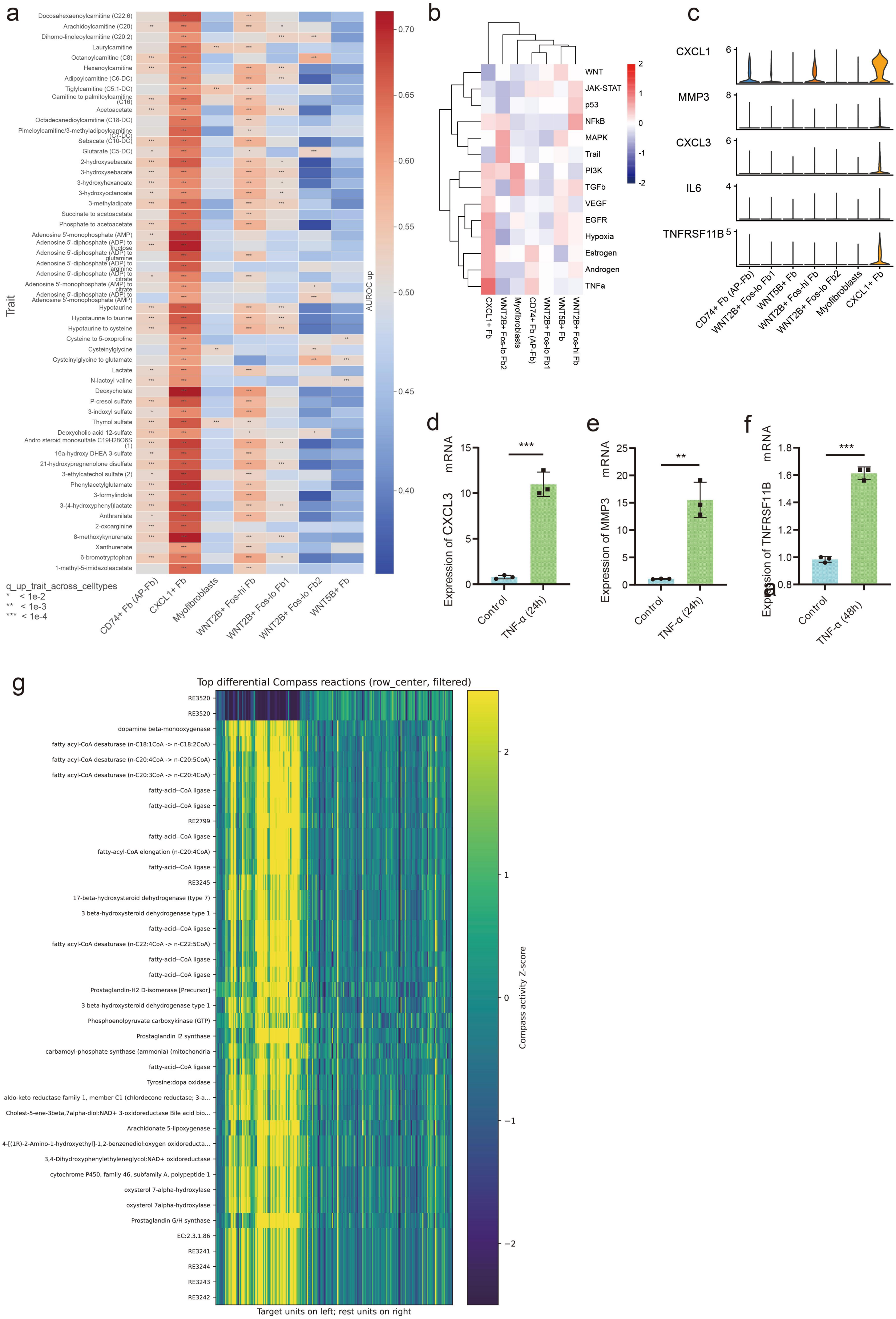
a
Docosahexaenoylcarnitine (C22:6)
Arachidoylcarnitine (C20)
Dihomo-linoleoylcarnitine (C20:2)
Laurylcarnitine
Octanoylcarnitine (C8)
Hexanoylcarnitine
Adipoylcarnitine (C6-DC)
Tiglylcarnitine (C5:1-DC)
Carnitine to palmitoylcarnitine (C16)
Acetoacetate
Octadecanedioylcarnitine (C18-DC)
Pimeloylcarnitine/3-methyladipoylcarnitine (C7-DC)
Sebacate (C10-DC)
Glutarate (C5-DC)
2-hydroxysebacate
3-hydroxysebacate
3-hydroxyhexanoate
3-hydroxyoctanoate
3-methyladipate
Succinate to acetoacetate
Phosphate to acetoacetate
Adenosine 5'-monophosphate (AMP)
Adenosine 5'-diphosphate (ADP) to fructose
Adenosine 5'-diphosphate (ADP) to glutamine
Adenosine 5'-diphosphate (ADP) to arginine
Adenosine 5'-diphosphate (ADP) to citrate
Adenosine 5'-monophosphate (AMP) to citrate
Adenosine 5'-diphosphate (ADP) to Adenosine 5'-monophosphate (AMP)
Hypotaurine
Hypotaurine to taurine
Hypotaurine to cysteine
Cysteine to 5-oxoproline
Cysteinylglycine
Cysteinylglycine to glutamate
Lactate
N-lactoyl valine
Deoxycholate
P-cresol sulfate
3-indoxyl sulfate
Thymol sulfate
Deoxycholic acid 12-sulfate
Andro steroid monosulfate C19H28O6S (1)
16a-hydroxy DHEA 3-sulfate
21-hydroxypregnenolone disulfate
3-ethylcatechol sulfate (2)
Phenylacetylglutamate
3-formylindole
3-(4-hydroxyphenyl)lactate
Anthranilate
2-oxoarginine
8-methoxykynurenate
Xanthurenate
6-bromotryptophan
1-methyl-5-imidazoleacetate
Trait
AUROC up
0.70
0.65
0.60
0.55
0.50
0.45
0.40
q_up_trait_across_celltypes
* < 1e-2
** < 1e-3
*** < 1e-4
CD74+ Fb (AP-Fb)
CXCL1+ Fb
Myofibroblasts
WNT2B+ Fos-hi Fb
WNT2B+ Fos-lo Fb1
WNT2B+ Fos-lo Fb2
WNT5B+ Fb
b
WNT
JAK-STAT
p53
NFkB
MAPK
Trail
PI3K
TGFb
VEGF
EGFR
Hypoxia
Estrogen
Androgen
TNFa
2
1
0
-1
-2
CXCL1+ Fb
WNT2B+ Fos-lo Fb2
Myofibroblasts
CD74+ Fb (AP-Fb)
WNT2B+ Fos-lo Fb1
WNT5B+ Fb
WNT2B+ Fos-hi Fb
c
CXCL1
MMP3
CXCL3
IL6
TNFRSF11B
CD74+ Fb (AP-Fb)
WNT2B+ Fos-lo Fb1
WNT5B+ Fb
WNT2B+ Fos-hi Fb
WNT2B+ Fos-lo Fb2
Myofibroblasts
CXCL1+ Fb
d
Expression of CXCL3 mRNA
***
Control
TNF-α (24h)
e
Expression of MMP3 mRNA
**
Control
TNF-α (24h)
f
Expression of TNFRSF11B mRNA
***
Control
TNF-α (48h)
g
Top differential Compass reactions (row_center, filtered)
RE3520
RE3520
dopamine beta-monooxygenase
fatty acyl-CoA desaturase (n-C18:1CoA -> n-C18:2CoA)
fatty acyl-CoA desaturase (n-C20:4CoA -> n-C20:5CoA)
fatty acyl-CoA desaturase (n-C20:3CoA -> n-C20:4CoA)
fatty-acid--CoA ligase
fatty-acid--CoA ligase
RE2799
fatty-acid--CoA ligase
fatty-acyl-CoA elongation (n-C20:4CoA)
fatty-acid--CoA ligase
RE3245
17-beta-hydroxysteroid dehydrogenase (type 7)
3 beta-hydroxysteroid dehydrogenase type 1
fatty-acid--CoA ligase
fatty acyl-CoA desaturase (n-C22:4CoA -> n-C22:5CoA)
fatty-acid--CoA ligase
fatty-acid--CoA ligase
Prostaglandin-H2 D-isomerase [Precursor]
3 beta-hydroxysteroid dehydrogenase type 1
Phosphoenolpyruvate carboxykinase (GTP)
Prostaglandin I2 synthase
carbamoyl-phosphate synthase (ammonia) (mitochondria
fatty-acid--CoA ligase
Tyrosine:dopa oxidase
aldo-keto reductase family 1, member C1 (chlordecone reductase; 3-a...
Cholest-5-ene-3beta,7alpha-diol:NAD+ 3-oxidoreductase Bile acid bio...
Arachidonate 5-lipoxygenase
4-[(1R)-2-Amino-1-hydroxyethyl]-1,2-benzenediol:oxygen oxidoreducta...
3,4-Dihydroxyphenylethyleneglycol:NAD+ oxidoreductase
cytochrome P450, family 46, subfamily A, polypeptide 1
oxysterol 7-alpha-hydroxylase
oxysterol 7alpha-hydroxylase
Prostaglandin G/H synthase
EC:2.3.1.86
RE3241
RE3244
RE3243
RE3242
Compass activity Z-score
Target units on left; rest units on right

**Supplementary Figure S7. CXCL1⁺ inflammatory fibroblasts exhibit lactate-associated metabolic remodeling and TNFα-driven transcriptional activation.**

**a.** Heatmap showing gmMAP-predicted metabolite-associated programs across fibroblast subpopulations. Color intensity indicates AUROC enrichment scores for positive metabolite-associated programs. $CXCL1^{+}$ fibroblasts display the strongest enrichment of lactate-associated metabolic signatures together with multiple energetic-stress-associated metabolite traits.

**b.** decoupleR-based pathway activity inference across fibroblast subpopulations. $CXCL1^{+}$ fibroblasts exhibit the highest activity of TNFα-associated transcriptional programs.

**c.** Violin plots showing expression of representative inflammatory mediators and TNF-responsive genes across fibroblast states. $CXCL1^{+}$ fibroblasts preferentially express CXCL1, CXCL3, MMP3 and TNFRSF11B compared with other fibroblast populations.

**d–f.** qPCR validation of TNFα-responsive inflammatory programs in primary human colonic fibroblasts. TNFα stimulation significantly induces expression of CXCL3 (d), MMP3 (e) and TNFRSF11B (f), supporting activation of a TNFα-dominated inflammatory transcriptional state. Data are presented as mean ± s.d. Statistical significance was assessed using two-sided Student's *t*-test. P values are indicated in the figure ($P < 0.01$, $P < 0.001$).

**g.** Metabolic flux signatures upregulated in $CXCL1^{+}$ fibroblasts predicted by COMPASS.

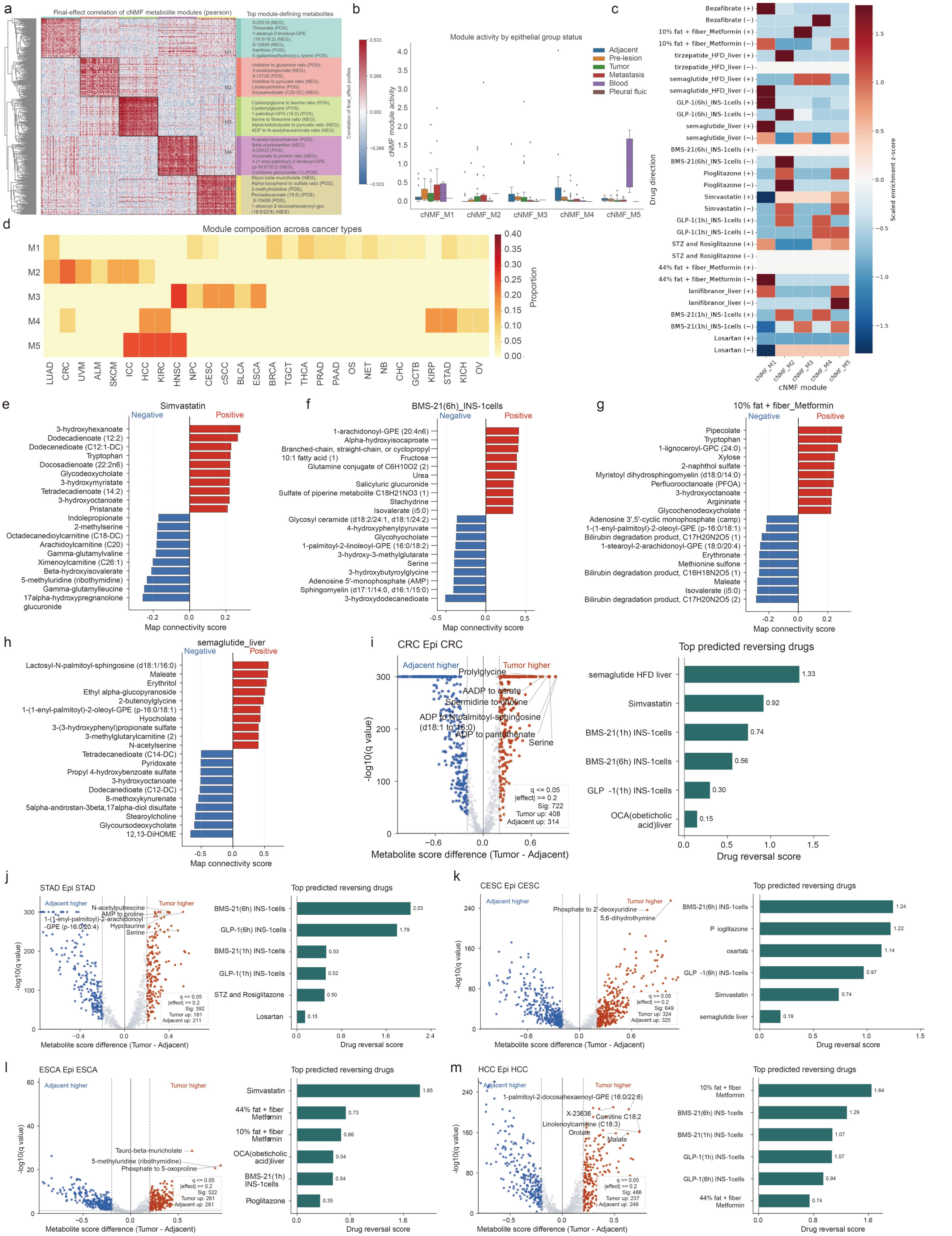

**Supplementary Figure S8. gmMAP-drug identifies metabolism-modulating drugs predicted to reverse pan-cancer epithelial metabolic programmes.**

**a.** Consensus NMF decomposition of the pan-cancer epithelial metabolite final-effect matrix. Left, correlation heatmap of cNMF-derived metabolite modules based on final-effect profiles. Right, representative top module-defining metabolites for each epithelial metabolic module. **b.** Module activity across epithelial disease states, including adjacent, pre-lesion, tumor, metastatic, blood and pleural-fluid epithelial groups. Points represent epithelial cancer-state groups or samples, and bars show module activity summaries. **c.** Row z-scored enrichment heatmap of the top 100 drug-connected metabolites across cNMF modules and selected gmMAP-drug perturbations. Drug perturbations are annotated by predicted direction, and colour indicates scaled enrichment z-score. **d.** Module composition across cancer types, showing the proportional contribution of each cNMF module to epithelial metabolic programmes in individual cancers. **e–h**. Bilateral drug–metabolite connectivity plots for representative metabolism-modulating perturbations, including simvastatin, BMS-21, metformin-associated and semaglutide-associated signatures. Red bars indicate positively connected metabolites and blue bars indicate negatively connected metabolites. **i–m**. Tumor-versus-adjacent differential metabolite activity and predicted drug reversal analysis for representative epithelial cancer contexts, including CRC, STAD, CESC, ESCA and HCC. Volcano plots show metabolite score differences between tumor and adjacent epithelial cells; red points denote tumor-enriched metabolites and blue points denote adjacent-enriched metabolites. Bar plots show the top predicted reversing drugs ranked by gmMAP-drug reversal score. Together, these analyses show that pan-cancer epithelial metabolic states can be decomposed into recurrent metabolite programmes and pharmacologically queried by gmMAP-drug to nominate metabolism-modulating drugs with predicted reversal potential.

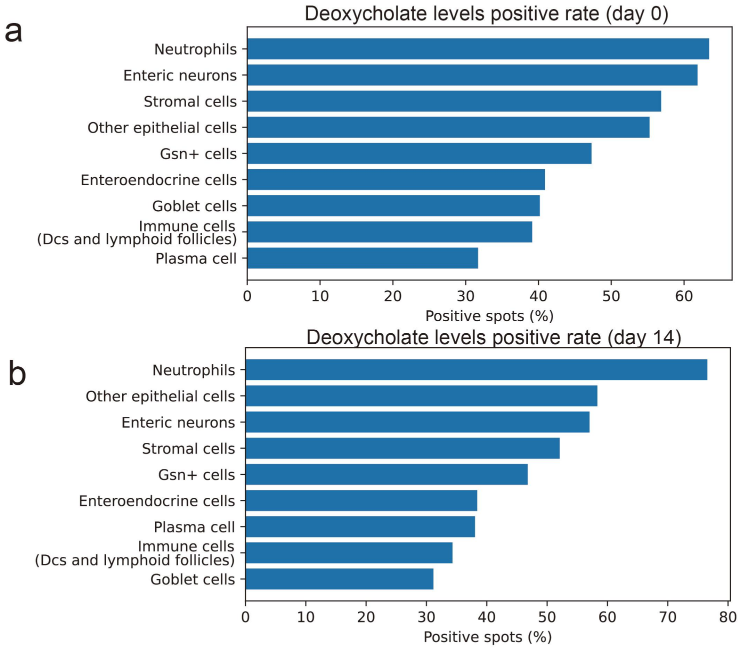


**Supplementary Figure S9. Cell-type-specific distribution of deoxycholate-associated programmes during colitis progression and recovery.**

**a.** Percentage of deoxycholate-positive spatial spots within each annotated cell population in healthy control tissue (Day 0). **b.** Percentage of deoxycholate-positive spatial spots within each annotated cell population during the recovery phase following DSS-induced colitis (Day 14). Positive spots were defined as spatial locations with deoxycholate-associated scores exceeding the predefined enrichment threshold. Neutrophils, epithelial populations and stromal cells exhibited the highest proportions of deoxycholate-positive spots in both conditions, with further enrichment observed during recovery. These results indicate persistent cell-type-specific exposure to deoxycholate-associated metabolic programmes throughout both the acute inflammatory and post-inflammatory repair phases of colitis.

### Methods

#### gmMAP-flux: Enzyme-constrained inference of single cell metabolic flow potential

We developed gmMAP-flux, an enzyme-constrained metabolic-flow potential model designed to infer directional metabolic activity across kidney developmental cell states. gmMAP-flux integrates metabolite-linked wAUCell-S3 scores, metabolite-ratio directionality, enzyme expression, subcellular-compartment support and cofactor-associated transcriptional support. The model is not a stoichiometric flux-balance analysis framework, but rather a transcriptome-constrained flux-potential model that estimates the relative direction and feasibility of metabolite-linked reaction-like modules across cell states.

The input consisted of three major data layers: a curated metabolite-reaction module table, a wAUCell-S3 metabolite score matrix and a matched gene-expression matrix. The module table contained module identifiers, input metabolite traits, output metabolite traits, optional ratio traits, curated ratio direction signs, pathway and subpathway annotations, and optional enzyme-gene annotations. The wAUCell-S3 matrix was defined with cells or cell types as rows and metabolite traits as columns. The gene-expression matrix was defined with matched rows and gene symbols as columns. When cell-level matrices were provided, both metabolite scores and gene-expression values were aggregated to cell types using the mean or median expression according to the supplied cell metadata.

##### Notation and parameter definitions

To facilitate interpretation of the mathematical definitions below, we summarize the main symbols and their corresponding implementation variables. Scale parameters are hyperparameters used only to normalize values before tanh transformation; all scale parameters were set to 1.0 unless otherwise specified.

**Supplementary table S2**

| Symbol | Code variable / output column | Definition |
|---|---|---|
| m | module_id | Metabolite-reaction module. |
| c | celltype | Cell type or aggregated cell state. |
| s^in_{m,c} | trait_in score | wAUCell-S3 score of the input metabolite trait in cell type c. |
| s^out_{m,c} | trait_out score | wAUCell-S3 score of the output metabolite trait in cell type c. |
| s^ratio_{m,c} | ratio_trait score | wAUCell-S3 score of the matched ratio trait when available. |
| d_m | direction_sign | Curated ratio direction sign; defaults to +1 if not provided. |
| w_r | ratio_weight | Weight assigned to ratio-derived directional support. |
| Delta s_{m,c} | delta_s3 | Output-input metabolite score difference. |
| A_{m,c} | activation | Bounded module activation based on absolute input/output metabolite-node intensity. |
| D_{m,c} | direction | Bounded directional score integrating endpoint-derived and ratio-derived evidence. |
| C_m | coverage | Input/output trait coverage. |
| N_{m,c} | node_support | Support from input and output metabolite-node signals. |
| R_{m,c} | ratio_factor | Ratio-strength factor or missing-ratio/no-ratio factor. |
| K_{m,c} | consistency | Directional concordance between endpoint-derived and |

| | | |
|---|---|---|
| | | ratio-derived signs. |
| E_{m,c} | enzyme_capacity | Enzyme expression support combined with enzyme-gene coverage. |
| P_{m,c} | compartment_support | Support from compartment-associated hallmark gene panels. |
| Q_{m,c} | cofactor_support | Support from cofactor- or pathway-associated hallmark gene panels. |
| F_{m,c} | feasibility / confidence | Final feasibility score used as confidence. |
| Phi_{m,c} | signed_flux | Final signed metabolic-flow potential. |
| Y_{p,c} | subpathway metric | Subpathway-level metric obtained by feasibility-weighted module aggregation. |

**Default parameters used in the implementation**

**Supplementary table S3**

| Parameter | Code argument | Default value | Role |
|---|---|---|---|
| lambda_A | activation_scale | 1.0 | Scale for module activation before tanh transformation. |
| lambda_D | direction_scale | 1.0 | Scale for directional evidence before tanh transformation. |
| lambda_S | signal_scale | 1.0 | Scale for metabolite-node support. |
| lambda_R | ratio_scale | 1.0 | Scale for ratio-strength factor. |
| lambda_E | enzyme_scale | 1.0 | Scale for enzyme-gene expression support. |
| lambda_P | compartment_scale | 1.0 | Scale for compartment hallmark gene support. |
| lambda_Q | cofactor_scale | 1.0 | Scale for cofactor hallmark gene support. |
| alpha_E | enzyme_weight | 1.0 | Exponent controlling enzyme-capacity contribution. |
| alpha_P | compartment_weight | 0.5 | Exponent controlling compartment-support contribution. |
| alpha_Q | cofactor_weight | 0.25 | Exponent controlling cofactor-support contribution. |
| p_missing-ratio | ratio_missing_penalty | 0.5 | Penalty when a ratio trait is assigned but absent. |
| p_no-ratio | no_ratio_factor | 1.0 | No penalty for modules without ratio annotation. |
| p_no-enzyme | no_enzyme_factor | 0.8 | Default support for modules without enzyme-gene annotation. |
| p_no-compartment | no_compartment_factor | 0.9 | Default support when no compartment panel is assigned. |
| p_no-cofactor | no_cofactor_factor | 0.9 | Default support when no cofactor panel is assigned. |
| k | gpr_topk | 3 | Number of top expressed genes used for the default top-k mean gene-set score. |

For each metabolic module $m$ in cell type $c$, we extracted the S3-normalized input metabolite score, output metabolite score and, when available, ratio-trait score:

$$s_{mc}^{in}, \quad s_{mc}^{out}, \quad s_{mc}^{ratio}.$$

The average metabolite-node signal and the endpoint-derived directional difference were defined as:

$$\bar{s}_{mc} = \text{mean}(s_{mc}^{in}, s_{mc}^{out}),$$

$$\Delta s_{mc} = s_{mc}^{out} - s_{mc}^{in}.$$

Module activation was defined as a bounded function of the absolute input and output metabolite-node intensities:

$$A_{mc} = \tanh\left(\frac{\mathrm{mean}\left(|s_{mc}^{in}|, |s_{mc}^{out}|\right)}{\lambda_A}\right),$$

where $\lambda_A$ is the activation-scale parameter. This term represents whether the metabolite nodes connected by a reaction-like module are transcriptionally supported in a given cell type, independent of the sign of directionality.

When a matched metabolite-ratio trait was available, ratio-derived directional support was calculated as:

$$R_{mc}^{dir} = d_m s_{mc}^{ratio},$$

where $d_m \in \{-1, +1\}$ is the curated ratio direction sign. If the ratio direction sign was unavailable, $d_m$ was set to $+1$. The final direction input integrated endpoint-derived and ratio-derived evidence:

$$I_{mc}^{dir} = \Delta s_{mc} + w_r R_{mc}^{dir},$$

where $w_r$ is the ratio-weight parameter. If no ratio score was available, the direction input was defined only by the input-output difference:

$$I_{mc}^{dir} = \Delta s_{mc}.$$

The bounded module direction was then computed as:

$$D_{mc} = \tanh\left(\frac{I_{mc}^{dir}}{\lambda_D}\right),$$

where $\lambda_D$ is the direction-scale parameter. Positive values of $D_{mc}$ indicate output-dominant or forward-supported metabolic-flow potential, whereas negative values indicate input-dominant or reverse-supported metabolic-flow potential.

To quantify whether the metabolite nodes themselves were sufficiently supported, node support was calculated as:

$$N_{mc} = \mathrm{mean}\left[\tanh\left(\frac{|s_{mc}^{in}|}{\lambda_S}\right), \tanh\left(\frac{|s_{mc}^{out}|}{\lambda_S}\right)\right],$$

where $\lambda_S$ is the metabolite-signal scale. Trait coverage was defined as:

$$C_m = \frac{\mathbf{1}\left(s_m^{in}\ \text{available}\right) + \mathbf{1}\left(s_m^{out}\ \text{available}\right)}{2}.$$

Thus, modules with both input and output traits present received full coverage support, whereas modules with only one available endpoint were down-weighted.

For modules with an assigned ratio trait, the ratio-strength factor was defined as:

$$R_{mc} = \tanh\left(\frac{|s_{mc}^{ratio}|}{\lambda_R}\right),$$

where $\lambda_R$ is the ratio-scale parameter. If a ratio trait was assigned but missing from the score matrix, the module received a missing-ratio penalty:

$$R_{mc} = p_{\text{missing-ratio}}.$$

Modules without ratio annotation were not penalized and were assigned:

$$R_{mc} = p_{\text{no-ratio}}.$$

Directional consistency between endpoint-derived direction and ratio-derived direction was encoded as a soft factor:

$$K_{mc} = \begin{cases} 1, & \text{if no ratio-derived direction is available,} \\ 0.5, & \text{if } |\Delta s_{mc}| \approx 0 \text{ or } \left|R_{mc}^{dir}\right| \approx 0, \\ 1, & \text{if } \operatorname{sign}(\Delta s_{mc}) = \operatorname{sign}\left(R_{mc}^{dir}\right), \\ 0.25, & \text{if } \operatorname{sign}(\Delta s_{mc}) \neq \operatorname{sign}\left(R_{mc}^{dir}\right). \end{cases}$$

Thus, concordant endpoint- and ratio-derived directions received full support, whereas discordant signs were down-weighted.

Enzyme capacity was computed from module-specific enzyme genes using a GPR-like proxy. Gene symbols were normalized, intersected with the expression matrix and scored using a bounded gene-set expression function. For a module-specific enzyme-gene set $G_m$, available genes in cell type $c$ were denoted as $G_m^{avail}$. By default, enzyme-gene expression support was calculated using the top-$k$ mean expression:

$$X_{mc}^{enzyme} = \frac{1}{k} \sum_{g \in \operatorname{TopK}\left(G_m^{avail}\right)} x_{gc},$$

where $x_{gc}$ is the expression of gene $g$ in cell type $c$. The bounded enzyme-expression score was:

$$S_{mc}^{enzyme} = \tanh\left(\frac{X_{mc}^{enzyme}}{\lambda_E}\right),$$

where $\lambda_E$ is the enzyme-expression scale. Enzyme-gene coverage was defined as:

$$V_m^{enzyme} = \frac{\left|G_m^{avail}\right|}{|G_m|}.$$

For modules with annotated enzyme genes, enzyme capacity was calculated as:

$$E_{mc} = 0.5 S_{mc}^{enzyme} + 0.5 V_m^{enzyme}.$$

For modules without enzyme-gene annotation, a default no-enzyme factor was used:

$$E_{mc} = p_{\text{no-enzyme}}.$$

Compartment support and cofactor support were inferred from curated hallmark gene panels. Based on pathway class, subpathway annotation and metabolite names, each module was matched to one or more compartment-associated panels, including mitochondrial, peroxisomal, glycolytic/cytosolic, urea-cycle, tryptophan-kynurenine and transport-related panels. For a compartment panel $P_j$, the bounded panel score was calculated using the same top-$k$ gene-set scoring strategy:

$$S_{jmc}^{comp} = \tanh\left(\frac{\operatorname{TopKMean}\left(x_{gc}: g \in P_j \cap X_c\right)}{\lambda_P}\right),$$

where $\lambda_P$ is the compartment-support scale. If multiple compartment panels were assigned, compartment support was computed as the mean of valid panel scores:

$$P_{mc} = \text{mean}_j \left( S_{jmc}^{comp} \right).$$

If no compartment panel was assigned, a default no-compartment factor was used:

$$P_{mc} = p_{\text{no-compartment}}.$$

Similarly, cofactor-associated support was inferred from pathway-specific cofactor or functional-support panels. These included fatty-acid oxidation/TCA-associated genes, glycolysis/pyruvate-associated genes, urea-cycle genes and tryptophan-kynurenine-associated genes. For a cofactor panel $Q_m$, cofactor support was defined as:

$$Q_{mc} = \tanh\left( \frac{\text{TopKMean}\left(x_{gc}: g \in Q_m \cap X_c\right)}{\lambda_Q} \right),$$

where $\lambda_Q$ is the cofactor-support scale. If no cofactor panel was assigned, a default no-cofactor factor was used:

$$Q_{mc} = p_{\text{no-cofactor}}.$$

The final feasibility score, used as the confidence of each module-cell-type pair, was computed as the product of metabolite-node support, trait coverage, ratio support, directional consistency and the three transcriptome-derived soft constraints:

$$F_{mc} = C_m N_{mc} R_{mc} K_{mc} E_{mc}^{\alpha_E} P_{mc}^{\alpha_P} Q_{mc}^{\alpha_Q},$$

where $E_{mc}$, $P_{mc}$ and $Q_{mc}$ denote enzyme capacity, compartment support and cofactor support, respectively. The exponents $\alpha_E$, $\alpha_P$ and $\alpha_Q$ control the relative contributions of enzyme, compartment and cofactor constraints. In the default implementation, enzyme capacity was given the largest contribution, followed by compartment support and cofactor support:

$$\alpha_E = 1.0, \quad \alpha_P = 0.5, \quad \alpha_Q = 0.25.$$

The final signed metabolic-flow potential was then calculated as:

$$\Phi_{mc} = A_{mc} D_{mc} F_{mc}.$$

Here, $A_{mc}$ represents module activation, $D_{mc}$ represents directionality and $F_{mc}$ represents transcriptome-constrained feasibility. Therefore, $\Phi_{mc} > 0$ indicates forward or output-dominant metabolic-flow potential, whereas $\Phi_{mc} < 0$ indicates reverse or input-dominant metabolic-flow potential. The magnitude $|\Phi_{mc}|$ reflects the relative strength of the inferred flow potential. Because the model does not use stoichiometric reaction matrices or mass-balance constraints, $\Phi_{mc}$ should be interpreted as a relative, transcriptome-constrained metabolic-flow potential rather than an absolute biochemical flux.

For subpathway-level inference, module-level scores were aggregated within each subpathway using feasibility/confidence as weights. For a score $Y_{mc}$, such as activation, direction, signed flux, ratio support, metabolite-node signal, input-output difference, enzyme capacity, compartment support or cofactor support, the subpathway-level score for subpathway $p$ in cell type $c$ was calculated as:

$$Y_{pc} = \frac{\sum_{m \in p} F_{mc} Y_{mc}}{\sum_{m \in p} F_{mc}}.$$

Subpathway-level confidence was calculated as the mean module feasibility within the subpathway:

$$F_{pc} = \text{mean}_{m \in p}(F_{mc}).$$

The model exported module-level and subpathway-level matrices for activation, direction, confidence, signed flux, ratio support, enzyme capacity, compartment support, cofactor support, feasibility, mean S3 score and input-output delta score. Diagnostic outputs were also generated to record missing metabolite traits, ratio-trait availability, available enzyme genes and constraint-gene usage for each module.

### Genetically informed metabolite scoring for single-cell, spatial and bulk transcriptomic data

$$\text{wAUCell}_{c,t} = \frac{\sum_{r=1}^{K} \mathbf{1}\left(g_{c,r} \in G_t\right) \cdot w_{g_{c,r}} \cdot \rho_r}{\sum_{g \in G_t} w_g}$$

This table defines each symbol used in the weighted AUCell-like score for genetically informed metabolite-trait mapping at the single-cell, spatial spot, or sample level.

**Supplementary table S4**

| Symbol | Meaning | Interpretation / Notes |
|---|---|---|
| wAUCell(c,t) | Weighted AUCell-like score for observation c and metabolite trait t. | A normalized rank-based enrichment score measuring whether genes associated with metabolite trait t are highly ranked in observation c. |
| c | Index of an observation. | Usually denotes a single cell or spatial spot. In bulk RNA-seq applications, c can also represent a sample. |
| t | Index of a metabolite trait. | A metabolite trait can be a metabolite level, metabolite ratio, or another GWAS-derived metabolite-associated feature. |
| G_t | Gene set associated with metabolite trait t. | In practice, this can be the up gene set G_t^up or the down gene set G_t^down derived from MAGMA gene-level Z-scores. |
| K | Number of top-ranked genes retained in each observation. | In the implemented workflow, K is defined as the top 5% of all genes ranked by expression within each observation: K = floor(0.05 × G). |
| G | Total number of genes considered in the expression matrix. | Used to determine K, the number of genes retained for rank-based enrichment. |
| r | Rank position within observation c. | Genes are ordered from high to low expression within each observation; r = 1 corresponds to the highest-ranked gene. |
| g(c,r) | Gene ranked at position r in observation c. | This is the gene encountered at rank r among the top-K genes of observation c. |
| 1(g(c,r) ∈ G_t) | Indicator function. | Equals 1 if the ranked gene g(c,r) belongs to the trait-associated gene set G_t; otherwise equals 0. This restricts the summation to genes in G_t. |
| w[g(c,r)] | Weight of gene g(c,r). | A genetically informed gene weight assigned to the gene ranked at position r in observation c. |
| ρ_r | Rank-position weight. | A decreasing positional weight that gives greater contribution to higher-ranked genes. In the implemented workflow, ρ_r = (K − r + 1) / K. |
| Σ_{r=1}^{K} | Summation over the top-K ranked genes in observation c. | Only genes among the top-K expression ranks are evaluated in the numerator. |
| Σ_{g∈G_t} w_g | Total weight of all genes in trait gene set G_t. | Used as the denominator to normalize the enrichment score across traits with different total gene weights. |

| Symbol | Meaning | Interpretation / Notes |
|---|---|---|
| w_g | Genetically informed weight for gene g. | Typically defined as w_g = z_g / σ_g, so genes with stronger genetic association and lower expression variability receive larger weights. |
| z_g | Gene-level genetic association score for gene g. | Usually a MAGMA-derived gene-level Z-score for the metabolite trait. For down signatures, the absolute magnitude of negative Z-scores is used. |
| σ_g | Gene-specific expression variability or noise term. | Estimated from the variance of gene expression across all observations: σ_g = sqrt(Var(X_g)). |
| X_g | Expression vector of gene g across observations. | Used to estimate gene-specific expression variability. |
| X(c,g) | Expression of gene g in observation c. | Although not explicitly shown in the wAUCell formula, it determines the within-observation expression rank of each gene. |

Practical interpretation: the numerator sums the weights of trait-associated genes that appear among the top-ranked genes of an observation, while also accounting for their rank positions. The denominator normalizes this value by the total genetic weight of the trait gene set.

$$\mathrm{sMRS}_{s,t} = \frac{\sum_{g \in G_t} X_{s,g}\, w_{g,t}}{\sum_{g \in G_t} |w_{g,t}|} \, ,$$

The formula defines a normalized weighted-sum metabolite relevance score for a sample or expression profile. The symbols are explained below.

**Supplementary table S5**

| Symbol | Explanation |
|---|---|
| sMRS_{s,t} | The sample-level Metabolite Relevance Score for sample or expression profile s and metabolite trait t. |
| s | Index of the sample or expression profile. In bulk RNA-seq, s denotes an individual bulk sample; in a generalized matrix, it may also denote any observation-level profile. |
| t | Index of the metabolite trait, such as a metabolite level or metabolite ratio. |
| G_t | The metabolite trait-associated gene set for trait t. In practice, this can be the up-associated gene set G_t^up or the down-associated gene set G_t^down. |
| g | Index of a gene included in the trait-associated gene set G_t. |
| X_{s,g} | The expression value of gene g in sample or expression profile s after the selected expression preprocessing procedure. |
| w_{g,t} | The genetically informed weight assigned to gene g for metabolite trait t. It reflects the MAGMA/GWAS-derived association strength and may be scaled by gene-level expression variability. |
| \|w_{g,t}\| | The absolute value of the gene weight. In the denominator, it ensures that the score is normalized by the total magnitude of the trait-specific weights. |
| Σ_{g∈G_t} | Summation over all genes g belonging to the metabolite trait-associated gene set G_t. |
| Numerator | Σ_{g∈G_t} X_{s,g} w_{g,t}; the weighted expression signal of all trait-associated genes in sample s. |
| Denominator | Σ_{g∈G_t} \|w_{g,t}\|; the total absolute weight of the trait-associated gene set, used to normalize the score across traits with different weight magnitudes. |

**Interpretation.** sMRS summarizes whether genes genetically associated with metabolite trait t are highly expressed in sample s. The numerator aggregates genetically weighted expression, whereas the denominator rescales the score by the total absolute weight of the gene set.

**Optional weight definition.** When gene-noise scaling is used, the weight can be written as $w_{g,t}=z^*_{g,t}/\sigma_g$, where $z^*_{g,t}$ is the positive directional gene-level association magnitude for trait t and $\sigma_g$ is the expression variability of gene g across samples or observations.

### Information of quantitative real-time PCR (qPCR) primer sequences

**Supplementary table S6.** qPCR primer sequences

| Gene | Gene ID | **Forward primer (5′→3′)** | **Reverse primer (5′→3′)** |
|---|---|---|---|
| Cxcl1 | 14825 | CCCAAACCGAAGTCATAGCC | CAGGTGCCATCAGAGCAGTC |
| Tnfrsf11b | 18383 | ACGGAGACACAGCTCACAAG | GATCTTCTTCCCAGGCAGGC |
| Mmp3 | 17392 | CATGGAGCCAGGATTTCCCA | TGGGTCAAATTCCAACTGCG |
| HIf1a | 15251 | GGACGATGAACATCAAGTCAGCA | GGACGATGAACATCAAGTCAGCA |
| Cxcl3 | 330122 | CCCAGACAGAAGTCATAGCC | CGTTGGGATGGATCGCTTTTC |
